# DNA and RNA from the same single nucleus reveals interactions between genomic and transcriptomic landscapes in human tumor samples

**DOI:** 10.1101/2023.10.04.560973

**Authors:** Siran Li, Joan Alexander, Jude Kendall, Peter Andrews, Elizabeth Rose, Hope Orjuela, Sarah Park, Craig Podszus, Liam Shanley, Nissim Ranade, Patrick Morris, Danielle Stauder, Daniel Bradford, Michael Ronemus, Arvind Rishi, Rong Ma, David L. Donoho, Gary L. Goldberg, Michael Wigler, Dan Levy

## Abstract

To deepen our understanding of cancer heterogeneity and uncover the dynamic interactions of tumor and host cells, we introduce hybrid BAG-seq: a high-throughput, multi-omic method that simultaneously captures DNA and RNA from tens of thousands of individual single nuclei. This method provides dual molecular layer information: DNA to distinguish tumor from stroma, identify tumor subclones, and detect mutant stromal sub-populations; and RNA to characterize distinct cell types, cell states, and signatures of aberrant expression. Additionally, we developed a suite of analysis tools to illuminate cluster phylogeny and connections between DNA identity and RNA expression. We applied this hybrid protocol to 65,499 single nuclei from samples of five uterine cancer patients, and validated the clustering using RNA-only and DNA-only protocols on 34,651 and 21,432 nuclei, respectively, from the same tissues. Multiple tumor genome or expression clusters were often present within a patient, with different tumor clones projecting into distinct or shared expression states, demonstrating nearly all possible genome-transcriptome correlations across the cohort. While tumor expression profiles were highly unique to each patient, the stromal cell types generally recurred across samples, but certain patients and tissues exhibited unique stromal sub-types characterized by aberrant expression. Moreover, we identified mutant stroma in various cell types from several patients with a significant loss of the X-chromosome. This study reveals the complex landscape of genome and transcriptome interactions at the resolution of single nuclei, providing new insights into mutant stroma and tumor heterogeneity.

## Introduction

Single-cell analysis offers an opportunity for insight into cancers and their interactions with the host. From single-cell RNA data, we can obtain valuable information about the cell type and state. This is key in characterizing the various types of normal stroma as well as the diverse expression states within the tumor. From single-cell DNA data, we can determine important lineage information. This enables us to distinguish tumor cells from normal cells and partition tumor cells into sub-clones. The fusion of these two types of analysis, high-throughput data of RNA and DNA from the same single cells, presents a significant advancement. By assigning both a distinct DNA and RNA identity to each single cell, we may begin to observe the complex interplay between cancer cells and stroma, the emergence of malignant cells from pre-malignant ones, the driving forces behind specific expression states, and possible patterns of mutation in certain types of host stroma.

Methods for the high-throughput capture of either single-cell DNA^1–3^ or RNA^4–11^ are well-established and widely utilized in current research. However, when it comes to investigating both omics simultaneously from single cells, existing techniques present certain limitations. There are low-throughput methods capable of capturing both nucleic acids from single cells^12–24^, yet these may not be appropriate for large-scale studies. Additionally, approaches for inferring copy number from high throughput RNA-alone data have been described^25–29^, but these methods rely on the assumption that gene expression and copy number are reliably correlated. The integration of high-throughput genome and transcriptome analysis in a single method has been previously described^30–32^, none have been applied to human tumor samples to systematically investigate the cancer heterogeneity and stromal mutations with the throughput demonstrated in this study. A method, proven to be robust and reliable in this context, could mark a significant leap forward in our understanding of cancer biology at the single-cell level.

In this study, we introduce, characterize, and implement a hybrid high-throughput droplet-based technology that enables the capture of both DNA and RNA templates from the same cell nucleus. This multi-omic technology is an evolution of our **BAG** platform^33^, where single cells are encapsulated into individual **b**alls of **a**crylamide **g**el, with nucleic acid templates captured by Acrydite primers and copolymerized into the gel matrix. We employ a pool-and-split method to assign unique cell-barcodes and varietal tags to each template. After sequencing, the nucleic acid reads from each cell are partitioned into distinct **DNA and RNA layers** based on the characteristics of the mapped read.

We utilized this **hybrid BAG** approach on the frozen tissue obtained from five patients diagnosed with uterine cancer. We found sufficient transcriptional complexity in the nuclear RNA to clearly distinguish cell types. The DNA layer provided sufficient genome copy number information to differentiate stroma from tumor, to identify distinct subclones within the tumor, and in some cases to identify mutant stroma. We further confirmed these clustering patterns through comparison with results obtained from established RNA-only and DNA-only BAG protocols.

Clustering in single-cell analysis presents a unique challenge. Popular methods like tSNE^34^ and UMAP^35,36^ provide compelling visualizations; however, underlying algorithmic constraints force cells into a pre-determined number of clusters. To quantify the likelihood of each cell’s cluster affiliation, we enhanced our analysis by modeling clusters with multinomial distributions^37^. Generalizing to a space of pairwise-linear combinations, we identify and remove most doublet collisions – instances of mistaken cellular identity that arise when two distinct cells are assigned the same identity. We also use multinomial distributions to measure an inter-cluster distance: two clusters are “far apart” if very few unique templates are required to determine that a random cell from one cluster is unlikely to have originated from the other.

The tumor and host expression clusters from each patient exhibited distinct and unique characteristics. We use the gene count vectors for the clusters to determine differential gene expression sets that inform labels of cell type and state. Combining expression data from all five patients’ samples, we observed that the profiles of the somatic cell types are broadly consistent across all patients, but with some interesting differences.

Employing high-quality, high-throughput single-cell RNA and DNA data, we explore the complex genomic and expression landscape of five uterine tumors. We observe that cells resembling stromal cell types in their expression profiles are mainly diploid. However, we also identify intriguing instances of sub-clones within normal cells. In one tumor we find diploid cells with X-chromosome loss accounting for about half the plasma cell component. Upon closer examination of the expression data, we confirmed this DNA-RNA sub-cluster as the clonal expansion of a single B-cell lineage. Making such observations about somatic clonality requires a multi-omic approach.

Hybrid DNA-RNA data expose an extensive range of associations between DNA tumor sub-clones and distinct expression states. Even at this simplest level, these interrelationships span from one-to-one correspondences to more complex many-to-many interactions, and some patterns are strongly suggestive of epigenetic variation. These observations could not be made without a multi-omic approach. Consequently, even deeper analysis was undertaken. We find cases where DNA subclones reveal additional sub-populations within an RNA expression cluster: we separate the RNA expression cluster based on DNA subclones and derive differential gene sets for the subpopulations. These subtle details would also not be observable using RNA-alone data.

Overall, our data highlight the potential of multi-omic technologies to uncover the intricate details of cancer evolution, host response, and stromal mutations. This broad view, coming from single cells alone, can be further honed if we were to use the differential gene sets to explore the spatial relationships using multi-probe high-resolution microscopy^38,39^. Both the broad and detailed view of the inter-connectedness of subpopulations, host and cancer, may enlighten our view of cancer biology and guide future therapeutic strategies.

## Results

### Experimental design

#### Samples

We obtained fresh tissue samples of primary tumor from five patients with uterine cancer (**Table S1**), and in three cases, distal “normal” endometrium (see **Table S2** for a detailed description). The tumor types surveyed include two carcinosarcomas, a serous carcinoma, an endometrioid adenocarcinoma, and a leiomyosarcoma. Each sample was frozen and pulverized into a powder. From this powder, nuclei were extracted for single-cell DNA, single-cell RNA, and single-cell DNA-RNA ("hybrid") BAG platform. We also used the same source material to perform whole genome sequencing (WGS). This comprehensive approach assured that all types of cells were proportionately represented in each method of analysis. To refine our methodology and study **doublet collisions**, we mixed powders from different patients prior to extraction. **Table S2** provides a comprehensive overview of the datasets utilized in our study, detailing the combination of sample origin (including unique setups like the mixed powder experiment), the associated protocols, and the respective experimental parameters.

#### The hybrid BAG platform

Our study uses the BAG platform, a versatile method that captures templates from a single cell entity, either whole cells or nuclei. The BAG platform was built for flexibility, allowing for the reagent customization needed to capture DNA and RNA from the same single cells in a high-throughput manner, an adaptability not found in existing commercial methods. As described in **Figure 1A-D**, nucleic acid templates (or simply “templates”) are captured through primer hybridization to Acrydite-anchored primers embedded into **b**alls of **a**crylamide **g**el, shortened to **BAG**s. This process is followed by primer extension, transcribing the template information of each single cell to primers securely tethered to a single BAG^33^. To establish cell identity, we used a pool-and-split synthesis approach to affix a ***BAG tag*** to each template, randomly assigning one of a million identities (96^3^) to every BAG. During pool-and-split we also introduce a template (also known as varietal tag or UMIs) tag to each template. For the hybrid protocol, we used both oligo-TG primers and oligo-T primers to capture DNA and RNA templates, followed by using DNA polymerase and reverse transcriptase to transcribe the templates onto the anchored primers. We then prepared sequencing libraries by tagmentation.

**Figure 1.**
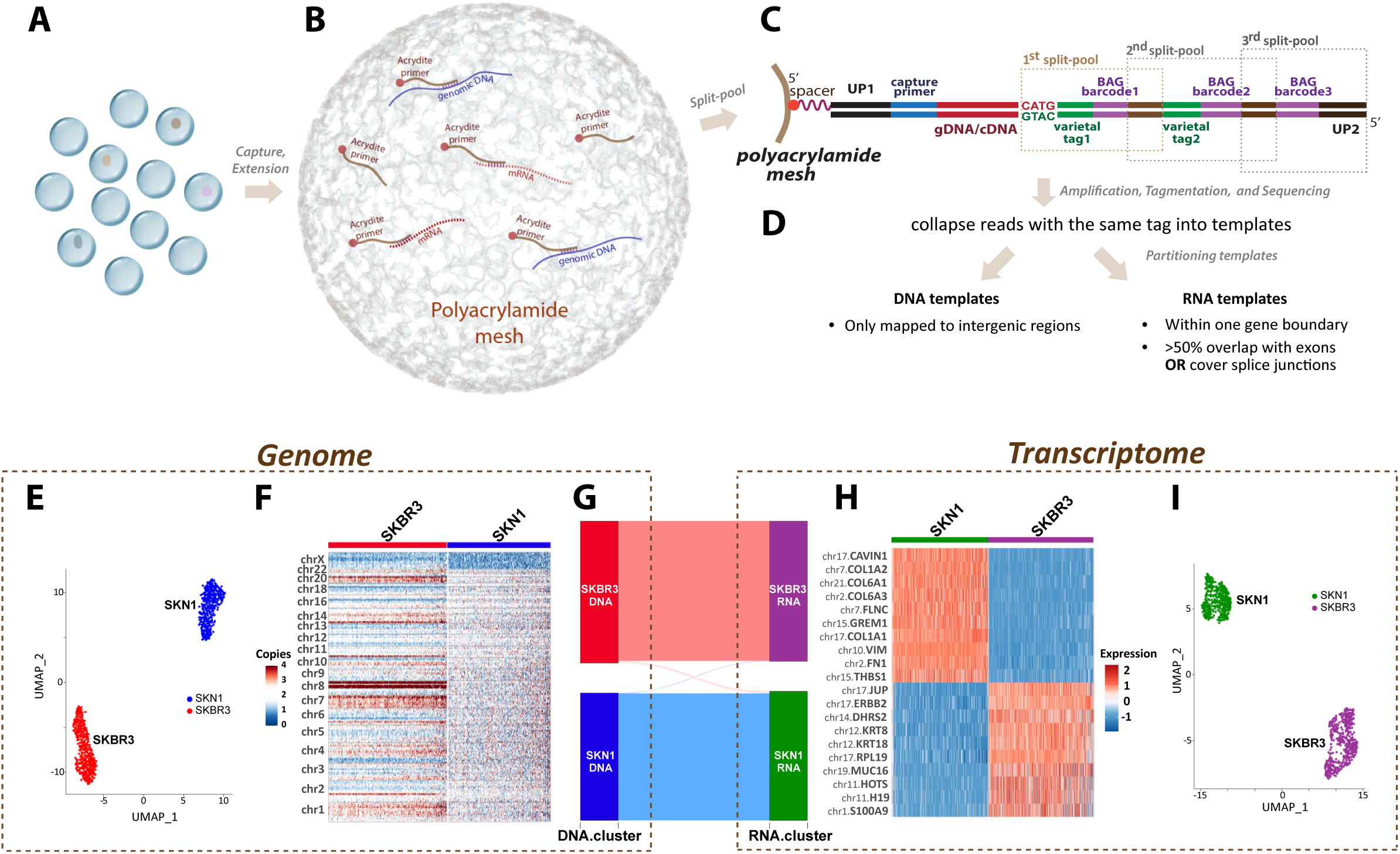
Overview of hybrid BAG-seq protocol and performance on cell-mixture experiment. **A-D:** Key steps for hybrid BAG-seq pipeline. **A**: *Encapsulation*: Individual cells or nuclei are encapsulated within droplets containing acrylamide and Acrydite−modified primers that are designed to capture both mRNA and genomic DNA. **B**: *Polymerization and Primer Extension*: Gel polymerization is followed by primer hybridization. Acrydite primers are extended by reverse transcriptase and DNA polymerase. **C**: *Split−and−Pool Barcoding*: The double−stranded cDNA and genomic DNA are cleaved with a restriction enzyme. During successive rounds of pool−and−split, BAG−specific barcodes (purple) and template−specific varietal tags (green) are added. This process uniquely tags each molecule while assigning a distinct BAG barcode to templates in the same droplet. **D**: *Sequencing and Layer Assignment*: Post−amplification, the molecules undergo tagmentation and subsequent sequencing. The sequencing reads are analyzed for expected structure, tags are extracted and reads with identical varietal tags are collapsed into single templates. Templates are then partitioned into either the DNA or RNA layers based on their mapping characteristics. **E-I**: Performance and genome−transcriptome correlation from a SKN1−SKBR3 mixture single-cell hybrid sequencing experiment: **E**: clustering based on DNA copy number; **F**: heatmap of DNA copy number variations across chromosomes for SKN1 and SKBR3 cells; **G**: correlation between genomic clusters and expression clusters; **H**: heatmap of marker genes of expression clusters, and **I**: clustering result based on gene-count matrix.

#### Genomic filters

All three protocols—DNA-only, RNA-only, and hybrid—require mapping reads to the genome and then organizing the mapped reads based on their BAG and template tags. First, each read is checked for having the expected structure of barcodes and primer sequences. Second, with the tag and primer sequences removed, the rest of the read is mapped to the reference genome. Reads that share a template tag and a BAG tag, and that co-localize in the genome, are collected into ***read aggregates***, and operationally considered a **captured template**. The average number of reads per captured template for each library are also reported in **Table S2**.

For classifying molecules into DNA or RNA origin, templates are assigned to either the RNA or DNA molecular layers based on the composition of their read aggregates. As detailed in the Methods section, templates with largely exonic read aggregates are assigned to the ***RNA layer***, whereas templates with strictly intergenic read aggregates are assigned to the ***DNA layer***. Any remaining templates are marked as ***indeterminate***. Using data from one tumor sample as an example, we demonstrated that DNA clustering identities are very similar (97.6% concordance) whether using all molecules or only DNA-layer molecules as bin counts (**Figure S1**), and a similar conclusion (95.2% concordance) was applied to gene expression clustering results (**Figure S2**) when restricting the analysis to the RNA layer or using all of the RNA templates mapped within transcripts. In our quality control process, we exclude BAGs where the ratio of DNA to RNA templates falls below one-fifth or exceeds five times the average ratio of that library, or those with fewer than 300 RNA-layer molecules. This measure discarded approximately 20% of all hybrid BAGs (for a detailed breakdown, see **Table S2** and **Methods**). We applied these categorization rules to all seven tissue nuclei datasets for all three protocols: hybrid, DNA-only, and RNA-only. The full distribution of each category per sample is shown in **Figure S3A**.

We initially apply the molecular-layer concept to a mixture experiment involving two human cell lines: a normal fibroblast, SKN1, and a breast cancer cell line, SKBR3. The distributions of the basic parameters from this experiment are reported in **Figure S3B-D**. We illustrate the clustering results and heatmaps based on the copy number and gene expression in **Figure 1E-I**. The genomic and transcriptomic features from the hybrid protocol successfully recapitulate the published features of these two cell lines^33,40^. The alluvial diagram (**Figure 1G**) shows the projection of the genomic clones into the expression clusters. As expected, we observed a good one-to-one correlation between the genome and transcriptome of each cell type. Only 1.06% (10 out of 941) of the cells from the DNA cluster of one cell type (either SKN1 or SKBR3) projected to the RNA cluster of the other cell type, probably due to cell doublets.

Despite some **layer spillover**, we observed consistent concordance between the clustering of the DNA layer from the hybrid protocol and the DNA-only data, and similarly between the RNA layer of the hybrid protocol and the RNA-only data. This is true across all five tumor samples as we will discuss in detail in the following Results section: “The hybrid platform is comparable to DNA-only and RNA-only platforms.”

#### Clustering RNA and DNA Layers

We employed the Seurat package^41^ for our single-cell sequencing analysis—a tool widely recognized for its utility in gene expression clustering. For the RNA layer, we followed the standard methodology for expression clustering via “RunUMAP” and “FindClusters” functions. We extended the application of Seurat to cluster the DNA layer. To do so, we first partitioned the genome into large bins (4.7-36.5 Mbp, median = 9.5 Mbp), and then treated the DNA template count within each bin as a “gene input” for the clustering process. This approach allowed us to identify shared copy number profiles that we could leverage in a manner similar to gene expression clustering (further detailed in **Methods**).

#### Multinomial wheel

Whether dealing with DNA or RNA data, a common question often arises: how well does a single cell “fit” within its assigned cluster? To investigate this question within the context of a pre-established set of clusters, we first use multinomial probabilities. We take as a given the ***N*** clusters that Seurat identifies. For each cluster, we sum the gene (or genomic bin) count data over the cells in that cluster and normalize by the total template count. This results in a probability vector that represents the average gene frequency of the cluster. Utilizing this vector, we can calculate the probability that an observed single-cell count vector arose from each of the ***N*** clusters.

However, multinomial probabilities do not translate into a useful metric for deviation from a cluster. Every cell is assigned with almost no ambiguity to one of the major clusters. To properly identify cells that fall between clusters, we incorporate mixed cluster states into a **multinomial wheel**. For every pair of clusters, **A** and **B** with multinomial vectors **p_A_** and **p_B_**, we also consider the mixed state **AB_α_** with multinomial vector **α p_A_** + (1 – **α**) **p_B_** for **α** in [0.1, 0.2, …, 0.9]. This results in a total of 9*(***N*** choose 2) + ***N*** cluster states. By doing this, we can segregate the cells into two categories: **core cluster members** that stay close to an original cluster, and **transitional members** that fall between two clusters.

### The hybrid platform is comparable to DNA-only and RNA-only platforms

In this section, we compare the hybrid BAGs to the DNA-only and RNA-only BAG platforms. We first focus on comparing the hybrid DNA layer to the DNA-only data, and then the hybrid RNA layer to the RNA-only data. We use the tumor tissue sample from Patient 2 as a representative example, with similar comparisons for the other tumor tissue samples provided in **Figure S4-S7**.

#### DNA layer

**Figure 2A** compares the DNA layer from the hybrid data (green labels, left and top) with the data from the DNA-only BAG protocol (red labels, right and bottom). The central scatter plots display the UMAP coordinates, with the hybrid data (in the green box) above and the DNA-only data (in the red box) below. Each point represents a single cell, color-coded by its DNA copy number cluster. Both methods resolved six clusters, which we manually aligned by naming conventions. The **N** cluster includes cells with a typical diploid profile, whereas the **Nx** cluster represents a subpopulation of diploid cells with loss of the X chromosome. The remaining clusters—**A**, **B**, **C** and **D**—exhibit varied aneuploid copy number profiles. The adjacent heatmaps illustrate the distribution of copy number changes across the genome, with deletions in blue, amplifications in red, and the diploid state in white. Each single cell comprises a column and the cells are grouped by their DNA cluster.

**Figure 2.**
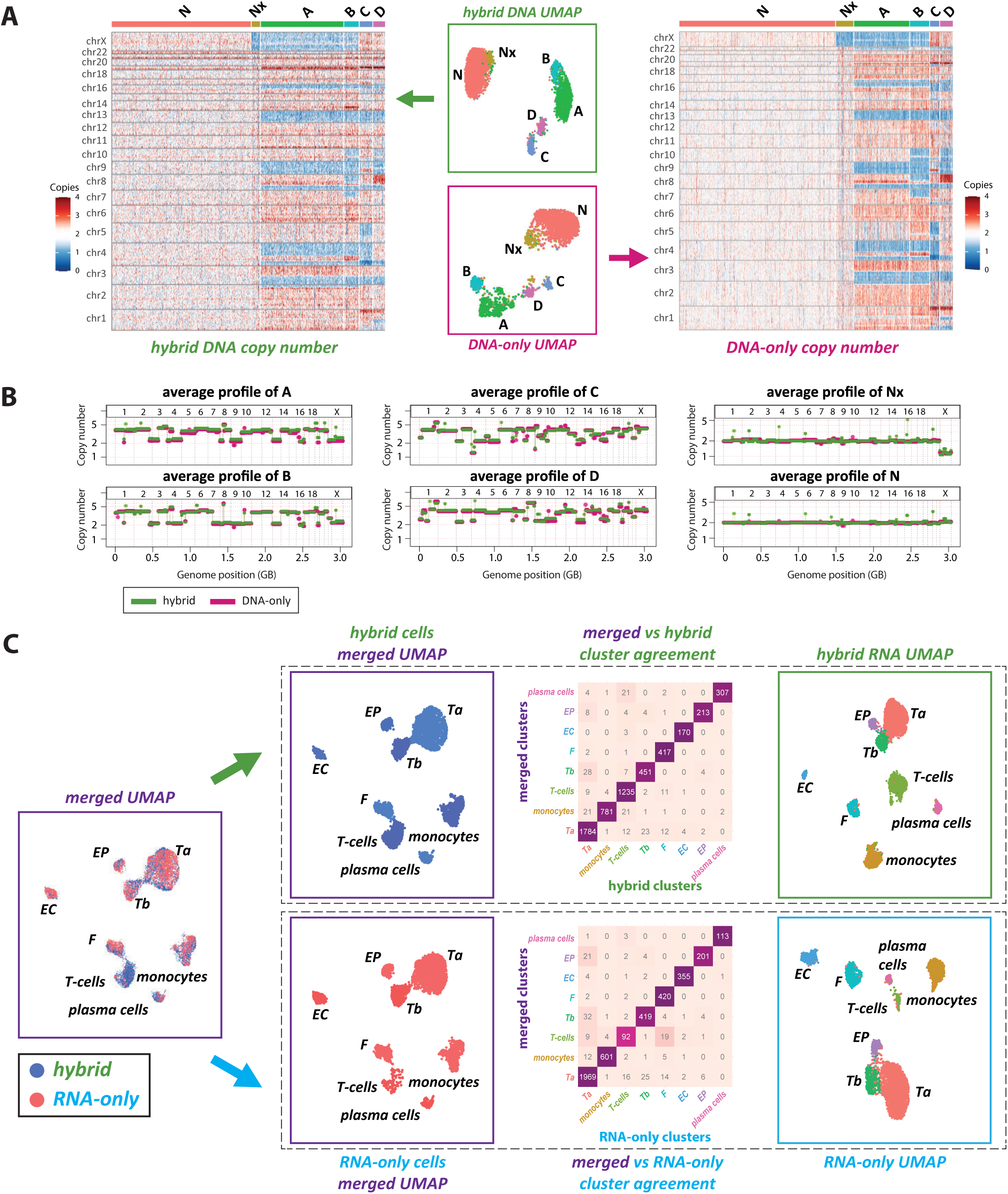
Comparative clustering analysis using the hybrid BAG-seq protocol versus DNA-only and RNA-only protocols. **A**: Single-cell DNA copy number analysis of a tumor sample from patient 2, comparing the hybrid protocol data (green) with the DNA-only protocol data (red). The tumor sample has six distinct copy number profiles: normal diploid cells (N), diploid cells with X-chromosome loss (Nx), and four aneuploid tumor clones (A, B, C, and D). Central plots show UMAP visualizations of single nuclei, color-coded according to cluster identity. Adjacent heatmaps display the binned copy number variations for each single nucleus, arranged by cluster identity on the x-axis against genomic bins on the y-axis, with red indicating amplification and blue indicating deletion. **B**: Aggregated copy number profiles derived from summing across all single nuclei within the same cluster, showing high concordance between hybrid (green) and DNA-only (red) datasets. **C**: RNA expression analysis of the hybrid data (green) compared to RNA-only data (blue). To the far left, hybrid and RNA-only data are co-clustered into eight expression clusters: two tumor expression clusters (***Ta*** and ***Tb***) and six somatic cell types—fibroblasts (***F***), epithelial cells (***EP***), endothelial cells (***EC***), T-cells, monocytes, and plasma cells. These merged clusters are then segregated by hybrid or RNA-only origin. To the far right, each protocol’s data are independently clustered into the same eight expression clusters. The central heatmaps quantify the agreement between the merged clusters with the respective hybrid and RNA-only clusters. Analogous plots for the other four patients are presented in the Supplementary Figures.

To verify the congruence of profiles between platforms, we compared the average copy number profiles for each cluster in **Figure 2B**, with DNA-only data in red and the DNA layer of the hybrid data in green. To quantify the similar clustering results between the two protocols, we used the "Multinomial Wheel" approach to measure the proximity of every single tumor nucleus to the centroids of tumor clones determined both by its own protocol (either hybrid or DNA-only) and by the other protocol. As shown in **Figure S8**, projecting DNA-only data to either DNA-only multinomial states or hybrid multinomial states showed no signal reduction (84.5% versus 84.5% nuclei within 2 units to the centroids), and similarly high concordance if projecting hybrid data to either hybrid multinomial states or DNA-only multinomial states (77.2% versus 68.9% nuclei within 2 units to the centroids). Additionally, we examined heterozygous SNPs in both platforms and found similar patterns of loss of heterozygosity (LoH) and allele imbalance. These allele imbalance patterns (**Figure S9**) align with the copy number calls, in that when the copy number is an odd integer, allele imbalance is always observed.

#### RNA layer

We next analyze the RNA layer of hybrid data compared to the RNA-only nuclei. We first combine all the nuclei from both the hybrid and RNA-only platforms and clustered the integrated datasets into 8 clusters as shown in **Figure 2C** (leftmost “merged UMAP” plot). While each cluster is labeled with a unique identifier, hybrid nuclei are shown in blue, and RNA-only nuclei in red. We then split the merged dataset by experimental origin with hybrid nuclei above and RNA-only nuclei below. The rightmost plots in the panel reflect the clustering of each dataset independently into the same eight identified categories.

We reserve a discussion of the differentially expressed genes for later, but currently label the clusters as ***monocytes***, ***T-cells***, ***F*** (fibroblasts), ***EC*** (endothelial), ***EP*** (epithelial), ***plasma cells***, and two distinct tumor RNA clusters, ***Ta*** and ***Tb***. The central agreement matrix, formatted as a heatmap, shows the consistency of cell classification across platforms within the merged dataset. The top panel correlates hybrid cluster assignments with merged dataset classifications while the bottom does the same for RNA-only data. For both datasets, a significant proportion (95%) of cells align diagonally, confirming that cluster identities are well preserved across the two platforms.

### Profiling genome and transcriptome of tumor samples using hybrid data

For each of the five tumors, we clustered the DNA and RNA layers of the hybrid data respectively (**Figure 3**). The DNA clusters are presented on the far left, accompanied by copy number heatmaps similar to the previous illustrations. On the far right, RNA clusters are displayed, along with a heatmap that illustrates the relative expression levels across sets of differentially expressed genes (blue for low expression, red for high).

**Figure 3.**
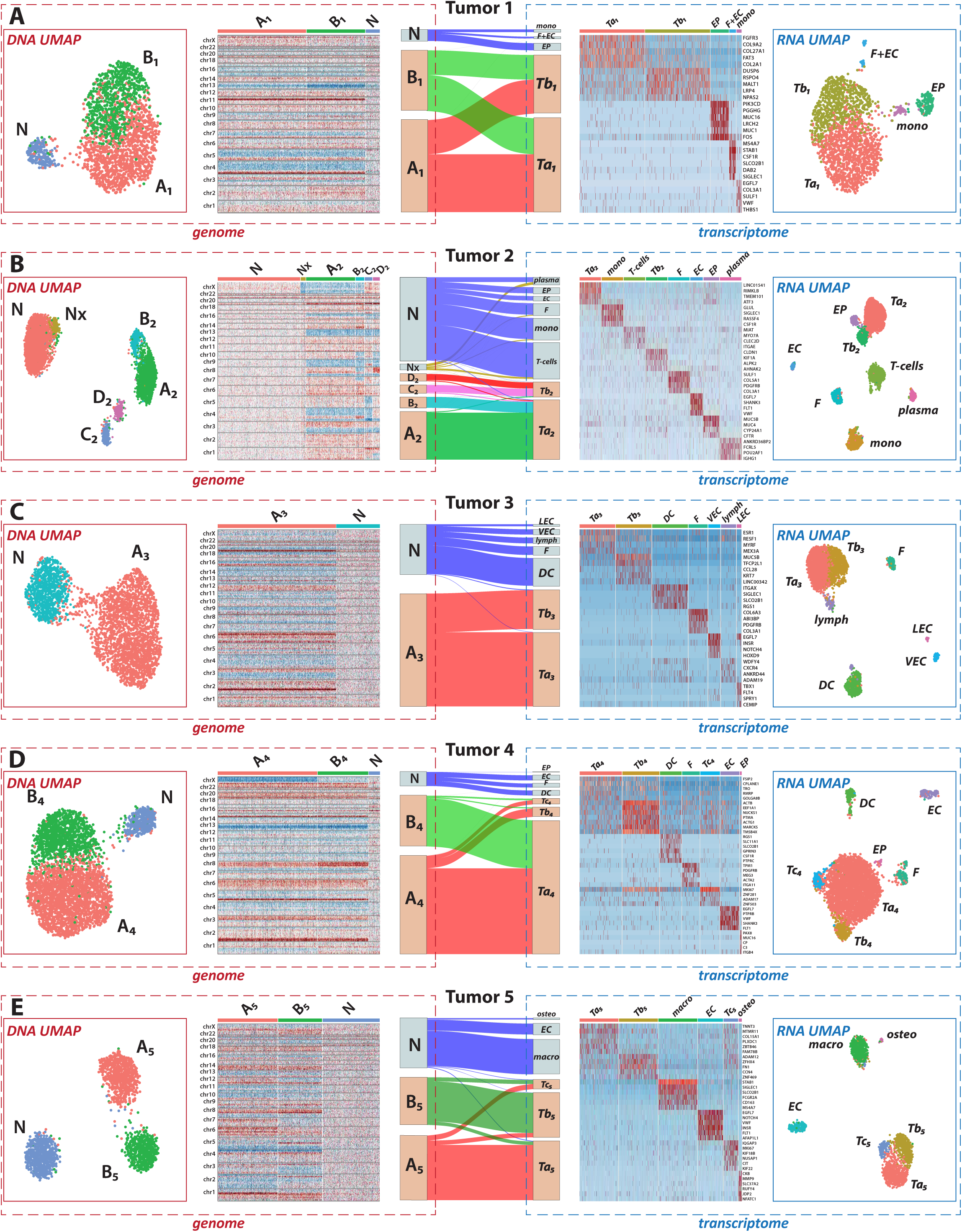
Connecting genomic and transcriptomic data across five tumor samples using alluvial diagrams. **A-E**: Summary of the hybrid data analysis of tumor tissue samples from five individual patients, show the various connections between genomic and transcriptomic landscapes. For each patient:

- The "genome" sections on the left, bordered in red, display the DNA layer information, including UMAP visualizations of single nuclei colored by genomic cluster identities and accompanying heatmaps showing copy number variations across the genome.
- The "transcriptome" sections, bordered in blue, present the RNA layer data, including UMAP plots of single nuclei colored by expression cluster identities. These are accompanied by heatmaps of marker genes that are distinctly upregulated within specific expression clusters.
- At the center of each panel, alluvial diagrams connect the DNA and RNA data, linking genomic cluster identities (left) to the RNA expression clusters (right). The thickness of the flow lines represents the proportion of nuclei that belong to a specific genomic cluster (***X***) and an expression cluster (***Y***), illustrating the integrative analysis facilitated by the hybrid BAG-seq platform.

Capitalizing on the dual DNA and RNA identities assigned to single cells in the hybrid dataset, we quantified the frequency with which cells from a specific DNA cluster appear in a given RNA cluster. This cross-layer association is visualized using **alluvial diagrams**. For example, in panel A from tumor sample 1, the alluvial diagram illustrates that the diploid **N** cells map exclusively with non-malignant expression profiles (***macrophage***, ***F + EC***, and ***EP***). Conversely, both tumor clones **A_1_**and **B_1_** map to the two tumor expression profiles (***Ta_1_***and ***Tb_1_***).

We now delve into the unique characteristics of each of the five tumor specimens, drawing comprehensive insights from both DNA and RNA layers. Focusing on the tumor genome projection patterns, each of the five tumors had a different projection pattern, and we observed almost all the possible patterns. To classify projections, we used a set of letters and numbers to represent the tumor genome and tumor RNA clusters, respectively. For example, [A:1,2; B:2] indicates tumor clone A projected into RNA clusters 1 and 2, whereas tumor clone B from the same primary tumor tissue projected only into RNA cluster 2. We have seen: distinct tumor clones could each project into distinct expression clusters (e.g., [A:1; B:2] for Tumor 5), into shared clusters (e.g., [A:1,2; B:1,2] for Tumor 1), or into a combination of distinct and shared clusters (e.g., [A:1,2; B:1] for Tumor 4). Alternatively, multiple DNA clones could project into a single RNA cluster (e.g., [A:1; B:1; C:2; D:2] for Tumor 2), or a single tumor clone could project into two RNA clusters (e.g., [A:1,2] for Tumor 3). We detail prominent findings and distinctive features uncovered in each case.

In addition, where distinct DNA clusters are associated with a common RNA expression profile, such as illustrated in panel A wherein **A_1_**and **B_1_** are associated with ***Ta_1_***. In these instances, we further dissect the shared RNA expression cluster to test for notable differences in gene expression between the DNA subgroups.

#### Case 1: uterine carcinosarcoma

The tumor 1 was diagnosed as a uterine carcinosarcoma, a complex cancer characterized by both epithelial and mesenchymal components. In our analysis, we observed that over 90% of the cells within the tumor tissue sample displayed significant copy number variations. The DNA layer analysis revealed a normal diploid DNA pattern, **N**, and two distinct tumor DNA clones, **A_1_**and **B_1_**. These DNA clones were differentiated only by a single copy number difference spanning 43 MB on chromosome 13.

At the RNA layer, normal cell clusters were under-resolved by one round of clustering, which we attribute to the high cellularity of tumor in this sample and the low count of normal cells. Despite this, differential gene expression analysis identified three distinct stromal expression clusters: ***monocytes*** (*MS4A7*, *STAB1*, *CSF1R*, *CD14*), epithelial cells (***EP***) (*KRT7*, *MUC1*, *MUC16*, *ELF3*) and a third cluster labeled (***F*** + ***EC***) showing mixed expression of fibroblasts (***F***) (*COL3A1*, *COL6A2*, *COL5A1*, *COL6A1*) and endothelial cells (***EC***) (*EGFL7*, *VWF*, *SULF1*). As detailed in the next section, when combining information from all datasets, this third cluster resolves into 17 fibroblast and 15 endothelial cells. Examining the expression profile of all tumor cells compared to the normal stromal types, we find upregulation of collagen genes (*COL2A1*, *COL9A2*) as well as genes for cell proliferation (*FGFR3*), migration (*PLXNA4*) and growth signaling (*HMGA2*, *PAX2*, *IGF2BP2*).

The tumor cells were divided into two expression subclusters ***Ta_1_***and ***Tb_1_***. Differential analysis between these clusters showed that the ***Ta_1_*** cluster has higher expression of collagen genes (*COL2A1*, *COL1A1*, *COL9A2*, *COL27A1*, *COL1A2*, *COL11A2*). We also observe significant upregulation of *FGFR3*, *AHNAK* and *HIF3A*, genes related to proliferation, structural remodeling, and hypoxic responses. Taken together, these genes suggest the possibility of epithelial-mesenchymal transition (EMT) under hypoxic conditions.

The ***Tb_1_*** cluster, on the other hand, is characterized by higher levels of *MKI67* and *MYC*, suggesting enhanced proliferative capacity and oncogenic signaling. Other genes with increased expression include *FN1*, associated with cell adhesion, and *CENPF*, linked to cell cycle progression. Genes involved in differentiation and growth signaling pathways (DUSP6, MALT1, PCDH7) are also upregulated in this cluster. Similar RNA clustering results were also observed using the 10x Chromium V3 RNA-seq protocol, as shown in **Figure S10**.

When we examined the interplay between the DNA and RNA profiles, we noted that both of the tumor DNA clones, **A_1_** and **B_1_**, contribute equally to the two distinct tumor RNA clusters, ***Ta*_1_** and ***Tb*_1_**. We split the tumor expression groups by their clonal origin. That is, we compare ***Ta*_1_-A_1_**, the cells from expression cluster ***Ta*_1_** and DNA cluster **A_1_**, with ***Ta*_1_-B_1_**, the cells from expression cluster **Ta_1_** and DNA cluster **B_1_**. We also compare ***Tb*_1_-A_1_** with ***Tb*_1_-B_1_**. In neither case do we observe significant expression differences. Similarly, splitting the DNA clusters by RNA identity, we observe no significant difference in copy number between **A_1_-*Ta*_1_** and **A_1_-*Tb*_1_**or between **B_1_-*Ta*_1_** and **B_1_-*Tb*_1_**. Furthermore, when we project the two tumor RNA clusters back into DNA UMAP space, we found that the nuclei from both RNA clusters were randomly distributed in DNA space and were not separated by their DNA cluster patterns; similar patterns were seen when projecting DNA tumor cluster identities onto RNA UMAP spaces. Detailed plots are in **Figure S11**.

### Case 2: uterine serous carcinoma

The tumor 2, identified as a uterine serous carcinoma, presented a more complex DNA and RNA landscape than Tumor 1. The tumor sample served as our example in the previous section comparing the BAG platforms (**Figure 2**). As noted previously, the DNA layer revealed two normal clusters, **N** and **Nx** (diploid with one X-chromosome), and four distinct tumor clusters **A_2_**, **B_2_**, **C_2_** and **D_2_**. Common copy number events across all clones include p-arm deletions on chromosomes 3 and 8 and the deletion of one copy of chromosome 13. **A_2_** and **B_2_** uniquely displayed loss of the X chromosome and deletions on chromosome 4. The allele imbalance data, detailed in **Figure S9,** further clarifies the clonal relationships, with shared and unique loss of heterozygosity (LoH) features among the clusters. These data show that **A_2_**and **B_2_** share a common ancestor that arose after **C_2_**and **D_2_** split.

At the RNA level, various immune cell types (***monocytes***, ***plasma cells,*** and ***T-cells***) along with fibroblasts (***F***), endothelial cells (***EC***), and epithelial cells (***EP***) were distinctly clustered. A comparison between all tumor cells and normal stroma highlighted upregulation of genes involved in cellular signaling (*PTPRF*, *NFKBIZ*, *ARHGAP29*), extracellular matrix remodeling (*LAMA5*, *VCAN*), and antioxidant defense (*SOD2*).

The tumor cells were further divided into two subclusters, ***Ta_2_***and ***Tb_2_***. Using differential gene analysis, we find that ***Ta_2_*** showed upregulation of genes associated with multiple cancer processes (*SGK1*), tumor cell survival and metastasis (*EGR1*), oxidative stress defense (*SOD2*), and vascular development (*RNF213*). In ***Tb_2_***, the upregulated genes related to cell adhesion (*FN1*, *CLDN1*), transcription regulation (*SOX4*) and developmental signaling (*FZD4*). These differences suggest that ***Ta_2_*** comprises a more aggressive and stress-resistant population.

The mapping of the DNA to RNA clusters shows that the normal cells (**N**) map to stromal expression clusters, the diploid cells with one X chromosome (**Nx**) map to the blood elements, and almost all tumor cells (**A_2_**, **B_2_**, **C_2_**, **D_2_**) map to tumor expression categories. From the alluvial plot, we observe that tumor DNA clusters **A_2_** and **B_2_** map to expression cluster ***Ta_2_***, while DNA clusters **C_2_**and **D_2_** map to expression cluster ***Tb_2_***. This is consistent with the phylogeny observed in the DNA data. Additionally, the downregulated expression of *XIST* in ***Ta_2_*** is consistent with the X chromosome loss observed in **A_2_** and **B_2_**.

To explore how DNA identity interacts with expression profiles, we split the cells from the ***Ta_2_*** expression cluster into two subclusters based on their DNA origins: ***Ta_2_*-A_2_**, cells from the ***Ta_2_*** expression cluster and the **A_2_**DNA cluster; and ***Ta_2_*-B_2_**, cells from the ***Ta_2_*** expression cluster and the **B_2_** DNA cluster. Differential gene expression analysis, detailed in **Figure S12A**, revealed distinct expression patterns in these subclusters. In ***Ta_2_*-A_2_**, there is a notable upregulation of genes associated with stress response and cellular metabolism (*HSPB1*, *TXNIP*), cellular excitability (*KCNQ2*), epithelial proliferation (*EPPK1*), and apoptosis (*TNFSF10*). Conversely, ***Ta_2_*-B_2_** is characterized by upregulation of genes involved in ECM remodeling (*COL7A1*, *LAMA1*, *COL6A2*) and transcriptional regulation (*ELL2*).

Similarly, we further segmented the ***Tb_2_*** expression cluster based on DNA identities into two subclusters: ***Tb_2_***-**C_2_**(cells from the ***Tb_2_*** expression cluster and the **C_2_**DNA cluster); and ***Tb_2_***-**D_2_** (cells from the ***Tb_2_*** expression cluster and the **D_2_** DNA cluster). The differential expression analysis, **Figure S12B**, also showed distinct differences in these subclusters. In ***Tb_2_***-**C_2_**, there is a significant upregulation of genes associated with intracellular transport and cell adhesion (*KIF1A*, *ITGA2*), cell motility (*ABR*), and cancer progression (*LAPTM4B*, *KMT2A*). Conversely, ***Tb_2_***-**D_2_** shows increased expression of genes involved in cell adhesion and migration (*FN1*), cellular differentiation (*ALPK2*), apoptosis inhibition (*BIRC3*), and immune system modulation (C3). The projections of tumor DNA clone identities onto the RNA UMAP, along with the projections of tumor RNA cluster identities onto the DNA UMAP, are shown in **Figure S12C**.

An interesting feature of this tumor are the normal cells showing loss of the X chromosome. The differential analysis of **N** and **Nx** restricted to the ***plasma cells***, shows significant differences in expression for *XIST* and *TSIX* in the **Nx** cells, consistent with the loss of the X chromosome. However, we also observed differential expression of the *IGHG1* and *IGHG2*, suggesting a difference in the compositions of these populations. We return to this analysis at the end of the Results section.

#### Case 3: endometrial adenocarcinoma

The tumorfrom Patient 3 was diagnosed as an endometrial adenocarcinoma. Unlike the previous cases, this sample did not present any discernible DNA tumor subclones. The DNA layer analysis distinguished the normal profiles (**N**) from the tumor profile (**A_3_**), accounting for 30% and 70% of the nuclei respectively.

At the RNA level, the stromal cells were clustered into five clusters: dendritic cells (***DC***), lymphocytes (***lymph***), fibroblasts (***F***), vascular endothelial cells (***VEC***), and lymphatic endothelial cells (***LEC***). A comparison of the tumor cells’ global expression pattern with that of the stromal cells revealed marked differences. The tumor cells showed marked upregulation of zinc finger proteins (*ZNF91*, *ZNF100*, *ZNF208*, *ZNF43*, *ZNF431*, *ZNF676*, *ZNF714*, *ZNF738*, *ZNF875*), indicating significant changes in transcriptional regulation. Additionally, genes implicated in endometrial carcinogenesis were notably overexpressed, including those involved in cell growth (*AKT2*), protective barrier formation (*MUC1*, *MUC5B*, *MUC16*), and metabolic reprogramming (*FASN*). The tumor cells show downregulation of *XIST*, consistent with the q-arm deletion on the X-chromosome.

The clustering algorithm further divided the tumor cells into two RNA expression subclusters, ***Ta*_3_**and ***Tb*_3_**. The ***Ta*_3_** was characterized by an upregulation of genes indicative of active cell proliferation and growth (*MKI67*, *ASPM*), and high expression of the estrogen receptor gene (*ESR1*). Consistent with this finding, immunostaining confirmed that approximately 50% of the tumor cells expressed the estrogen receptor (**Figure S13**). The ***Tb*_3_** cluster, with low expression of *ESR1*, showed upregulation of genes related to mucosal barrier function (*MUC16, MUC5B, MUC4*), immune response and chemotaxis (*CCL28*, *SOD2*), and extracellular matrix organization (*FN1*, *COL6A2*).

No significant differences were observed when splitting the **A_3_** DNA cluster based on RNA subclusters ***Ta_3_*** and ***Tb_3_***, suggesting that the gene expression variability within ***Ta_3_*** and ***Tb_3_*** is likely due to regulatory mechanisms other than DNA sequence variation.

#### Case 4: uterine carcinosarcoma

The tumor 4 was diagnosed as a uterine carcinosarcoma. The DNA layer had three clusters: diploid cells (**N**) and two distinct DNA subclones **A_4_** and **B_4_**. Both tumor subclones show similar complex copy number patterns, but with marked differences on the chromosomes 1p, 8, and Xq (**Figure 3D**). In this tumor sample, tumor nuclei comprised 80% of all nuclei.

At the RNA level, the stromal cells were clustered into four distinct groups: dendritic cells (***DC***) (*RGS1*, *SLC11A1*, *STAB1*, *CSF1R*, *ITGAX*, *PTPRC*), fibroblasts (**F**) (*PDGFRB*, *ACTA2*, *COL1A1*, *TPM1*, *MYH11*), endothelial cells (***EC***) (*EGFL7*, *PTPRB*, *VWF*, *FLT1*, *CDH5*), and epithelial cells (***EP***) (*PAX8*, *CLDN1*, *MUC16*, *CP*, *ITGB4*). Comparing expression in the tumor cells to normal stromal cell types, we observe significant upregulation in genes associated with cell proliferation and division (*ASPM*, *CENPF*), transcriptional regulation (*HNRNPH1*, *PABPC1*), and ECM organization (*MXRA5*, *COL2A*). The tumor cells are downregulated for genes involved in antigen presentation (*HLA-B*, *HLA-C*), immune response (*TCIRG1*, *CIITA*), and apoptosis (*MCL1*, *CFLAR*), reflecting potential mechanisms for immune evasion and enhanced tumor survival.

Tumor nuclei were further divided into three distinct expression clusters, ***Ta_4_***, ***Tb_4,_*** and ***Tc_4_***. About 90% of the tumor nuclei map to the ***Ta_4_*** cluster, with roughly equal contributions from the **A_4_** and **B_4_** DNA clusters. Differential expression analysis of ***Ta_4_*** compared to the other tumor subclusters shows upregulation of genes associated with transcriptional regulation (*KCNQ1OT1*, *ZBTB20*, *CPLANE1*), cell structure and motility (*FSIP2*, *SULF1*), and RNA processing (*HNRNPH1*, *SRRM2*). In the ***Tb_4_*** cluster, differential gene expression analysis reveals upregulation of genes related to cytoskeletal organization and cell motility (*ACTB*, *FLNA*, *ACTG1*), as well as those involved in protein synthesis and cellular metabolism (*EEF1A1*, *IGFBP5*, *MARCKS*). For the ***Tc_4_*** cluster, there is a notable upregulation in genes associated with cell proliferation and division (*CENPF*, *MKI67*), as well as transcriptional regulation and developmental processes (*SOX11*, *SOX4*), indicating a sub-cluster with rapid growth, suggesting a proliferative subcluster or replicating cell-state. In the ***Tb_4_*** and **Tc_4_** clusters, 88% and 73% of nuclei map from the **A_4_** DNA profile, respectively. Because of the X-loss in the **A_4_** profile, we observe a relative downregulation of *XIST* and *TSIX* in those subclusters when compared to ***Ta_4_***.

From the alluvial plots, we observe that only expression cluster ***Ta*_4_**shows a significant contribution from both tumor DNA clusters **A_4_**and **B_4_** (61% ***Ta*_4_-A_4_** and 39% ***Ta*_4_-B_4_**). As before, we conducted a differential analysis on ***Ta*_4_-A_4_** and ***Ta*_4_-B_4_** and found only two significant genes: the increased expression of *ANKRD20A8P* in **A_4_** and the increased expression of *PCK1* in **B_4_**(**Figure S14**).

#### Case 5: uterine leiomyosarcoma

The tumor5 was diagnosed as a uterine leiomyosarcoma, an aggressive uterine smooth muscle tumor. The DNA layer of this tumor showed a normal diploid profile (**N**) and two tumor clusters: **A_5_** and **B_5_**, with notable differences across chromosomes 1, 2, 6, 7, 8, 12, and X (**Figure S7A**). We also observed LoH events in **B_5_**not observed in **A_5_** (chromosomes 5q, 16q and 20, **Figure S9E**).

From the RNA layer, three stromal clusters were identified: macrophage (***macro***: *STAB1*, *SIGLEC1*, *CD163*, *CSF1R*, *PTPRC*, *FYB1*), osteoclasts (***osteo***: *ACP5*, *MMP9*, *TCIRG1*, *CTSK*, *SIGLEC15*) and endothelial cells (**EC**: *EGFL7*, *NOTCH4*, *VWF*, *FLT1*, *CD34*, *PECAM1*). Compared to the normal stroma, the tumor nuclei showed upregulation of genes involved in ECM remodeling and cell adhesion (*COL16A1*, *CCDC102B*, *COL1A1*, *COL6A1*, *VCAN*, *POSTN*, *LAMC3*), and several regulatory long non-coding RNAs (lncRNAs) (*DNM30S*, *MEG3*, *KCNQ1OT1*) implicated in oncogenesis.

The tumor cells are further subdivided into three distinct clusters: ***Ta*_5_**, ***Tb*_5_**, and ***Tc*_5_**comprising 53%, 39%, and 8% of the tumor cells respectively. Compared to the other two clusters, ***Ta_5_*** is upregulated for genes related to muscle cell growth (*TNNT3*, *HSPG2*), angiogenesis (*PDGFRB*, *PLXDC1*), ECM remodeling (*COL15A1*, *COL27A1*, *NCOR2*, *POSTN*, *LAMC3*) and different signaling pathways (*ADCY4*, *RGS5*, *ARHGEF17*). Compared to the other two clusters, ***Tb_5_*** has increased expression of a distinct set of collagen genes (*COL1A1*, *COL6A1*, *COL6A2*, *COL6A3*, *COL16A1*) and genes involved in cell adhesion and migration (*ADAM12*, *FN1*, *POSTN*). Finally, the third subcluster, ***Tc_5_***, shows elevated expression of genes involved in cell proliferation and division (*MKI67*, *CENPF*) and transcriptional regulation (*DLX6*, *DLX5*, *ZFHX4, JMJD1C*, *ID2*). This profile suggests that ***Tc_5_*** represents a dividing cell state.

From the alluvial plot, we observe that **A_5_** maps onto ***Ta*_5_** and **B_5_** maps onto ***Tb*_5_**, with very little crossover. However, both **A_5_** and **B_5_**map to the dividing cell state, ***Tc*_5_**, proportionate to their representation in the tumor sample.

### Combined clustering of all patient samples

#### Integrative clustering analysis and cluster phylogeny

In our previous sections, we examined one patient at a time, while in this section, we aim to understand how patient expression profiles interrelate and potentially achieve finer resolution in our clustering analysis. To gain such a global picture of the hybrid data, we performed cluster analysis on the combined data from all patients, separately for RNA layer and DNA layer, demonstrated in **Figure 4**. For RNA layer clustering, we lowered the threshold for RNA template count from 400 to 300. This analysis includes hybrid data from the five tumors (**Tumor *i***) presented in the previous section, as well as normal tissue samples from patients 1, 2, and 4 (**Normal *i***). We then explore which expression clusters map to which patient and within each patient to which genome cluster, diploid copy number (**flat**), diploid copy number with one chromosome X (**X-loss**), or complex aneuploid genome (**CN+**). In this way, we reveal diverse and common cellular and functional repertoires. These results are summarized in **Table 1**.

**Figure 4.**
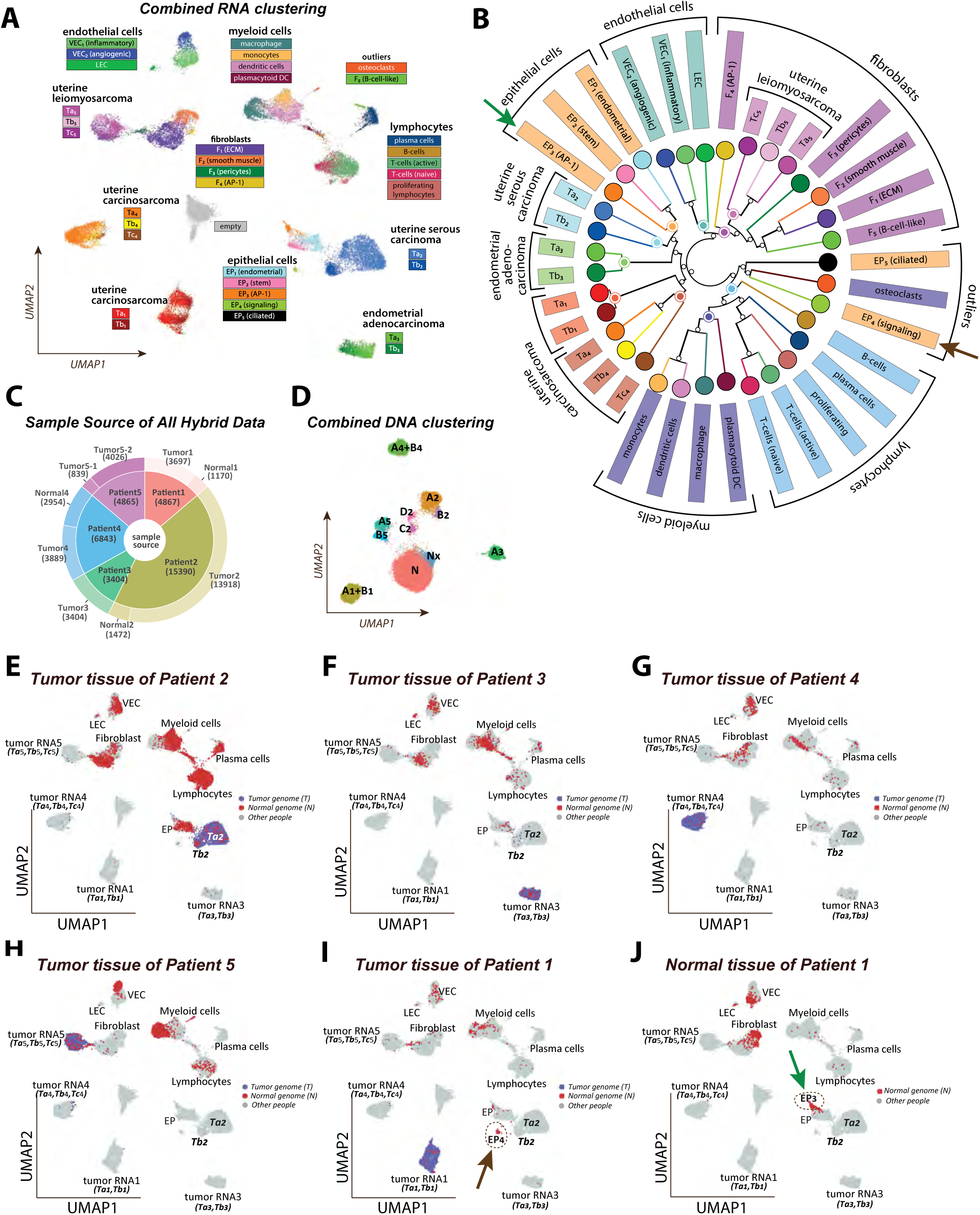
Integrated cluster analysis and unique cluster identification using aggregate tumor and normal tissue data from all patients. **A**: UMAP scatter plot showing the RNA layer data from both tumor and normal tissue samples across all patients, totaling 40,149 nuclei. Stromal clusters are defined through iterative sub-clustering, while tumor sub-clusters are defined from individual tumor analyses as presented in Figure 3. An empty cluster (grey) consists of nuclei mainly from the RNA layer of seven DNA-only experiments. Each point is colored according to the cluster with the highest likelihood as determined by a multinomial model. **B**: The neighbor-joining tree illustrates the relationships among the stromal and tumor sub-types. The tree is computed from inter-cluster distances based on multinomial distributions. **C**: The source of nuclei from the hybrid protocol used in the combined analyses. **D**: Combined DNA clustering after removing nuclei clustered to the “empty” state of the RNA clustering in (A). **E-J**: The tumor-genome (blue) and normal-genome (red) nuclei projections on the RNA UMAP space for six tissue samples. The tumor-genome or normal-genome information is determined by the combined DNA analysis in (C). Unique stromal components specific to certain tissues are circled in dashed lines and indicated by arrows in panels (I) and (J), and marked in (B).

**Table 1.**
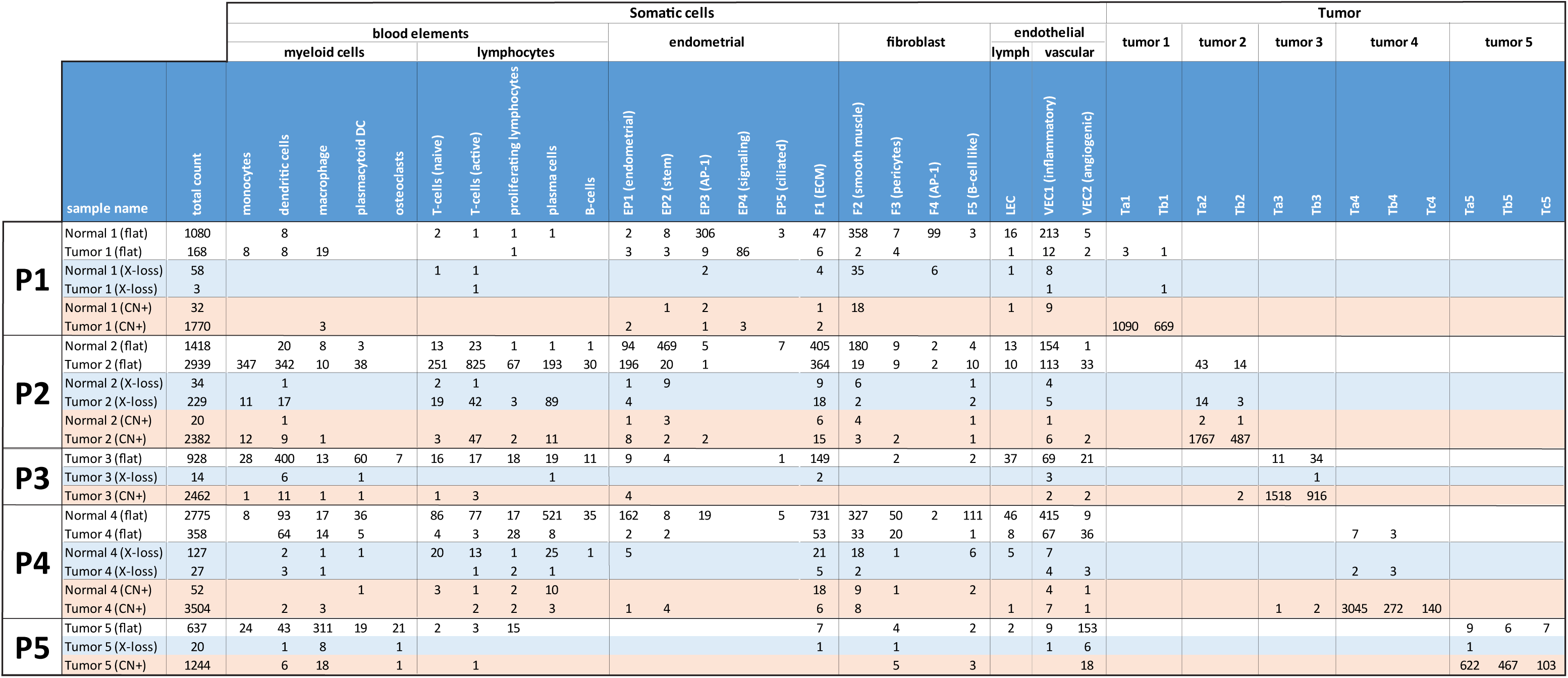
Co-clustering counts across all samples. This table presents the distribution of nuclei for each patient (P1 to P5) and tissue sample (Tumor or Normal) categorized by DNA cluster: diploid (flat, white), diploid with one X chromosome (X-loss, blue) and aneuploid (CN+, red). It details the total count of nuclei within each category, along with sub-counts for each expression cluster.

Specifically, our gene expression clustering procedure incorporates several novel features, which we detail below for clarity. First, we include 3500 “empty” cells derived from the RNA layer of DNA-only BAGs. These empty cells were clustered together in **Figure 4A**. This cluster also attracts hybrid BAGs with severe RNA template depletion. Overall, 2.4% of hybrid cells coalesce into the **empty state**.

As before, we use Seurat’s FindClusters and UMAP functions to cluster and display 40,149 nuclei, which uses the UMAP coordinates to visualize the single cells, as shown in **Figure 4A**. Detailed Seurat clustering parameters can be found in the Methods section. After the initial clustering, we further resolved the diploid cells to delineate subpopulations, described in the next section. Specifically, we chose four sub-regions of the initial UMAP (blood elements, fibroblasts, endothelial cells, epithelial cells) for further sub-clustering. Given the single-cell pool’s diverse cell type composition – ranging from tumor, epithelial, endothelial, myeloid, etc. – an iterative clustering approach is a reasonable strategy. For the tumor subclones, we used the cluster information from the previous section.

We use UMAP to see a planar representation of the cells, and our method of multinomial analysis to color the cells by expression type. The aggregate of templates in each sub-cluster forms a **gene probability vector**, wherein the total number of templates mapping to a gene is normalized by the total number of templates mapping to *any* gene, resulting in a probability distribution. This probability vector determines a multinomial distribution for each cluster that in turn, assigns a cluster probability for each single cell based on its gene counts. In **Figure 4A**, each point’s color reflects the cluster most likely to have generated the gene counts in that cell.

We then use the multinomial distributions to establish a distance metric between any two sub-clusters, each with its own multinomial distribution. The asymmetric disparity between two multinomial distributions, A and B, is determined by simulation, in which we determine the number of reads of a cell simulated from distribution A that are needed to preferentially assign the cell to A rather than to B, for some given confidence threshold. We then symmetrize the disparity measure, combining A to B and B to A. Clusters with distinct profiles will require fewer reads to establish separation, while those with similar profiles require more reads for differentiation. We invert this measure to establish a pairwise distance (**Figure S15**) and use neighbor-joining to build a phylogeny over the expression clusters (**Figure 4B**). We rank the genes that best separate two sets of expression clusters, **A** and **B**, using binomial methods, rather than the FindMarkers function within Seurat. We use the ratio of total reads in **A** and **B** to determine the null expectation of the ratio for any given gene. We then apply a binomial test to the observed counts for each gene in **A** and **B**. While there exist more advanced methodologies, the ranking of the top 20 genes would not significantly vary. The marker genes distinguishing tumor expressional sub-clusters for each patient can be found in **Table S3**, while those for stroma clusters across all five patients are listed in **Table S4**.

On the DNA front, from a total of 35,369 nuclei from the sources indicated in **Figure 4C** studied using the hybrid protocol, after removing the 2.4% (859 out of 35,369) low-quality nuclei that were clustered into the empty state in the gene expression clustering result, we cluster the DNA-layer all the remaining nuclei based on genomic bin counts. Similar to the RNA clustering result, in the DNA space in **Figure 4D**, tumor-genome clusters were quite distinct from the normal-genome clusters, and distinct clones within a patient mapped nearby to each other or merged into a single cluster at this resolution of clustering.

Combining RNA and DNA clustering information, the projections of nuclei from six sample sources into the combined RNA space (**Figure 4A**) are shown in **Figure 4E-J**, where nuclei from a given sample are highlighted either in blue or red, depending on whether they are classified by DNA as tumor or normal genomes. In each panel, the nuclei from other samples are colored in light grey. The projections of the tumor genomes are very distinct between patients, well-separated from each other and the projections of the normal genomes.

Although the normal-genome stromal clusters are mostly shared between patients, there are several gene expression clusters that only belong to certain tissues and certain patients, implying effects from tumor microenvironments and personal genomes. For example, the epithelial-like cluster EP_4_ (**Figure 4I**, pointed to by the brown arrow) from the tumor tissue of patient 1 is significantly different from the EP_3_ cluster (**Figure 4J**, pointed to by the green arrow) from the distal normal uterine site from the same patient, and they are both distinct from the main epithelial cluster EP_1_. These two clusters are also pointed to by arrows in the hierarchical wheel in **Figure 4B**, far away from each other. We believe these distinct clusters are not due to batch effects not only because they have very distinct and plausible gene expression patterns (**Table S4**), but also because these distinct clusters overlapped well in experimental replicates and showed the same patterns in RNA-only datasets (**Table S2**). These unique stromal subpopulations will be discussed in detail in the following section.

#### Common and unique features in stromal subpopulations

In the aggregation of the single-cell data, the tumor cells and diverse stromal cell types established distinctive clusters. In the multinomial tree (**Figure 4B**), the branches preserve cell-type categories. The blood elements share a common branch (blue labels) with a myeloid-derived sub-branch (dark blue) distinct from lymphocytes (light blue); a branch of epithelial cells (orange); fibroblasts (purple) and endothelial cells (green). Nearly all subclusters fall on their expected branch. The exception is a single sub-branch containing two epithelial sub-clusters, EP_4_ and EP_5_, and osteoclasts, a myeloid cell type. In general, clusters that are close together in the UMAP (**Figure 4A**) share a common branch in the tree (**Figure 4B**). The major exception is cluster F_5_, B-cell-like fibroblasts, which are near the B-cells in the UMAP but nearer the fibroblasts in the tree.

The tumor clones from each patient occupy distinct sub-branches in the tree. The uterine leiomyosarcoma (purple), a muscle-derived tumor, has expression subclusters on the fibroblast branch of the tree. The other four tumors share a deep branch with the epithelial cells. One sub-branch contains the two uterine carcinosarcomas (red and dark red). Nearer the epithelial cells in the tree, are the endometrial adenocarcinoma (green) and nearer still the uterine serous carcinoma (blue). The branch lengths provide a relative measure of similarity, showing that Ta_1_ and Tb_1_ are highly similar as are Ta_3_ and Tb_3_. In contrast, the subclones of Ta_2_ and Tb_2_ are far apart.

Next, we examine the expression state differences that contribute to the unique identities for each of the subclusters in the blood components, fibroblasts, epithelial, and endothelial cells.

##### Blood elements

We begin by exploring the subdivisions of the blood cells which comprise a major component of the somatic cell lineages across all the cancer samples. The additional sub-clustering step and differential gene analyses identify nine distinguishable sub-clusters. Several of the subclusters form a lymphoid lineage super-cluster including mature **B-cells** (*CD20*, *BANK1*, *CD19*, *CD22*), **plasma cells** (*IHGM*, *IGHG3*, *IGHG1*, *IGHA1*), **proliferative lymphocytes** (*MKI67*, *ASPM*, *TOP2A*, *EZH2*, *POLQ*), and two sub-populations of T-cells: **naïve T-cells** (*IL7R*, *ITK*, *TCF7*, *CCR7*) and **active T-cells** (*CD103*, *CCL3*, *CCL4*, *CD8A*). We also observe a second myeloid super-cluster comprised of **monocytes** (*SAT1*, *CD169*, *LRP1*, *STAT1*), **dendritic cells** (*CD11c*, *SLC11A1*, *CTSB*, *ADAM8*), **macrophage** (*CD163*, *CSF1R*, *CD169*, *FCGR2A*), and **plasmacytoid dendritic cells** (*CSF2RA*, *CD11c*, *IRF8*, *IRF7*, *CD74*, *CD13*). We also identify a cluster of blood elements with a unique profile that does not cluster with the other blood elements. The expression profile is suggestive of **osteoclasts** with marked upregulation of genes involved in bone-resorption (*TCIRG1*, *ACP5*, *CTSK*).

Examining the distribution of blood elements by sample (**Table 1**), we see very different distributions of immune cell types by sample and location. Patient 1 tumor tissue sample and its paired normal have the fewest blood cells, with some monocytes and macrophages in the tumor sample. Patient 2 has low counts of blood elements in her normal tissue sample but high concentrations of monocytes, dendritic cells, B-cells, and T-cells, and a large population of actively dividing lymphocytes in her tumor sample. Patient 3, with no paired normal, has a significant proportion of dendritic and plasmacytoid dendritic cells. Patient 4 tumor sample and her paired normal have high concentrations of immune cells. B-cells and plasma cells are at higher concentrations in the paired normal, whereas the dividing lymphocytes and dendritic cells are at higher concentrations in the tumor. Patient 5, with no paired normal, has a significant macrophage and osteoclast population in her tumor sample. Only Tumor 3 also contains some osteoclasts, but at a much lower amount.

##### Epithelial cells

The clustering identified five distinct epithelial cell clusters that form into two very distinct subclusters in the expression tree. The majority of cells fall into cluster **EP_1_**, wherein the differential analysis strongly suggests **endometrial epithelium** (*PAX8*, *MUC1*, *MUC16*, *MUC4*, *MUC5B*, *LAMB3*, *LAMA3*). Cluster **EP_2_** is similar to EP_1_, but shows an increased expression of **stem-cell markers** (*LGR5*, *PODXL*) and hormone receptors (*PGR*, *ESR1*). Cluster **EP_3_** shows activation of the **AP-1 pathway** (*FOS*, *JUN*, *FOSB*, *JUND*) and other cellular response genes (*BTG2*, *EGR1*, *NR4A1*, *RN7SK*). The other two epithelial cell clusters show distinctive patterns of expression: **EP_4_** shares upregulation of common endometrial epithelial markers (*PAX8*, *MUC16*, *KRT7*) but also markers of **signaling** (*PIK3CD*) and response to stimuli (*FOS*, *HSPA5*, *HSPB1*). The **EP_5_** cluster contains a very small number of cells; however, their profiles are so specific that they are clearly differentiated in this analysis. Their differential expression profile strongly suggests that these are **ciliated epithelial cells** with upregulation of dynein heavy chain genes (*DNAH9*, *DNAH12*, *DNAH11*, *DNAH7*) and cilia and flagella associated genes (*CFAP157*, *CFAP43*, *CFAP47*, *CFAP46*) and other ciliary genes (*SPAG17*, *TEKT1*, *HYDIN*, *DNAI1*).

When we look at the distribution of epithelial cell types across patient tissues, we find an interesting pattern of expression in the tumor tissue of Patient 1. The epithelial cells from the paired normal are predominantly from EP_3_ (activation of the AP-1 pathway) while the epithelial cells from within the tumor sample are predominately EP_4_ (genes for signaling and response to stimuli). This suggests that the epithelial cells in the tumor microenvironment have significantly altered expression. Most of the EP_2_ cells (stem-like epithelial cells) are present in the paired normal from Patient 2. The EP_5_ ciliated cells were scattered throughout tumor samples of Patients 1-4 while the tumor sample of Patient 5 had no epithelial cells at all.

##### Fibroblasts

The fibroblast-like cells split into five distinct clusters. The first cluster, **F_1_**, shows upregulation of collagen genes (*COL6A1*, *COL7A1*, *COL6A2*, *COL1A1*, *COL3A1*, *COL6A3*, *COL1A2*) and genes involved in organization and production of extracellular matrix (*THBS1/2*, *AEBP1*, *VCAN*, *SERPINE1*) suggesting a role in **ECM** remodeling. Cluster **F_2_** appears to be comprised of **smooth muscle** cells, with upregulation of collagen genes, but also genes for contractile proteins (*CNN1*, *ACTA2*, *MYH11*, *TPM1*) and signaling pathways for muscle contraction (*PPP1R12B*, *CACNA1C*, *MYLK*). The expression profiles for the cells in cluster **F_3_**are suggestive of **pericytes** with classic markers (*PDGFRB*, *RGS5*), Notch signaling (*NOTCH3*, *HEYL*, *JAG1*), and ECM and membrane genes (*COL4A1*, *MCAM*, *CSPG4*). The **F_4_** cluster is a fibroblast or myofibroblast (*COL1A1*, *COL1A2*, *FN1*, *THBS1/2*, *CCN1/2*) that shows activation of the **AP-1 pathway** (*FOS*, *FOSB*, *JUN*) and other early response genes (*EGR1*, *NR4A1*, *KLF4*).

There is one outlier in this cluster: **F_5_** shows characteristics of both **fibroblasts and B-cells**. When we compare F_5_ to the other somatic cell types, we observe upregulation of genes associated with B-cell and plasma cell genes (*IGHG1*, *IGHM*, *IGHG3*, *FCRL5*, *POU2AF1*, *MZB1*). However, when we compare the F_5_ cluster to the lymphoid cluster as a whole, we find that F_5_ is highly enriched for collagen genes (*COL6A1*, *COL6A2*, *COL5A1*, *COL1A1*, *COL3A1*) and extra-cellular matrix (*THBS1*, *THBS2*).

The ECM fibroblasts (***F_1_***) were found in the stromal populations of all five tumors, but the other fibroblast populations were variably distributed. The smooth muscle cells (F_2_) were found primarily in Tumors 1, 2, and 4. Pericytes (***F_3_***) were found in all tumors but at highest concentration in Tumor 4. The fibroblasts showing AP-1 activation (***EP_4_***) were only present in the normal distal tissue of Tumor 2, the same sample in which we observed AP-1 activation of the endometrial cells (EP_3_). The hybrid B-cell/ECM cluster F_5_ are mostly from the normal adjacent tissue of Tumor 4.

##### Endothelial cells

The endothelial cells fall into three distinct clusters. The first cluster labeled, ***LEC***, shows upregulation of common markers genes for **lymphatic endothelial cells** (*FLT4*, *PROX1*, *CCL21*, *TBX1*, *EGFL7*). The other two clusters ***VEC_1_*** and ***VEC_2_*** show the expression patterns of vascular endothelial cells (*VWF*, *FLT1*, *ROBO4*, *NOTCH4*, *SEMA3F*, *HSPG2*). The differential expression patterns of ***VEC_1_*** suggest an inflammatory response (*SLCO2A1*, *FN1*, *ZFP36*, *KLF2*, *SELE*) whereas ***VEC_2_*** is upregulated for structural genes (*COL4A1*, *COL27A1*, *LAMA4*), vascularization and angiogenesis (*ESM1*, *INSR*, *ANGPT2*, *FLT1*). All of the endothelial cell types are found in each patient. The lymphatic endothelial cells (***LEC***) are at highest concentrations in samples from Patients 2, 3, and 4. We find the inflammatory vascular endothelial cells (***VEC_1_***) are abundant in all but patient 5. The sample from Patient 5 has the highest concentration of structural/angiogenic endothelial cells (***VEC_2_***).

### Multinomial wheel and crossovers

In the previous section, we observed that the majority of nuclei exhibit concordant DNA and RNA profiles: diploid DNA with somatic RNA expression (**flat**, ***N***) or complex DNA patterns with tumor RNA expression (**CN+**, ***T***). These **concordant** nuclei constitute the expected biological behavior; however, there are a subset of cells that do not match this pattern. **Table S5** shows the counts for each patient of cells that are diploid or complex (**flat** or **CN+**) and map to normal clusters or tumor clusters (***N*** or ***T***). Across all patients and tissue samples, 1-5% of nuclei have flat copy number profiles and tumor expression patterns or copy number variations and somatic expression patterns. Although accounting for a small proportion of the total dataset, these **crossover** nuclei may represent an interesting population. Alternatively, they may be the result of unresolved doublet collisions^42^. **Table S6** totals the counts and calculates the proportion of concordant and crossover nuclei per patient.

To determine if crossover nuclei are a unique biological state or collision artifacts, we employed the multinomial wheel to differentiate mixed states. For the DNA layer, we constructed a multinomial wheel with tumor clusters, diploid cells and diploid cells with X-loss as the individual spokes of the wheel (**Figure S16A**). Some tumor DNA clusters are so similar (**A_1_** and **B_1_**, **A_4_**and **B_4_**) that we collapse each of those pairs into a single node (**AB_1_** and **AB_4_**). This results in a multinomial wheel with 11 spokes and 9*(11 choose 2) = 495 intermediate states.

From the DNA counts of each nucleus, we evaluate the probability that those observations derived from each of the (495 + 11) = 506 multinomial distributions in the wheel. We assign each nucleus to the node with the highest posterior probability. Notably, better than 60% of nuclei align with a ‘pure’ cluster on the multinomial wheel, and 85% are situated within two units’ distance from a pure cluster (**Figure S16A**). Informed by two mixture experiments, detailed in the **Methods** and **Figure S16B-E**, we apply a **collision filter**, marking for removal nuclei that are 3 or more units from a spoke *and* fall between a normal DNA cluster (**N**, **Nx**) and a tumor DNA cluster (**A_i_**, **B_i_**, etc). **Tables S5 and S6** (filtered nuclei) show the counts after applying the collision filter.

If crossovers are the result of cryptic collisions, the collision filter will disproportionately reduce the frequency of crossovers compared to the concordant nuclei between the DNA and RNA layers. Indeed our observations align with these expectations: while the collision filter removed 14% of concordant nuclei, it removed a substantial 71% of crossover nuclei (**Table S6**). Removing collisions does not alter the results presented in the previous section. A version of **Table 1** restricted to nuclei that pass the collision filter can be found in **Table S7**.

### Loss of X chromosome in somatic lineage

In this final analysis, we revisit the occurrence of X chromosome loss in diploid cells, particularly noted in the plasma cell population from Patient 2’s tumor sample. X chromosome loss, while reported in some cancers^43^, and in the peripheral blood of some cancer patients which may be related to aging^44^, presented an unusual prevalence in our dataset among the stromal cell types in the tissue. For example, one-third of the plasma cells in Patient 2 exhibited X chromosome loss, a finding that merited further investigation into its biological implications.

To ascertain whether the X chromosome loss was the result of a clonal event or a random convergent state, we performed a haplotype analysis: If the cells all lost the same X chromosome, the nuclei are likely related whereas if they lost different copies of the X, this is probably a convergent state. The tumor genome from patient 2 has two subclones with X-loss (**A_2_**, **B_2_**), and we used these two subclones to phase the SNPs on the X chromosome. Upon aggregating SNPs within each nucleus, we found that the ***Plasma*-Nx** nuclei share a common haplotype, whereas the ***T-cell*-Nx** population shows both haplotypes (**Figure 5A**). Given the robust coverage of the ***Plasma*-N** and ***Plasma*-Nx** sub-clusters, we were able to conduct a differential analysis on their aggregate gene expression data. Shown in **Figure 5B**, there are four genes with significantly different expressions: *XIST* and *TSIX* are under-expressed in the X-loss population, affirming the X-loss observed in the DNA layer. The other two genes are *IGHG1* and *IGHG2*, hinting at a potential compositional difference between these plasma cell populations. As *XIST* RNA is required for X-chromosome inactivation and is only expressed from the inactive X-chromosome^45–47^, these results also verify that nuclei from the “Nx” DNA clone lost their inactivated X chromosomes.

**Figure 5.**
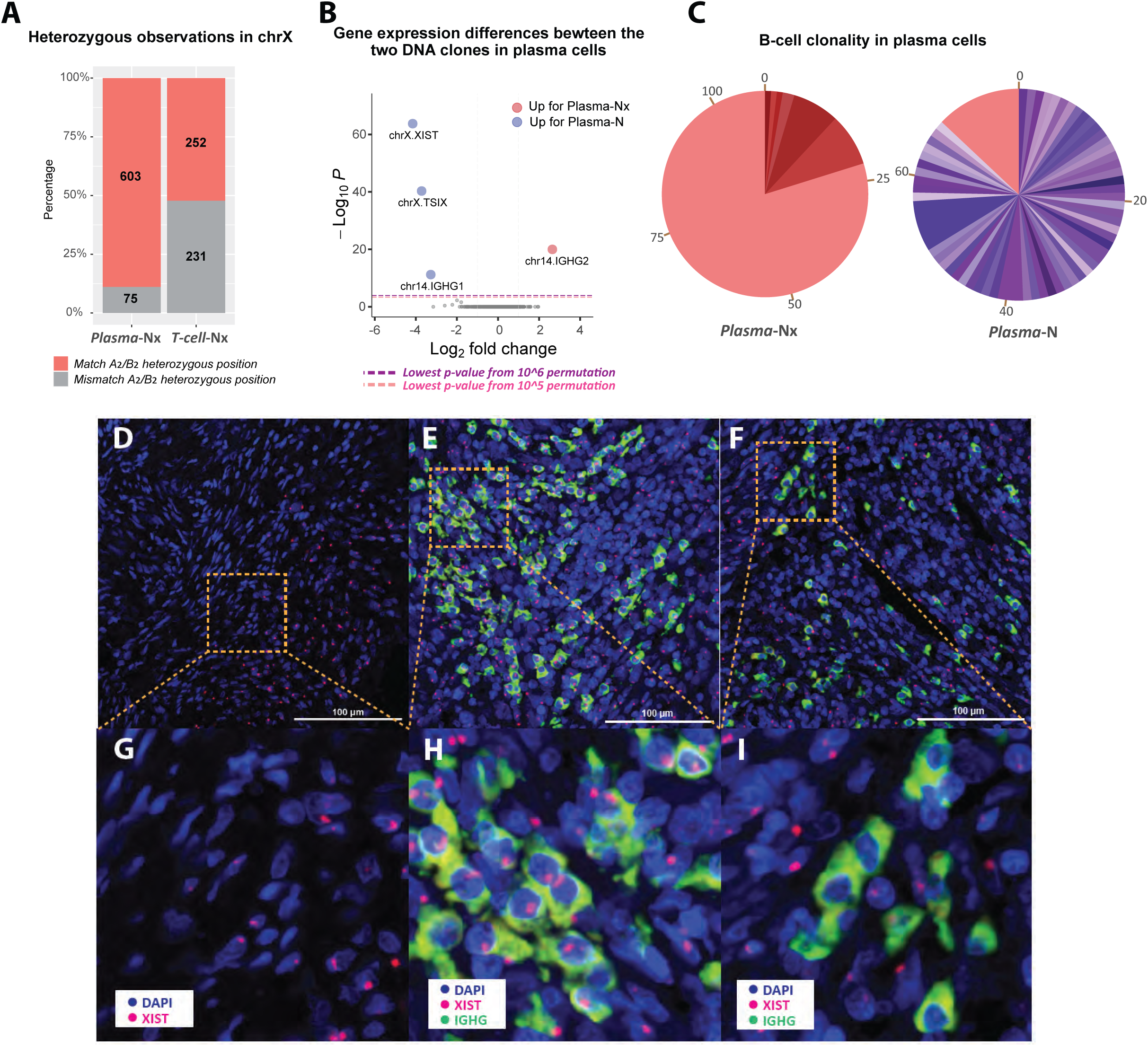
Analysis of chromosome X loss in somatic cells of the primary tumor 2. **A**: Bar plot showing the ratio of X-haplotype observations in the X-loss populations of Plasma (Plasma-Nx) and T-cell (T-cell-Nx) nuclei from patient 2. Tumor subclones A2 and B2 with only one copy of the chromosome X are used to phase the X chromosome SNPs in the Plasma-Nx and T-cell-Nx populations as belonging to one haplotype (red, match A2/B2) or the other (grey, mismatch A2/B2). T-cell-Nx nuclei exhibit a balanced distribution of SNVs from both haplotypes, while Plasma-Nx nuclei show a pronounced bias toward the A2/B2 haplotype. **B**: A volcano plot shows the genes with statistically significant expression differences between diploid plasma cells (Plasma-N) and plasma cells with one X chromosome (Plasma-Nx). **C**: Pie charts showing VDJ recombination results for Plasma-N and Plasma-Nx nuclei. Each color represents a unique B-cell clone identified by its CDR3 sequence, with the size indicating the clone’s prevalence. The Plasma-N chart shows a diverse clonal makeup with few dominant clones, while the Plasma-Nx chart shows a clear dominance of one lineage (red), constituting 80% of the population and matching the signature of the primary clone in Plasma-N. **D-I**: Spatial transcriptomics images illustrating XIST expression in the same tumor tissue of patient 2. **D**: RNAscope images displaying the expression of XIST (red) along with DAPI staining of nuclei, showing half of the cells with no XIST signal detection. **E-F**: IGHG (green, plasma cell marker) and XIST (red) expression along with DAPI staining of nuclei. **E** shows a region consisting of normal plasma cells with XIST gene expression, while **F** shows another region of plasma cells with no XIST red dots detected. **G-I** are zoomed-in images of (D-F), respectively.

Considering the distinct IGH expression in the plasma subclusters, we further investigated the clonality of the ***Plasma*-Nx** population through VDJ recombination analysis. Using MiXCR^48^ to count the unique VDJ recombination patterns, our analysis strongly suggests that the ***Plasma*-Nx** cells are primarily derived from a singular B-cell lineage expansion, in contrast to the ***Plasma*-N** cells, which originated from a diverse array of lineages (**Figure 5C**).

To corroborate our findings and further visualize the loss of the X chromosome, we employed RNAScope technology on the same tumor tissue from Patient 2. The spatial transcriptomics images (**Figure 5D-I**) provide a vivid illustration of the XIST expression patterns. DAPI staining (blue) marks the nuclei, while XIST (red) and IGHG (green) show specific gene expressions. In **Figure 5D**, half of the cells display no XIST signal, indicating the X-loss, likely from tumor A_2_ or B_2_ clones with deletions in the X chromosome. **Figure 5E** and **F** show IGHG, the plasma cell marker, and XIST expression within two different regions, showing two opposite phenomena. While most of the plasma cells in **Figure 5E** contain the XIST signal, implying the presence of the inactivated X chromosome; the majority of the plasma cells in **Figure 5F** lack the XIST signal, suggesting that these cells have lost the inactivated X chromosome. Both the plasma cells with or without XIST signals were found spatially localized in the tumor microenvironment.

We next extended our X-loss analysis to the other tissue samples. For patient 1, we noted a 5% X-loss in the diploid component of the normal distal sample, predominately in the smooth muscle (***F_2_***) expression cluster. However, establishing clonality in this context was challenging due to the tumor’s retention of the X chromosome in its subclones.

In patient 4, we identified significant X chromosome loss in the blood component of the normal distal sample, particularly in naïve and active T-cell populations. Additionally, a smaller but notable percentage of fibroblast ECM and smooth muscle cells (***F_1_*** and ***F_2_***) also exhibited X-loss. In contrast, Patients 3 and 5 displayed minimal X chromosome loss, accounting for 0.5% of the diploid population.

## Discussion

Single-cell analysis has improved our understanding of biological phenomena. If one interprets this phrase expansively it began with microscopy itself, which subsequently led to the cell-based theory of life. More recently, single-cell molecular sequencing has improved our picture of tissue structures and their pathologies, such as cancer. Here we focus on two types of molecules, RNA and DNA.

Single-cell DNA sequencing can differentiate diploid cells from tumor cells, and among tumor cells identifies distinct sub-clones, each characterized by a unique set of copy number variations. Sampling tumors from multiple sites provides insights into clonal phylogeny and mobility^49^, and clinical outcome^50^. Present-day single-cell DNA analysis can also uncover some somatic variants^51^, such as those marked by the loss of the X-chromosome observed in this study.

Single-cell RNA sequencing generates a fine-grained transcriptomic census of tumor and stroma. This approach generates a low-resolution description of the transcriptional profile of each single-cell, then aggregates these profiles into clusters of common expression patterns. This approach determines the proportions of diverse stromal cell types and cancer expression states, while uncovering gene markers that distinguish these different cell types or states. These gene markers may then be applied to transcriptomic microscopy to obtain spatially resolved information.

Unfortunately, RNA-only data cannot definitively distinguish a stromal expression state from that of the tumor or tumor precursors. Identifying which expression patterns belong to stromal cells and which belong to tumor cells becomes particularly challenging when stromal populations exhibit atypical expression or when tumor cells mimic normal cells. Moreover, there is little hope of connecting tumor genomic changes marked by DNA clonality with changes in expression states marked by RNA clusters.

To close this gap in our understanding requires additional tools such as single-cell DNA and RNA together from the same nuclei, ideally with thousands of nuclei per tissue sample. We achieved this by leveraging our **BAG** (**b**alls of **a**crylamide **g**el) platform. The BAG platform employs a microfluidics oil-emulsion technique to co-encapsulate single nuclei with Acrydite-modified primers and acrylamide monomers into a droplet. The primers hybridize to the cellular nucleic acids, which are subsequently copolymerized to convert the droplet into a permeable gel matrix or **BAG**. Depending on primer design, hybridization conditions, and polymerases, BAGs can capture either DNA or RNA alone, as we have previously shown.

In this paper, we introduce a new method that captures DNA and RNA together in **hybrid BAGs**. We demonstrated that hybrid BAGs provide sequence data that are consistent with those obtained from DNA-only or RNA-only BAGs. To do this, we clustered the single-cell data using DNA-only, RNA-only and the DNA-layer and RNA-layers from the hybrid data. Although the coverage in the hybrid data is lower than for DNA-only or RNA-only, the hybrid data are sufficient to establish the same genomic and expression identities identified by the other two methods. In the five patients we examined, we observed comparable and consistent clusters (**Figure 2, Figure S4-S7**). This includes even subtle features, such as a single 43 MB deletion that distinguished two tumor subclones from Patient 1, and loss of the X chromosome in some diploid cells.

Having established the effectiveness of the hybrid method for each modality, we then explored the interplay between the DNA and RNA layers. We found that the combination of expression and genomic data from hybrid BAGs clarified the sub-population composition of the primary site of tumors. We used several standard tools from the Seurat package, including cell type clustering, subset selection, and UMAP visualizations. We also developed additional modules: (1) using Seurat to cluster the DNA copy number data; (2) alluvial plots to visualize the mappings of genomic clusters into expression clusters; (3) multinomial methods to determine cell cluster membership and to filter collisions; (4) establishing distance metrics relating clusters; and (5) cluster back propagation which uses information from the DNA-layer to divide RNA clusters, and the reverse.

Combining information from the DNA and RNA layers, we first examined the RNA clusters and found that for every cluster, either (1) the nuclei map predominantly to the diploid DNA clusters (**N** or **Nx**), or (2) the nuclei map predominately to the aneuploid DNA cluster(s) of a single patient. We used this information to label some RNA clusters as tumor and others as somatic.

The tumor nuclei from the same patient show common copy number features and distinctive loss-of-heterozygosity patterns in the DNA-layer. These nuclei predominately map to a set of 1-3 RNA expression clusters that are unique to that tumor. By examining the alluvial plots in **Figure 3**, we note varied patterns of mapping between the tumor DNA clones and the RNA expression clusters.

Patient 1, diagnosed with uterine carcinosarcoma, presents a clear example of two distinct DNA clones, **A_1_** and **B_1_**, mapping to the same two expression clusters, ***Ta_1_*** and ***Tb_1_***. The specific 43 MB deletion unique to **B_1_** nuclei suggests a clonal origin for this subpopulation. The RNA expression clusters have notable differences: The ***Ta_1_*** RNA cluster shows increased expression of collagen genes whereas the ***Tb_1_*** RNA cluster shows increased expression of genes related to differentiation and growth signaling pathways. However, both **A_1_** and **B_1_**map equally to the same two expression clusters and with similar proportions, strongly suggesting that the phenotypic flexibility of the tumor was preserved through the single-cell bottleneck event that generated the **B_1_**population.

The tumor of Patient 4 presents a more clinically advanced uterine carcinosarcoma. In the DNA-layer, we identified two distinct tumor clones **A_4_** and **B_4_** distinguished by several copy number changes comprising whole chromosome arms. The **A_4_** subpopulation displayed LoH and copy number losses relative to **B_4_**, suggesting a greater divergence from the initial tumor clone. From the RNA-layer, we identified three expression states: a major ***Ta_4_*** cluster and two much smaller expression clusters: an “active” ***Tb_4_*** cluster marked by increased expression of metabolic genes (*EEF1A1*, *IGFBP5*, *MARCKS*), and a “dividing” ***Tc_4_*** cluster with increased expression of cell cycle genes (*MKI67*, *CENPF*). Both clonal populations map to all three expression states, suggesting that this diversity was preserved during tumor evolution. However, there is a notable difference in their proportionate contributions to the three distinct expression states; namely, the **A_4_** clone contributes more significantly to the “active” and “dividing” states.

Given that both **A_4_** and **B_4_** contribute substantially ***Ta_4_***, we can use a process unique to hybrid data that we call **back-propagation**. In back-propagation, we use the DNA-layer identities to look for additional substructures in the RNA-layer expression states (and vice-versa). In the case of ***Ta_4_***, we split the expression cluster into two groups, ***Ta*_4_-A_4_** and ***Ta*_4_-B_4_**, and then tested those groups for significant differences in expression. We found that the two sub-groups are highly similar, with two genes showing significant deviations that distinguish the lineages.

Patient 2 presents a contrasting story, with tumor cells demonstrating a marked dependency between genomic identity and RNA expression. This tumor tissue sample comprised four sub-clones **A_2_**, **B_2_**, **C_2_** and **D_2_** mapping into two expression clusters ***Ta_2_*** and ***Tb_2_***. The LoH analysis determined that the **A_2_** and **B_2_** sub-clones share a common ancestor that arose after the split of **C_2_** and **D_2_**. The alluvial plot shows that the mapping of the DNA clusters into the RNA-layer recapitulates this split: **A_2_** and **B_2_** map to ***Ta_2_*** and **C_2_** and **D_2_**map to ***Tb_2_***. Differential gene analysis in the RNA-layer suggests that ***Ta_2_*** comprises a more aggressive and stress-resistant population. In this instance, we also utilized back-propagation to test for additional divisions in the two expression clusters. In both cases, we found a subset of unique genes that are differentially expressed in the hybrid RNA-DNA product clusters: ***Ta*_2_-A_2_**, ***Ta*_2_-B_2_**, ***Tb*_2_-C_2_**, and ***Tb*_2_-D_2_**. These genes may have important implications for the evolution of the tumor. They can also serve as lineage-specific expression markers in technologies that are limited to RNA.

In the combined data from all samples, we identified 23 distinct somatic expression clusters. The majority of these clusters (21 of 23) include nuclei from more than one patient, suggesting that many somatic cell types have consistent expression patterns across individuals. In contrast, every tumor cluster occurs in one and only one patient with little in common between the tumor clusters from different patients.

By analyzing highly expressed genes, we accurately determined the likely cell type or state for most of these clusters. Our analysis revealed two stromal expression clusters unique to Patient 1: a fibroblast population in the distal normal tissue with activation of the AP-1 pathway, and an epithelial population in the tumor tissue with unique makers for signaling and response to stimuli. This epithelial population was distinct in both the UMAP and the multinomial tree. Without the DNA-layer from the hybrid data, we could not have confidently concluded that this unique cluster derives from diploid cells.

While most of the diploid cells are copy number 2 everywhere (labeled as **N**), in every patient we identified a subset of diploid nuclei that exhibited the loss of an X chromosome (labeled **Nx**). While this phenomenon has been previously observed in both tumors and somatic populations, the hybrid BAGs provide new insights into their role in tumor biology. Because we obtain DNA and RNA from the same single cells, we can pinpoint which stromal populations are showing the loss the X chromosome. In patient 2, we identified an X-loss event in a significant proportion of the plasma cell and T-cell populations. Using the RNA data and the DNA data together, we determined that each plasma cell lost the same X chromosome, whereas the T-cells lost either one or the other equally. This strongly suggested that the plasma cell population may have a common origin. Comparing the expression profiles of ***plasma*-N** and ***plasma*-Nx**, we observed reduced expression of *XIST* and *TSIX*, in the **Nx** population, further confirming that the X-loss observed in the DNA layer is not a sequencing artifact but a genuine genomic event. Furthermore, the differential expression analysis showed differences the *IGH1* and *IGH2* genes. This led us to explore the VDJ recombination region, providing decisive evidence of a clonal origin for this mutant plasma cell population.

In our analysis of the hybrid data, we observed a small but significant population of cells that appear to violate their identities: tumor cells with normal expression and diploid cells with tumor expression. The former suggests a sort of mimicry in which the cancer cells appear like normal stroma while the latter may point to a pre-aneuploid tumor precursor state or stromal cells dramatically altered by the tumor microenvironment. However, the application of a collision detection method significantly reduced these crossover populations. This reduction has diminished our initial belief in the prevalence of mimicry or progenitor states as observed in the hybrid dataset. The existence of such cells cannot be completely dismissed, and further refinements to the method, larger datasets, and additional data types will be required to determine their frequency and biological significance.

Most tumors exhibit detectable copy number differences^52^. Because of the instability of aneuploid genomes during replication, we can often find unique copy number changes that trace a tumor’s lineage. As we have observed in this study, even somatic cells may carry detectable copy number changes that mark their lineal descent. Unfortunately, some tumors do not have copy number changes and many bifurcations in the tumor phylogeny go unremarked by copy number events; and the vast majority of phylogenic branches of the stroma do not carry a unique, detectable copy difference.

To enhance our ability to identify unique subclones in both the tumor and stroma, we are focusing on smaller genomic changes such as single nucleotide variants (SNVs)^53–55^, small insertions and deletions (indels), and microsatellite length variations (MSLVs). We are developing a protocol using hybrid BAGs to capture both RNA and microsatellites, taking advantage of the fact that BAG allows for sequence-specific capture, as opposed to transposase-based multiomic technologies. With the high mutability of microsatellites and the advent of new, more accurate measurement methods^56^, this strategy promises to provide detailed insights into tumor phylogeny and also identify unique fingerprints for somatic subpopulations.

In this paper, we used the DNA-only and RNA-only datasets to confirm the validity of the hybrid BAG data. From that point forward, our analysis relied solely on the information from the hybrid data alone. However, the other two datasets provide deeper coverage of the DNA and RNA data separately. This suggests that one could use hybrid BAGs primarily as a bridge, mapping between RNA and DNA clusters. In such a workflow, information passes in both directions: the RNA-only and DNA-only data provide improved resolution for clustering the hybrid data while back-propagation in the hybrid dataset informs additional splits in the DNA-only and RNA-only data. This approach could be especially valuable in single-cell transcriptomics spatially resolved by multi-gene-target microscopy.

With advanced platforms like Xenium, MERSCOPE spatially resolving individual transcripts in hundreds of thousands of cells^38,39^, researchers can now custom order several hundred target genes to query their sample. Here, hybrid BAGs could play a pivotal role. From a hybrid BAG experiment, we can determine the set of marker genes that maximally capture the somatic and tumor expression patterns. By using methods like back-propagation, we can identify lineage-specific markers genes that may enable researchers to use these RNA-only transcriptome microscopes to determine spatial information about genomic variation as well. Hybrid BAGs provide a unique method for selecting the optimal palette of genes to spatially resolve both expression and lineage in a tumor sample.

## Methods

### Pulverization of frozen tissue samples in liquid Nitrogen

All patient tissue samples were pulverized in Liquid Nitrogen (LN2) with a sterile mortar and pestle prior to analysis. Mortar and pestles were submerged in LN2 and cooled to LN2 temperature. The cooled vessels were then partially filled with fresh LN2 and transferred to a basin containing a shallow pool of LN2. The presence of LN2 in both the mortar and basin helped maintain a constant temperature during the pulverization process and prevent sample heating due to friction. The tissue samples were then transferred to the sterile mortar, submerged in LN2, and pulverized until they were mostly a fine, homogeneous powder. Once pulverized, residual tissue material was scraped off the pestle back into the mortar with a sterile, LN2-cooled disposable spatula. The mortar was then removed from the basin to allow for the LN2 to evaporate out of the mortar. Subsequently, pulverized tissue was immediately collected with a fresh, sterile, LN2-cooled disposable spatula into 2.0 mL DNA LoBind Eppendorf tubes submerged in LN2. Pulverized samples were placed on dry ice with the caps open to allow for temperature equilibration before closing the tubes, and then stored at -80°C until further use. All samples were pulverized with separate sterile mortar and pestles to avoid cross-contamination between tumor and normal tissues.

### The sample cohort

We studied samples from five patients (patient 1 – patient 5). The samples were from their uterine cancers (Tumor 1 – Tumor 5), and in three patients also from an adjacent normal endometrial sites (Normal 1, Normal 2, and Normal 4). For most samples (Normal 1, Tumor 1, Tumor 2, Tumor 3, Normal 4, Tumor 4, Tumor 5), we sequenced single nuclei of the same sample on each of three platforms: DNA-only, RNA-only, and the hybrid protocol. We performed comparison analyses and showed the validity of the hybrid protocol mainly using the above five trio data sets.

### Hybrid BAG generation

We dissolved the pulverized tissue in ice-cold NST detergent buffer ^49^ and stained with DAPI. We performed single-nuclei sorting using DAPI-H vs. DAPI-A single-nuclei gate on a FACSAria II SORP cell sorter to remove debris and clumps. We confirmed (data not shown) that single-nuclei sorting based on ploidy would not be able to distinguish cancer cells from normal cells because the hypodiploid peak of cancer cells often overlaps with the diploid peak of normal cells ^49^. Single nuclei were loaded into the microfluidic device described in detail in a previous publication ^33^. Nuclei were encapsulated into droplets with an average diameter of 120 microns. For the capture of nucleic acids, we used 5’ Acrydite oligonucleotides. All the Acrydite-modified oligonucleotides became covalently co-polymerized into the gel ball matrix. They also all contained, at their 5’ end, a universal PCR primer (UP1) for subsequent amplification. For RNA-only protocol, we used oligo-dT; for DNA-only protocol, we used random T/G primers, and followed their respective published protocols ^33^. To capture both RNA and DNA together in the new hybrid protocol, we used both Acrydite primer designs, but we altered the protocol in two important ways.

The first critical change was an incubation step at 85°C for 5 minutes instead of 95°C for 12 minutes for DNA denaturation in the DNA-only protocol. Otherwise, we observed significant destruction of the RNA.

The second critical change took place after the BAGs were formed. The RNA and genomic DNA trapped in the BAGs were used as templates to make covalently bound copies, and in the new hybrid protocol, both reverse transcriptase and DNA polymerase were used. Template-switch-oligos were also introduced in the hybrid protocol so that the cDNA products which were covalently linked to the BAG matrix ended with a double-stranded region. This double-stranded DNA region included an NLA-III cleavage site. Subsequently, DNA polymerase (Klenow) was added to extend the captured genomic DNA from primers, forming a copy that was also covalently linked to the BAGs. Some, perhaps most, of the cDNA-mRNA sequence was further partially converted to double-stranded cDNA. BAGs were pooled and the covalently captured DNA and cDNA were cleaved with NLAIII leaving a sticky end used for subsequent extensions.

BAG barcodes and varietal tags were added to the 3’ ends of the covalently captured nucleic acids in split-and-pool reactions. The BAG barcodes were present on both the genomic-DNA and RNA copies. The varietal tags were used for counting. The first BAG barcode and varietal tag were added by ligation extension (described in detail in the supplementary experimental protocol), leaving a common 3’ sequence identical across all the molecules and BAGs. The second BAG barcode and varietal tag were added by hybridization extension of the common 3’ sequence, along with a second common sequence adapter for the third split-and-pool step. The third barcode was added by a split PCR, using the first universal PCR primer (UP1) and the second common sequence adapter as part of the PCR primer sequences.

These amplified products were pooled and converted by tagmentation into paired-end Illumina sequencing libraries. One end of the reads contained BAG barcode and varietal tag, as well as genomic or transcriptomic sequence information. The other end from random tagmentation was mostly genomic or transcriptomic sequence information.

### Initial data processing

Sequencing libraries were sequenced in paired-end 150 bp format using an Illumina NovaSeq 6000. Briefly, each processing step is described in more detail in the immediately following sections. We first checked the structure of each read pair in the fastq files. For the good read pairs with the correct structure as shown in Figure 1C, we extracted the BAG barcode, varietal tag, and genomic sequences from both reads. We then mapped the genomic (including transcriptomic) sequence to the reference genome with gene transcript information. Finally, we combined the mapping information from all reads belonging to each varietal tag for each BAG barcode. In the end, we obtained a template data table with each row containing the information of an original template/molecule. In the following section, we explain each processing step from the fastq file to the template table in detail.

#### Step 1 – Check sequence structure

First, we filtered out reads from the fastq files where either Read 1 or Read 2 were less than 100 bases. Second, we examined if the sequences from the expected BAG barcode positions exactly matched one of the 96×96×96 barcodes, and if the "CATG" cutting site was in the expected location, allowing for one base mismatch. We removed read pairs that did not satisfy these requirements. Third, from Read 1 which started with barcodes and varietal tags, we trimmed away the first 80 bases containing the BAG barcode, varietal tag, and adapter sequences, and also checked if the reverse complementary sequence of the universal primer ("CCAAACACACCCAA") or oligo-dT ("AAAAAAAAAAAAAA") was present. If present, it meant we had reached the end of the template, so these primer-related sequences were trimmed off for downstream mapping. Similarly, for Read 2, the tagmentation end, we checked and removed the adapter sequence ("GAGCGGACTCTGCG") from the first split-and-pool if it existed. After trimming, we required both Read 1 and Read 2 to be at least 30 bases long. All the bases from Read 1 and Read 2 after trimming were then used for paired-end mapping (Step 3).

#### Step 2 – Extract BAG barcode and varietal tags

If a read pair passed Step 1, we extracted the BAG barcode and varietal tag information from the first 80 bases of Read 1, and this information was appended to the read ID. The 17 base BAG barcodes came from three cycles of the split-and-pool procedure, of which five bases came from the 1^st^-split, six bases came from the 2^nd^-split, and six bases came from the 3^rd^-split. There were 96 different barcodes for each split, so there were altogether 96×96×96 (≈ 1 million) varieties. The 12 base varietal tag came from both the split-and-pool primers and the genomic sequence. Out of these twelve bases, four bases came from the 1^st^-split, four bases came from the 2^nd^-split, and four bases came from the genomic sequence that was two bases away from the "CATG" cutting site. These twelve bases provide 4^12^ (≈ 16 million) varieties for each BAG.

#### Step 3 – Map to the human genome

After steps 1 and 2 above read pairs were mapped to the UCSC hg19 human genome using HISAT2 version 2.1.0 ^57^. The reference genome we used included the primary chromosomes and unlocalized and unplaced contigs. Alternate haplotypes were not included in the genome index. HISAT2 can take a file with known splice sites to use for alignment. This file was generated using a gtf formatted file extracted from the NCBI refSeq gene annotation table from the UCSC genome browser and the HISAT2 program, hisat2_extract_splice_sites.py. The bam files were then sorted and indexes using samtools. In subsequent data analysis steps we designate by mapped reads the reads that HISAT2 marks as being part of a proper pair and a primary mapping having a read mapping quality score greater than zero.

#### Step 4 – Combine read information with original template information

We grouped the mapped reads based on their BAG barcodes. For the reads with the same BAG barcode, we sorted the varietal tags by the number of reads associated with each tag in descending order. We performed a "rollup" algorithm on the sorted varietal tags, and discarded varietal tags within a Hamming distance of one from a more abundant varietal tag having at least ten times more reads. We assumed the eliminated varietal tags originated from the tags with more abundant reads but contained sequencing or PCR errors.

Using the varietal tags from the above "rollup" step, we aggregated the mapped segments for all the reads with the same varietal tag. We checked the total coverage of each varietal tag against all exons and transcript boundaries from the NCBI refSeq gene annotation file downloaded from the UCSC genome browser, and wrote out one line per varietal tag with all the useful information into a "template table". Each line of the template table contains the following information: BAG barcode, varietal tag, chromosome, start mapping position, end mapping position, start and end mapping position for each fragment if there was more than one continuous fragments, total bases covered by this template, number of reads, number of genes, gene list, bases overlapping with the transcript of the best-matched genes, bases overlapping with exons, number of splice junctions, number of unspliced sites, bases overlapping with the coding regions, 5’UTR, and 3’UTR of the gene. The downstream data analyses were mainly based on the information from this table. The best-mapped gene was deemed to be the gene from the annotated transcript file having the highest overlap to the transcript. If more than one transcript had the same overlap then best was determined by overlap to exons, then overlap to coding sequence, then the number of splice junctions, then the fewest unspliced sites. If more than one gene tied for all these criteria, then all genes are listed in the template table.

### Template processing

#### Sequence classification

Starting from the template data table described above, each initial molecule was classified as one of the four categories: "Exonic"," Intronic", "Intergenic", and "Uncategorized". This process was applied uniformly regardless of the protocol types (RNA-only, DNA-only, or hybrid). We classified a template as an "Exonic" template if over 90% of its bases were mapped within one gene. Furthermore, we refined the "Exonic" classification only if 50% or more covered bases from this template were exonic, or if 20% or more covered bases were exonic and at least one splicing event was observed (RNA layer). If all the bases from a template were mapped to intergenic regions, we classified it as "Intergenic". If a template was not classified as "Intergenic", but less than 10% of its covered bases were exonic and no splicing events were observed, this template was classified as "Intronic". Only a small proportion of templates failed to be classified into the above three categories, and these templates were classified as "Uncategorized".

For expression clustering, we only used "Exonic" templates assigned to a single gene regardless of protocols. For copy number clustering, we tested four versions of template choices on all the libraries, which we will discuss in the next section.

#### Copy number plot varieties

We demonstrated four progressively improved versions of copy number estimation, named "all_molecules", "no_exon", "no_gene", and "no_gene.avoid50closeTN". The method "all_molecules" simply used all molecules for each retained nucleus for copy number as the name would imply. The method "no_exon" used molecules both classified as "Intronic" and "Intergenic" in the previous paragraph. The method "no_gene" only used "Intergenic" templates with no bases covering a transcript.

The method "no_gene.avoid50closeTN" (DNA layer) only retained the "Intergenic" molecules from the "no_gene" method that were at least 50 bases distant from RNA hotspots. We defined an RNA hotspot as the genomic region between two "Intergenic" templates that were within 50 bases of each other in RNA-only libraries from all tumor and normal samples in the cohort. RNA hotspots were expected to be some combination of actual unannotated transcripts and regions of DNA that were prone to being copied by reverse transcriptase. As these hotspot sequences distorted copy number profiles in normal and tumor tissue specimens, we eliminated certain intergenic regions when determining copy number profiles for the hybrid protocol.

#### Empirical bin boundary generation for copy number

Separately for each of the four copy number molecule selection methods above, we used the genomic positions of all molecules from normal DNA samples to determine empirical bin boundaries for 300 bins with approximately equal molecule counts per bin. Excluding any molecules mapping to chromosome Y, we assigned to each chromosome 1-22 and X a number of bins in proportion to its fraction of total molecule counts. Within each chromosome, bin boundaries were assigned greedily from the start of the chromosome so that all but the final bin contained at least the same number of molecules that was equal to the total counts of molecules (or referred to as templates) divided by the number of bins for that chromosome. The observed count of molecules per bin was recorded as a normalization factor for later use during per-sample copy number estimation. This normalization factor could vary by up to 30% between chromosomes because a small number of bins (300) can only be imperfectly allocated by chromosomal molecule counts.

#### Copy number estimation

For each copy number variant separately, each selected molecule incremented a bin based on the established bin boundaries for that method. Each bin count was then divided by the per-bin normalization factor, and the result was multiplied by 2 divided by the median value over all bins. Assuming a mostly diploid sample, this process resulted in a copy number profile for the sample that was centered at a value of 2. Circular binary segmentation (DNAcopy version 1.50.1) ^58^ in R was then performed on the copy number profile using parameters alpha=0.02, nperm=1000, undo.SD=0.5 and min.width=2. For each profile, we also computed a quantity we call ’terrain’ which was the sum of the absolute value of adjacent bin copy number differences. To produce copy number input for the "CreateSeuratObject" function of Seurat, the per-bin normalization factor was applied to each raw bin count for each cell, and a second per-cell normalization factor was then used so that each cell’s total normalized count was set equal to its total unnormalized count.

### RNA clustering

The RNA clustering was performed using Seurat package (version 3.1.5) following the standard Seurat clustering pipeline^41^. The gene names were also appended with the chromosome information to distinguish any ambiguous locations. We removed the ribosomal protein genes for clustering. For comparing expression clustering between the hybrid protocol and RNA-only protocol, we normalized the gene-template matrix by cell and excluded the PCA components that most significantly distinguished protocol differences. We typically used at least 15 PCA components for clustering. This approach gave us similar clustering results as the "IntegrateData" function in Seurat v4. For the combined RNA clustering of all the hybrid data, we downsampled the gene matrix to 400 Exonic templates per nucleus, and included nuclei with more than 300 Exonic templates for clustering. In the clustering process, we only used genes that showed up in at least 30 nuclei, and nuclei with at least 150 genes; we used the top 5,000 variable gene features for PCA analysis and used the first 50 PCA components for subsequent UMAP and FindCluster functions. The detailed Seurat parameters and R code have been uploaded to GitHub and can be found at: https://github.com/siranli01/DNA_RNA.

### RNAScope imaging analysis

RNAScope Multiplex Fluorescent Reagent Kit v2 (Advanced Cell Diagnostics) was employed to visualize RNA transcripts within tissue sections. Formalin-fixed, paraffin-embedded (FFPE) tissue slides from Tumor 2 were prepared according to the manufacturer’s instructions. Following deparaffinization and rehydration, sections underwent hydrogen peroxide treatment and target retrieval. Protease treatment was then applied to facilitate probe penetration.

The following RNAScope probes were used for target detection: IGHG-pool (Cat No. 481901), which targets IGHG (1-4) with 11-19 ZZ pairs, and human XIST (Cat No. 311231-C2), which targets the XIST RNA transcript. Signal amplification was performed using the RNAScope Multiplex Fluorescent Reagent Kit v2. The TSA Vivid Dyes were used for fluorescent signal development: IGHG was visualized using TSA Vivid Fluorophore 520 dye (Cat No. 323271), and XIST was visualized using TSA Vivid Fluorophore 570 dye (Cat No. 323272). Amplification steps followed the standard protocol provided by Advanced Cell Diagnostics, ensuring optimal signal-to-noise ratio.

Imaging was conducted on a Nikon Ti spinning-disk confocal microscope equipped with a YOKOGAWA spinning-disk system and controlled by Nikon Elements software AR 5.42.04. Fluorophores were excited with the following laser lines: DAPI at 405 nm, IGHG (TSA Vivid Fluorophore 520) at 488 nm, and XIST (TSA Vivid Fluorophore 570) at 561 nm. Fluorescent signals for DAPI, IGHG, and XIST were pseudo-colored for visualization.

### Copy number clustering

Similar to RNA clustering, we used "RunUMAP" and "FindClusters" functions of Seurat to cluster nuclei based on copy number. For each library, we had a bin-counts matrix, similar to the gene matrix for RNA clustering. There were 300 rows in the matrix, representing 300 genomic bins. Each column represented a nucleus. Each element of the 2D matrix represented the tag counts of the corresponding bin in the corresponding nucleus. We first normalized the matrix by columns: for each nucleus, we divided each bin count by the mean of 300 bins and then multiplied by 2. We not only used these 300 normalized single bin counts for clustering; additionally, we also included the median normalized bin counts of every two and three adjacent bins, as long as these adjacent bins were within the same chromosome. The reason for this step was that copy number segmentation usually requires similar amplification or deletion patterns in at least two contiguous bins. By doing this, we appended another 277 rows from the two adjacent bins and 254 rows from the three adjacent bins onto the original 300-row normalized bin-count matrix.

We performed clustering using the new matrix with 831 rows. We used a workflow similar to that for RNA clustering, but we did not use "NormalizedData" function since the matrix had already been normalized. For “FindVariableFeatures” function, we used the top 500 features by inputting “selection.method = “vst”, nfeatures = 500”.

### Copy number heatmap

The single-nucleus copy-number heatmap was plotted using Seurat "DoHeatmap" function. Each row represented the median normalized counts of two adjacent bins, except for the first bin of each chromosome, in which we used the normalized count of that single bin. The total of 300 rows were sorted in genomic order, with chromosome Y eliminated.

### Multinomial distributions

For a cluster X, we sum the RNA template counts over each gene for all cells in the cluster. If a gene contains zero counts over the population of cells, we assign a value of ½, to avoid zero probabilities when comparing to a cell that containing a gene unobserved in cluster X. We normalize by the count vector by its total to obtain a probability distribution over the set of genes. We compute this probability vector for each somatic expression cluster as determined by the Seurat iterative clustering of the somatic cell types, for each tumor cluster as determined in the individual tumor RNA clusters, and for an “empty” cluster composed of the RNA-layer from 500 DNA-only BAGs. The expression multinomial distributions are used for two purposes:

1. **Cluster assignment.** We assign each nucleus to the distribution with the maximum likelihood of generating its observed counts, assuming a uniform prior on the space of clusters. These identities determine the color of the points in Figure 4A and the counts in Table 1.
2. **Inter-cluster dissimilarity**. We compute a pairwise dissimilarity between clusters using the multinomial distributions induced by the cluster average. For any two clusters **X** and **Y**, and template count **t** between 1 and 300, we compute by simulation (100,000 samples per data point) the posterior probability that a single nucleus generated from the multinomial distribution on **X** and a count of **t** templates comes from **X**, conditioned on the prior probability of originating from **X** or **Y** with equal probability. For each value of **t**, we compute the AUROC (equivalent to the average posterior probability). To define a measure of how likely **X** is correctly identified against **Y**, we compute **M(X, Y)**, the smallest value of **t** for which the AUROC exceeds 0.999 (Figure S15). To convert this matrix into a pairwise dissimilarity between clusters, we compute a similarity measure **S(X, Y)** = ½ * [**M(X, Y)** + **M(Y,X)**] and invert this to obtain a dissimilarity measure **D(X, Y)** = 300 – **S(X, Y)**. We then apply the neighbor-joining algorithm to the dissimilarity measure to obtain a tree (Figure 4B).

### Multinomial Wheel

To build a multinomial wheel in DNA space, we first computed a multinomial vector to represent each Seurat cluster. Each multinomial vector had 300 elements, representing 300 genomic bins. Each element was the total bin counts from all the nuclei in that cluster. We normalized each vector to sum to one, serving as the multinomial probability vector representing that cluster. Next, we computed the linear combination of multinomial probability vectors of every two Seurat clusters, and created 9 equally spaced sampling states *C*_1,2,…,9_ = *pA* + (1 − *p*)*B*, for *p* = (0.1, 0.2,…, 0.9), where A and B are the two original states. We then assigned the nucleus to the state with the highest likelihood. In R language, we used the "dmultinom" function to compute multinomial probabilities.

We applied a similar idea to create the RNA multinomial wheel. Different from the DNA multinomial vector where each element was a genomic bin, in RNA space, each element represented one of the 29,637 genes. We computed the sum of gene counts for each Seurat cluster *V*_1,2,…,*n*_ (n is the number of Seurat clusters, and *V*_*i*_ is a 29,637-element vector, *i* = 1,2, …, n), but unlike DNA, there were many elements still being zero which could not be used as a multinomial probability vector. We solved the problem by adding a small value to each element that was proportional to the total expression level of every gene, so that each vector *V*^∗^does not contain zero elements. For each gene element j, we did the following transformation: 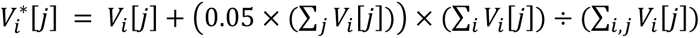. We then normalized each vector 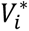 to obtain the multinomial probability vector for cluster i.

## Supporting information

Supplementary Protocol

## Acknowledgments

We thank P. Moody for cell sorting; J. Preall, C. Regan, E. Zhang in CSHL single-cell core facility for assistance; Q. Gao for histology assistance; A. Runnels from NYGC and E. Ghiban from CSHL for Illumina sequencing assistance; M. Yao, D. Fearon, T. Janowitz, D. Tuveson from CSHL and N. Chiorazzi, M. Frimer from Northwell for helpful discussion, A. Kapedani, M. A. Green, and S. Chin from the Clinical Research Team in the Department of Obstetrics and Gynecology of Northwell Long Island Jewish Medical Center for clinical information assistance. Additionally, we extend our gratitude to S. S. Fox, M. Heywood, K. Quinn, B. M. Weil, J. Jacob from Biobanking/Anatomic Pathology Team in the Northwell Health Biospecimen Repository (NHBR) in Northwell Health Cancer Institute for sample transfer and pathological information assistance. This work was supported by a grant from the Simons Foundation, Life Sciences Founders Directed Giving-Research (award number 519054 to M.W.); The Breast Cancer Research Foundation (BCRF, to M.W.); and by support from the Cold Spring Harbor Laboratory and Northwell Health Affiliation (awarded to M.W.).

## Data availability

Illumina sequencing data for all the single-nucleus libraries are available at NCBI Sequencing Read Archive (SRA) with accession code (PRJNA773107).

## Code availability

Code is available through https://github.com/siranli01/DNA_RNA.

## References

1. Lan, F., Demaree, B., Ahmed, N. & Abate, A.R. Single-cell genome sequencing at ultra-high-throughput with microfluidic droplet barcoding. Nature biotechnology 35, 640–646 (2017).

2. Vitak, S.A. et al. Sequencing thousands of single-cell genomes with combinatorial indexing. Nature methods 14, 302–308 (2017).

3. Andor, N. et al. Joint single cell DNA-Seq and RNA-Seq of cancer reveals subclonal signatures of genomic instability and gene expression. Biorxiv, 445932 (2018).

4. Macosko, E.Z. et al. Highly parallel genome-wide expression profiling of individual cells using nanoliter droplets. Cell 161, 1202–1214 (2015).

5. Klein, A.M. et al. Droplet barcoding for single-cell transcriptomics applied to embryonic stem cells. Cell 161, 1187–1201 (2015).

6. Gierahn, T.M. et al. Seq-Well: portable, low-cost RNA sequencing of single cells at high throughput. Nature methods 14, 395–398 (2017).

7. Cao, J. et al. Comprehensive single-cell transcriptional profiling of a multicellular organism. Science 357, 661–667 (2017).

8. Rosenberg, A.B. et al. Single-cell profiling of the developing mouse brain and spinal cord with split-pool barcoding. Science 360, 176–182 (2018).

9. Habib, N. et al. Div-Seq: Single-nucleus RNA-Seq reveals dynamics of rare adult newborn neurons. Science 353, 925–928 (2016).

10. Gao, R. et al. Nanogrid single-nucleus RNA sequencing reveals phenotypic diversity in breast cancer. Nature communications 8, 1–12 (2017).

11. Zheng, G.X. et al. Massively parallel digital transcriptional profiling of single cells. Nature communications 8, 1–12 (2017).

12. Dey, S.S., Kester, L., Spanjaard, B., Bienko, M. & Van Oudenaarden, A. Integrated genome and transcriptome sequencing of the same cell. Nature biotechnology 33, 285–289 (2015).

13. Macaulay, I.C. et al. G&T-seq: parallel sequencing of single-cell genomes and transcriptomes. Nature methods 12, 519–522 (2015).

14. Han, K.Y. et al. SIDR: simultaneous isolation and parallel sequencing of genomic DNA and total RNA from single cells. Genome research 28, 75–87 (2018).

15. Hou, Y. et al. Single-cell triple omics sequencing reveals genetic, epigenetic, and transcriptomic heterogeneity in hepatocellular carcinomas. Cell research 26, 304–319 (2016).

16. Bian, S. et al. Single-cell multiomics sequencing and analyses of human colorectal cancer. Science 362, 1060–1063 (2018).

17. Han, L. et al. Co-detection and sequencing of genes and transcripts from the same single cells facilitated by a microfluidics platform. Scientific reports 4, 1–9 (2014).

18. Van Strijp, D. et al. Complete sequence-based pathway analysis by differential on-chip DNA and RNA extraction from a single cell. Scientific reports 7, 1–9 (2017).

19. Kong, S.L. et al. Concurrent single-cell RNA and targeted DNA sequencing on an automated platform for comeasurement of genomic and transcriptomic signatures. Clinical chemistry 65, 272–281 (2019).

20. Cheow, L.F. et al. Single-cell multimodal profiling reveals cellular epigenetic heterogeneity. Nature methods 13, 833–836 (2016).

21. Rodriguez-Meira, A. et al. Unravelling intratumoral heterogeneity through high-sensitivity single-cell mutational analysis and parallel RNA sequencing. Molecular cell 73, 1292–1305.e8 (2019).

22. Li, W., Calder, R.B., Mar, J.C. & Vijg, J. Single-cell transcriptogenomics reveals transcriptional exclusion of ENU-mutated alleles. Mutation Research/Fundamental and Molecular Mechanisms of Mutagenesis 772, 55–62 (2015).

23. Yu, L. et al. scONE-seq: A single-cell multi-omics method enables simultaneous dissection of phenotype and genotype heterogeneity from frozen tumors. Science Advances 9, eabp8901 (2023).

24. Vandereyken, K., Sifrim, A., Thienpont, B. & Voet, T. Methods and applications for single-cell and spatial multi-omics. Nature Reviews Genetics 24, 494–515 (2023).

25. Gao, R. et al. Delineating copy number and clonal substructure in human tumors from single-cell transcriptomes. Nature biotechnology 39, 599–608 (2021).

26. Fan, J. et al. Linking transcriptional and genetic tumor heterogeneity through allele analysis of single-cell RNA-seq data. Genome research 28, 1217–1227 (2018).

27. Elyanow, R., Zeira, R., Land, M. & Raphael, B.J. STARCH: Copy number and clone inference from spatial transcriptomics data. Physical Biology 18, 035001 (2021).

28. Patel, A.P. et al. Single-cell RNA-seq highlights intratumoral heterogeneity in primary glioblastoma. Science 344, 1396–1401 (2014).

29. Erickson, A. et al. Spatially resolved clonal copy number alterations in benign and malignant tissue. Nature 608, 360–367 (2022).

30. Yin, Y. et al. High-throughput single-cell sequencing with linear amplification. Molecular cell 76, 676–690.e10 (2019).

31. Olsen, T.R. et al. Scalable co-sequencing of RNA and DNA from individual nuclei. bioRxiv, 2023.02. 09.527940 (2023).

32. Lazareva, O. et al. HIPSD&R-seq enables scalable genomic copy number and transcriptome profiling. bioRxiv, 2023.10. 09.561487 (2023).

33. Li, S. et al. Copolymerization of single-cell nucleic acids into balls of acrylamide gel. Genome research 30, 49–61 (2020).

34. Van der Maaten, L. & Hinton, G. Visualizing data using t-SNE. Journal of machine learning research 9(2008).

35. McInnes, L., Healy, J. & Melville, J. Umap: Uniform manifold approximation and projection for dimension reduction. arXiv preprint arXiv:1802.03426 (2018).

36. Becht, E. et al. Dimensionality reduction for visualizing single-cell data using UMAP. Nature biotechnology 37, 38–44 (2019).

37. Townes, F.W., Hicks, S.C., Aryee, M.J. & Irizarry, R.A. Feature selection and dimension reduction for single-cell RNA-Seq based on a multinomial model. Genome biology 20, 1–16 (2019).

38. Chen, K.H., Boettiger, A.N., Moffitt, J.R., Wang, S. & Zhuang, X. Spatially resolved, highly multiplexed RNA profiling in single cells. Science 348, aaa6090 (2015).

39. Janesick, A. et al. High resolution mapping of the tumor microenvironment using integrated single-cell, spatial and in situ analysis. Nature Communications 14, 8353 (2023).

40. Navin, N. et al. Tumour evolution inferred by single-cell sequencing. Nature 472, 90–94 (2011).

41. Stuart, T. et al. Comprehensive integration of single-cell data. Cell 177, 1888–1902.e21 (2019).

42. Xi, N.M. & Li, J.J. Benchmarking computational doublet-detection methods for single-cell RNA sequencing data. Cell systems 12, 176–194.e6 (2021).

43. Spatz, A., Borg, C. & Feunteun, J. X-chromosome genetics and human cancer. Nature Reviews Cancer 4, 617–629 (2004).

44. Zhou, Y. et al. Single-cell multiomics sequencing reveals prevalent genomic alterations in tumor stromal cells of human colorectal cancer. Cancer cell 38, 818–828.e5 (2020).

45. Panning, B., Dausman, J. & Jaenisch, R. X chromosome inactivation is mediated by Xist RNA stabilization. Cell 90, 907–916 (1997).

46. Weakley, S.M., Wang, H., Yao, Q. & Chen, C. Expression and function of a large non-coding RNA gene XIST in human cancer. World journal of surgery 35, 1751–1756 (2011).

47. Huang, K.-C. et al. Relationship of XIST expression and responses of ovarian cancer to chemotherapy. Molecular cancer therapeutics 1, 769–776 (2002).

48. Bolotin, D.A. et al. MiXCR: software for comprehensive adaptive immunity profiling. Nature methods 12, 380–381 (2015).

49. Navin, N. et al. Tumour evolution inferred by single-cell sequencing. Nature 472, 90 (2011).

50. Alexander, J. et al. Utility of single-cell genomics in diagnostic evaluation of prostate cancer. Cancer research 78, 348–358 (2018).

51. Nam, A.S., Chaligne, R. & Landau, D.A. Integrating genetic and non-genetic determinants of cancer evolution by single-cell multi-omics. Nature Reviews Genetics 22, 3–18 (2021).

52. Krasnitz, A., Kendall, J., Alexander, J., Levy, D. & Wigler, M. Early detection of cancer in blood using single-cell analysis: a proposal. Trends in molecular medicine 23, 594–603 (2017).

53. Nam, A.S. et al. Somatic mutations and cell identity linked by Genotyping of Transcriptomes. Nature 571, 355–360 (2019).

54. Penter, L. et al. Integrative genotyping of cancer and immune phenotypes by long-read sequencing. Nature Communications 15, 32 (2024).

55. Lareau, C.A. et al. Massively parallel single-cell mitochondrial DNA genotyping and chromatin profiling. Nature biotechnology 39, 451–461 (2021).

56. Wang, Z., Moffitt, A.B., Andrews, P., Wigler, M. & Levy, D. Accurate measurement of microsatellite length by disrupting its tandem repeat structure. Nucleic Acids Research 50, e116–e116 (2022).

57. Kim, D., Paggi, J.M., Park, C., Bennett, C. & Salzberg, S.L. Graph-based genome alignment and genotyping with HISAT2 and HISAT-genotype. Nature biotechnology 37, 907–915 (2019).

58. Olshen, A.B., Venkatraman, E., Lucito, R. & Wigler, M. Circular binary segmentation for the analysis of array-based DNA copy number data. Biostatistics 5, 557–572 (2004).

