## Supplementary Figures for "DNA and RNA from the same single nucleus reveals interactions between genomic and transcriptomic landscapes in human tumor samples"

### Figure S1

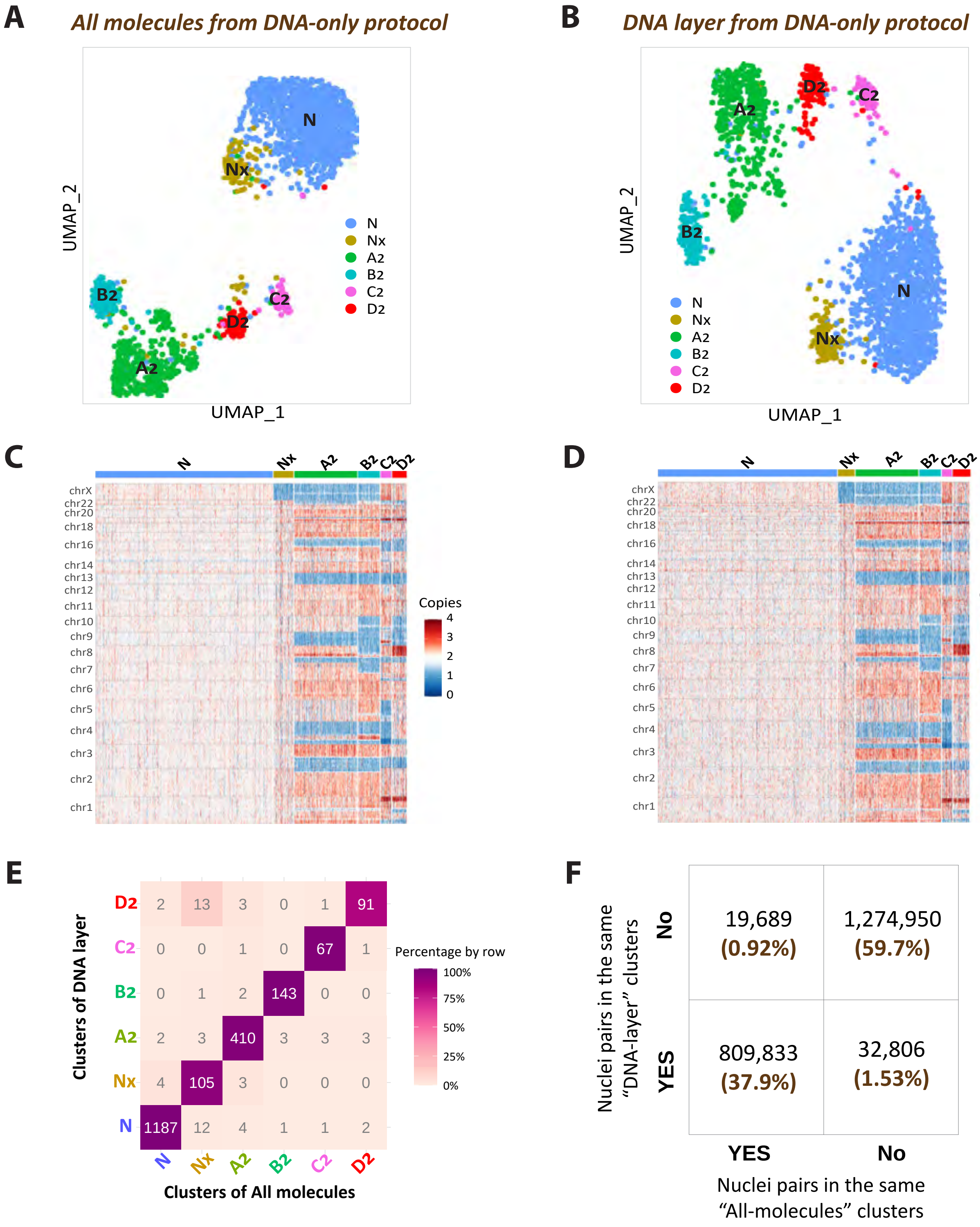


| | N | Nx | A2 | B2 | C2 | D2 |
|---|---|---|---|---|---|---|
| D2 | 2 | 13 | 3 | 0 | 1 | 91 |
| C2 | 0 | 0 | 1 | 0 | 67 | 1 |
| B2 | 0 | 1 | 2 | 143 | 0 | 0 |
| A2 | 2 | 3 | 410 | 3 | 3 | 3 |
| Nx | 4 | 105 | 3 | 0 | 0 | 0 |
| N | 1187 | 12 | 4 | 1 | 1 | 2 |

**Figure S1. Clustering outcomes from the DNA-only protocol using all molecules versus the DNA layer.**
**A**: Genomic clustering results based on bin counts from the DNA-only protocol of Tumor 2, using all molecules. **B**: Genomic clustering results using only the DNA layer molecules from the same group of nuclei as in (A). **C-D**: Copy-number heatmaps for all molecules (C) and only DNA-layer molecules (D), both exhibiting similar copy number patterns. **E**: The number of nuclei in each cluster when including all molecules (columns) versus just the DNA layer (rows), highlighting cluster retention. **F**: Illustration of whether nuclei pairs fall into the same cluster or not across the all-molecules clustering (columns) versus DNA-layer clustering (rows), providing a direct comparison of clustering consistency.

### Figure S2

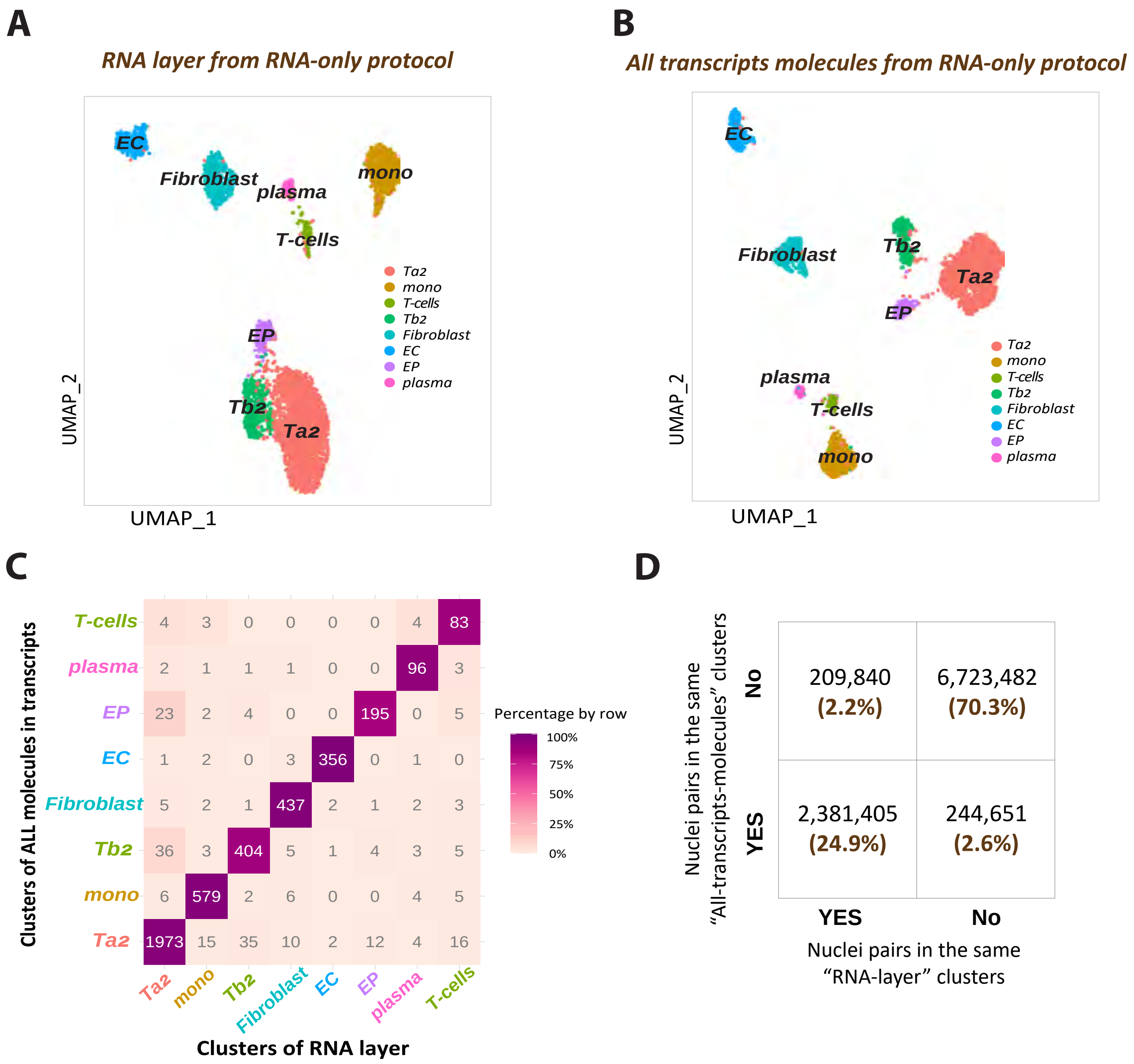


**Figure S2. Comparative RNA clustering using RNA layer and all transcript molecules from an RNA-only protocol.** **A**: UMAP visualization of RNA clustering based on only RNA-layer molecules from an RNA-only experiment in Tumor 2. **B**: UMAP visualization of RNA clustering using all molecules mapped within transcripts from the same set of nuclei as in Panel A. **C**: Comparison matrix showing the number of nuclei in each cluster identified using all-transcript molecules (columns) versus RNA-layer molecules (rows). **D**: A contingency table indicating whether nuclei pairs are grouped in the same cluster in the all-transcripts-molecules clustering (columns) compared to the RNA-layer-only clustering (rows).

### Figure S3

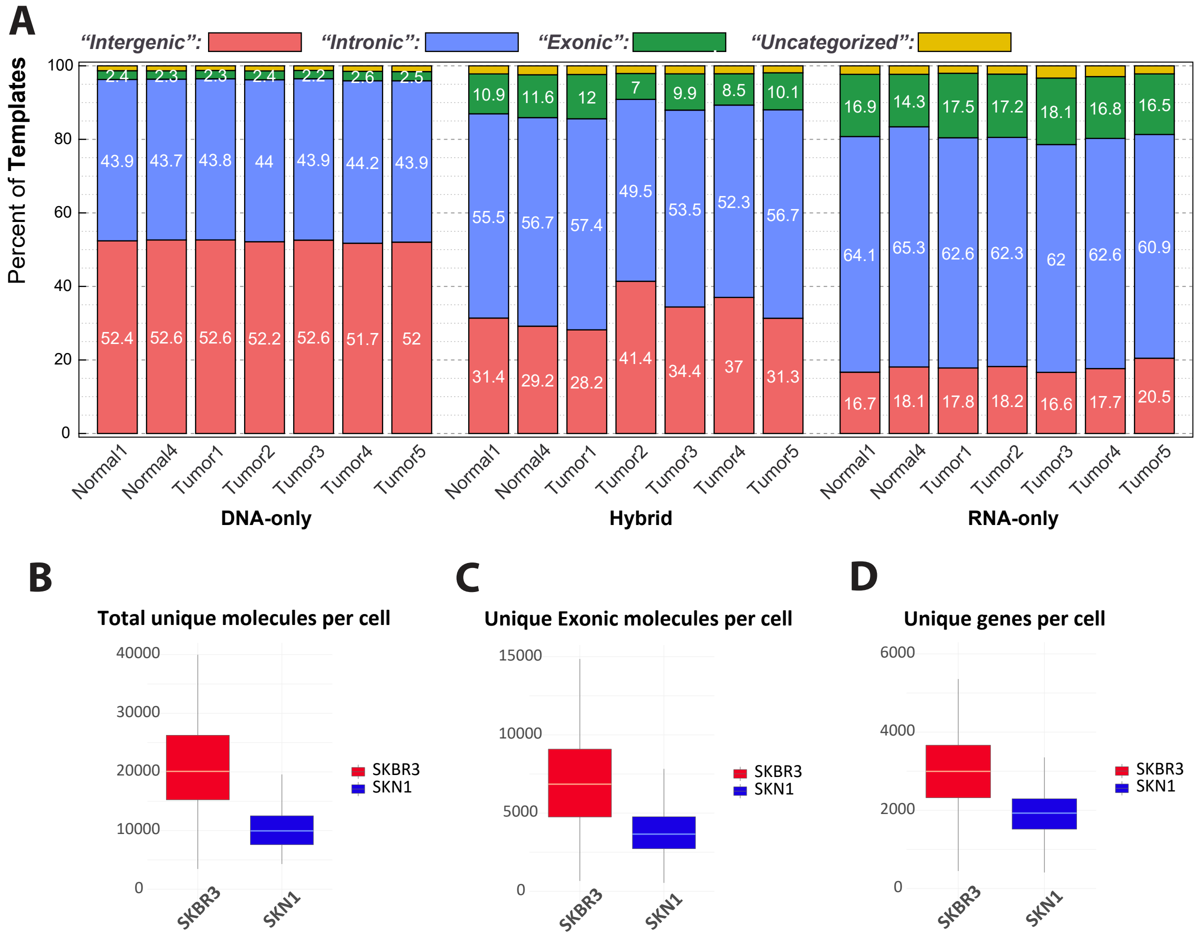


**Figure S3. The template distribution of all the nuclei samples and basic QC of the SKN1-SKBR3 cell-mixture experiment. A**. Distribution of templates for each tissue sample under each protocol, categorized into four categories: intergenic, intronic, exonic, and uncategorized. **B-D.** Total unique molecules (B), unique exonic molecules (C), and unique genes per cell (D) of a cell-mixture experiment using the hybrid BAG-seq protocol.

#### Figure S4

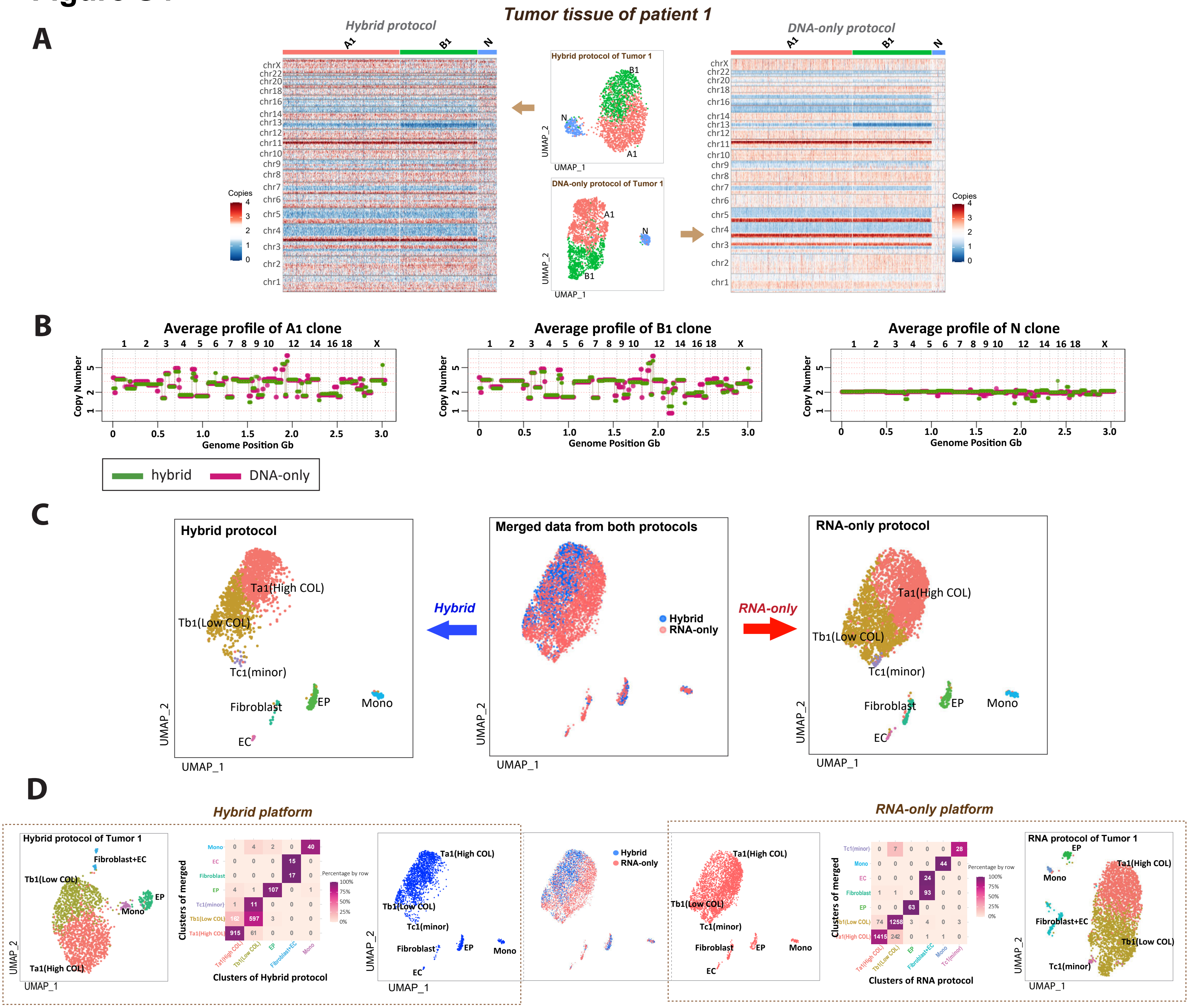


**Figure S4. Comparison of DNA and RNA clustering results between hybrid protocol and DNA-only/RNA-only protocols for Patient 1.** Following the structure of Figure 2, this figure provides a comparison of the hybrid protocol data to the DNA-only and RNA-only protocol datasets for the tumor sample of patient 1. **A**: Single-cell DNA copy number analysis of the tumor sample comparing the hybrid protocol data (left) with the DNA-only protocol data (right). The tumor sample has three distinct copy number profiles: normal diploid cells (N) and two aneuploid tumor clones (A1, B1). The central UMAP plots present DNA clustering results based on copy number in each bin. Adjacent heatmaps show the copy number variations for each single nucleus, arranged by cluster identity on the x-axis against genomic bins on the y-axis, with red indicating amplification and blue indicating deletion. **B**: Aggregated copy number profiles derived from summing across all single cells within the same cluster, comparing hybrid (green) and DNA-only (red) data. **C**: RNA expression analysis of merged hybrid data and RNA-only data. In the center, hybrid and RNA-only data are co-clustered into seven expression clusters: three tumor expression clusters (Ta1, Tb1, and Tc1) and four somatic cell types—epithelial cells (EP), endothelial cells (EC), fibroblasts, and monocytes (Mono). The merged clusters are then split into nuclei from the hybrid protocol (left) and the RNA-only protocol (right). **D**: RNA-layer expression data from each platform showed similar clustering results. The nuclei from both platforms are either clustered independently (hybrid data on the far left and RNA-only data on the far right) or clustered together as shown in the center plots, taken from Panel C. The heatmaps quantify the agreement between the co-clustering results with the respective hybrid and RNA-only clustering results.

### Figure S5

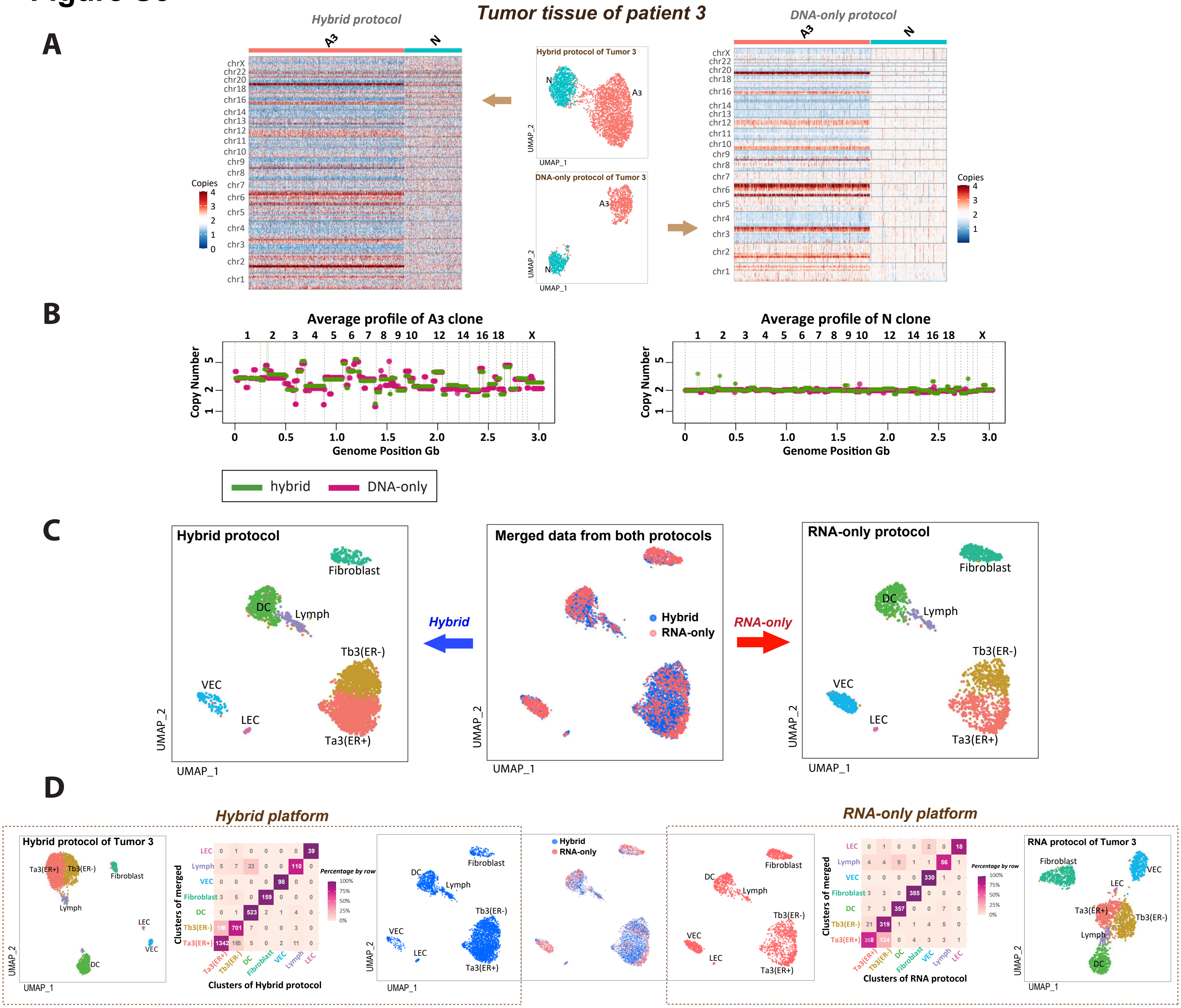


**Figure S5. Comparison of DNA and RNA clustering results between hybrid protocol and DNA-only/RNA-only protocols for Patient 3.** Following the structure of Figure 2, this figure provides a comparison of the hybrid protocol data to the DNA-only and RNA-only protocol datasets for the tumor sample of patient 3. **A**: Single-cell DNA copy number analysis of the tumor sample comparing the hybrid protocol data (left) with the DNA-only protocol data (right). The tumor sample has two distinct copy number profiles: normal diploid cells (N) and one aneuploid tumor profile (A3). The central UMAP plots present DNA clustering results based on copy number in each bin. Adjacent heatmaps show the copy number variations for each single nucleus, arranged by cluster identity on the x-axis against genomic bins on the y-axis, with red indicating amplification and blue indicating deletion. **B**: Aggregated copy number profiles derived from summing across all single cells within the same cluster, comparing hybrid (green) and DNA-only (red) data.
**C**: RNA expression analysis of merged hybrid data and RNA-only data. IIn the center, hybrid and RNA-only data are co-clustered into seven expression clusters: two tumor expression clusters, Ta3 and Tb3, that are ER+ and ER- respectively, and five somatic cell types—vascular endothelial cells (VEC), lymphatic endothelial cells (LEC), Fibroblast, and lymphocytes (Lymph) dendritic cells (DC). The merged clusters are then split into nuclei from the hybrid protocol (left) and the RNA-only protocol (right). **D**: RNA-layer expression data from each platform showed similar clustering results. The nuclei from both platforms are either clustered independently (hybrid data on the far left and RNA-only data on the far right) or clustered together as shown in the center plots, taken from Panel C. The heatmaps quantify the agreement between the co-clustering results with the respective hybrid and RNA-only clustering results.

### Figure S6

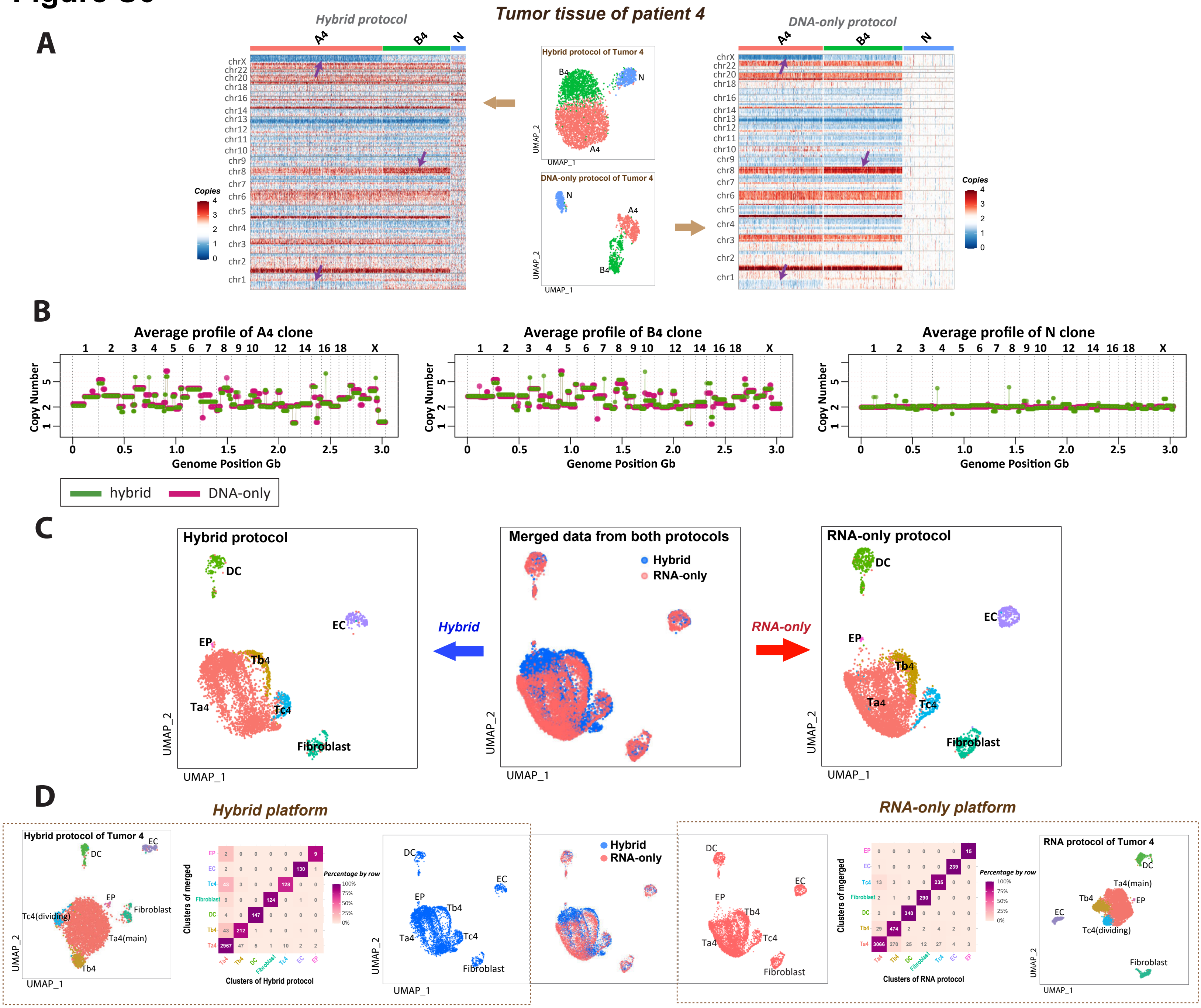


**Figure S6. Comparison of DNA and RNA clustering results between hybrid protocol and DNA-only/RNA-only protocols for Patient 4.** Following the structure of Figure 2, this figure provides a comparison of the hybrid protocol data to the DNA-only and RNA-only protocol datasets for the tumor sample of patient 4. **A**: Single-cell DNA copy number analysis of the tumor sample comparing the hybrid protocol data (left) with the DNA-only protocol data (right). The tumor sample has three distinct copy number profiles: normal diploid cells (N) and two aneuploid tumor clones (A4, B4). The central UMAP plots present DNA clustering results based on copy number in each bin. Adjacent heatmaps show the copy number variations for each single nucleus, arranged by cluster identity on the x-axis against genomic bins on the y-axis, with red indicating amplification and blue indicating deletion. **B**: Aggregated copy number profiles derived from summing across all single cells within the same cluster, comparing hybrid (green) and DNA-only (red) data. **C**: RNA expression analysis of merged hybrid data and RNA-only data. In the center, hybrid and RNA-only data are co-clustered into seven expression clusters: three tumor expression clusters (Ta4, Tb4, and Tc4) and four somatic cell types—epithelial cells (EP), endothelial cells (EC), dendritic cells (DC), and Fibroblast. The merged clusters are then split into nuclei from the hybrid protocol (left) and the RNA-only protocol (right). **D**: RNA-layer expression data from each platform showed similar clustering results. The nuclei from both platforms are either clustered independently (hybrid data on the far left and RNA-only data on the far right) or clustered together as shown in the center plots, taken from Panel C. The heatmaps quantify the agreement between the co-clustering results with the respective hybrid and RNA-only clustering results.

### Figure S7

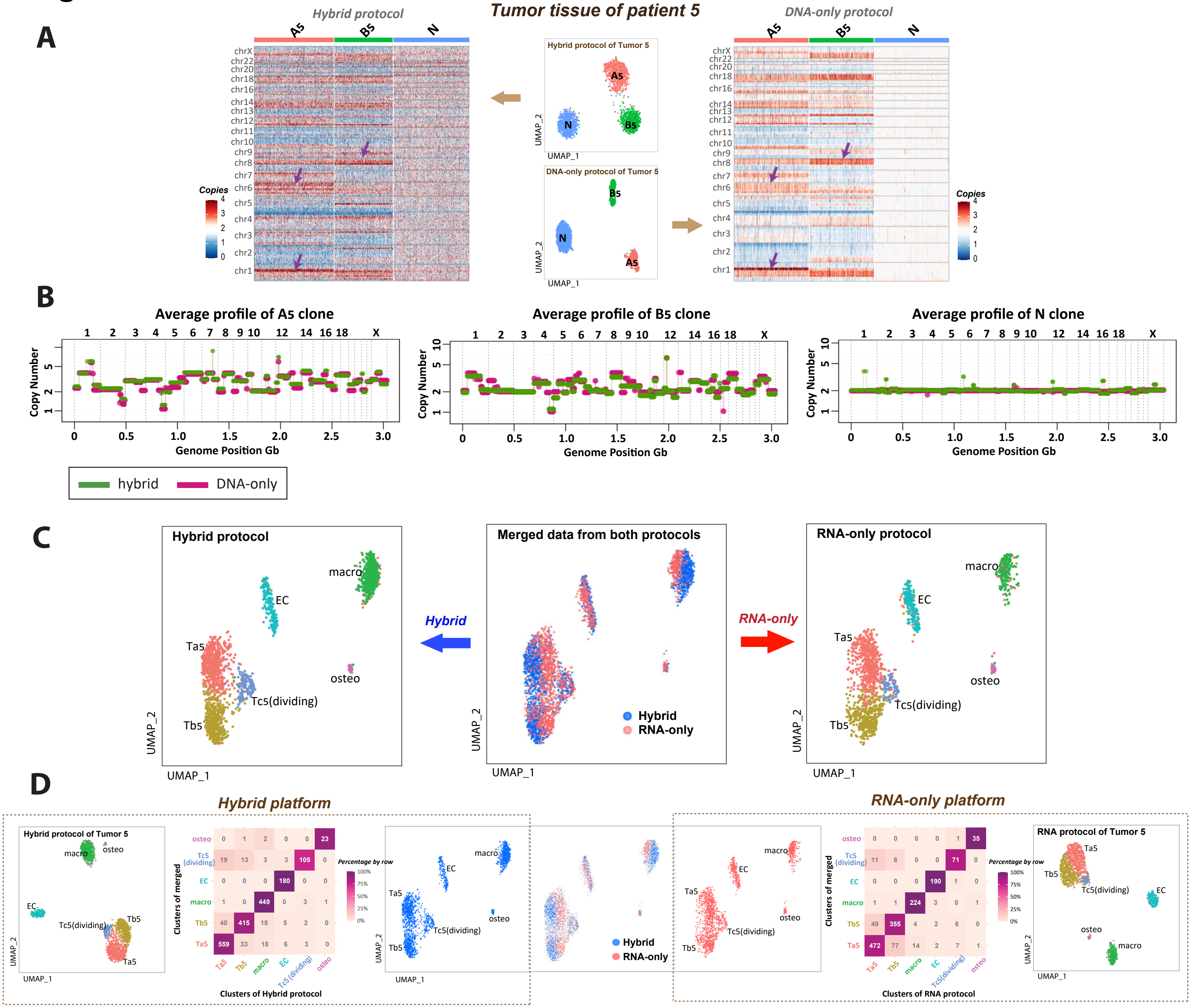


Hybrid platform heatmap (rows: clusters of merged; columns: clusters of Hybrid protocol):

| | Ta5 | Tb5 | macro | EC | Tc5 (dividing) | osteo |
|---|---|---|---|---|---|---|
| osteo | 0 | 1 | 2 | 0 | 0 | 23 |
| Tc5 (dividing) | 19 | 13 | 3 | 3 | 105 | 0 |
| EC | 0 | 0 | 0 | 180 | 0 | 0 |
| macro | 1 | 0 | 449 | 0 | 3 | 1 |
| Tb5 | 40 | 415 | 18 | 5 | 2 | 0 |
| Ta5 | 559 | 33 | 18 | 6 | 3 | 0 |

RNA-only platform heatmap (rows: clusters of merged; columns: clusters of RNA protocol):

| | Ta5 | Tb5 | macro | EC | Tc5 (dividing) | osteo |
|---|---|---|---|---|---|---|
| osteo | 0 | 0 | 0 | 0 | 1 | 35 |
| Tc5 (dividing) | 11 | 6 | 0 | 0 | 71 | 0 |
| EC | 0 | 0 | 0 | 190 | 1 | 0 |
| macro | 1 | 1 | 224 | 3 | 0 | 1 |
| Tb5 | 49 | 355 | 4 | 2 | 6 | 0 |
| Ta5 | 472 | 77 | 14 | 2 | 7 | 1 |

**Figure S7. Comparison of DNA and RNA clustering results between hybrid protocol and DNA-only/RNA-only protocols for Patient 5.** Following the structure of Figure 2, this figure provides a comparison of the hybrid protocol data to the DNA-only and RNA-only protocol datasets for the tumor sample of patient 5. **A**: Single-cell DNA copy number analysis of the tumor sample comparing the hybrid protocol data (left) with the DNA-only protocol data (right). The tumor sample has three distinct copy number profiles: normal diploid cells (N) and two aneuploid tumor clones (A5, B5). The central UMAP plots present DNA clustering results based on copy number in each bin. Adjacent heatmaps show the copy number variations for each single nucleus, arranged by cluster identity on the x-axis against genomic bins on the y-axis, with red indicating amplification and blue indicating deletion. **B**: Aggregated copy number profiles derived from summing across all single cells within the same cluster, comparing hybrid (green) and DNA-only (red) data. **C**: RNA expression analysis of merged hybrid data and RNA-only data. In the center, hybrid and RNA-only data are co-clustered into six expression clusters: three tumor expression clusters (Ta5, Tb5, and Tc5) and three somatic cell types—endothelial cells (EC), osteoclasts (osteo), and macrophage (macro). The merged clusters are then split into nuclei from the hybrid protocol (left) and the RNA-only protocol (right). **D**: RNA-layer expression data from each platform showed similar clustering results. The nuclei from both platforms are either clustered independently (hybrid data on the far left and RNA-only data on the far right) or clustered together as shown in the center plots, taken from Panel C. The heatmaps quantify the agreement between the co-clustering results with the respective hybrid and RNA-only clustering results.

### Figure S8

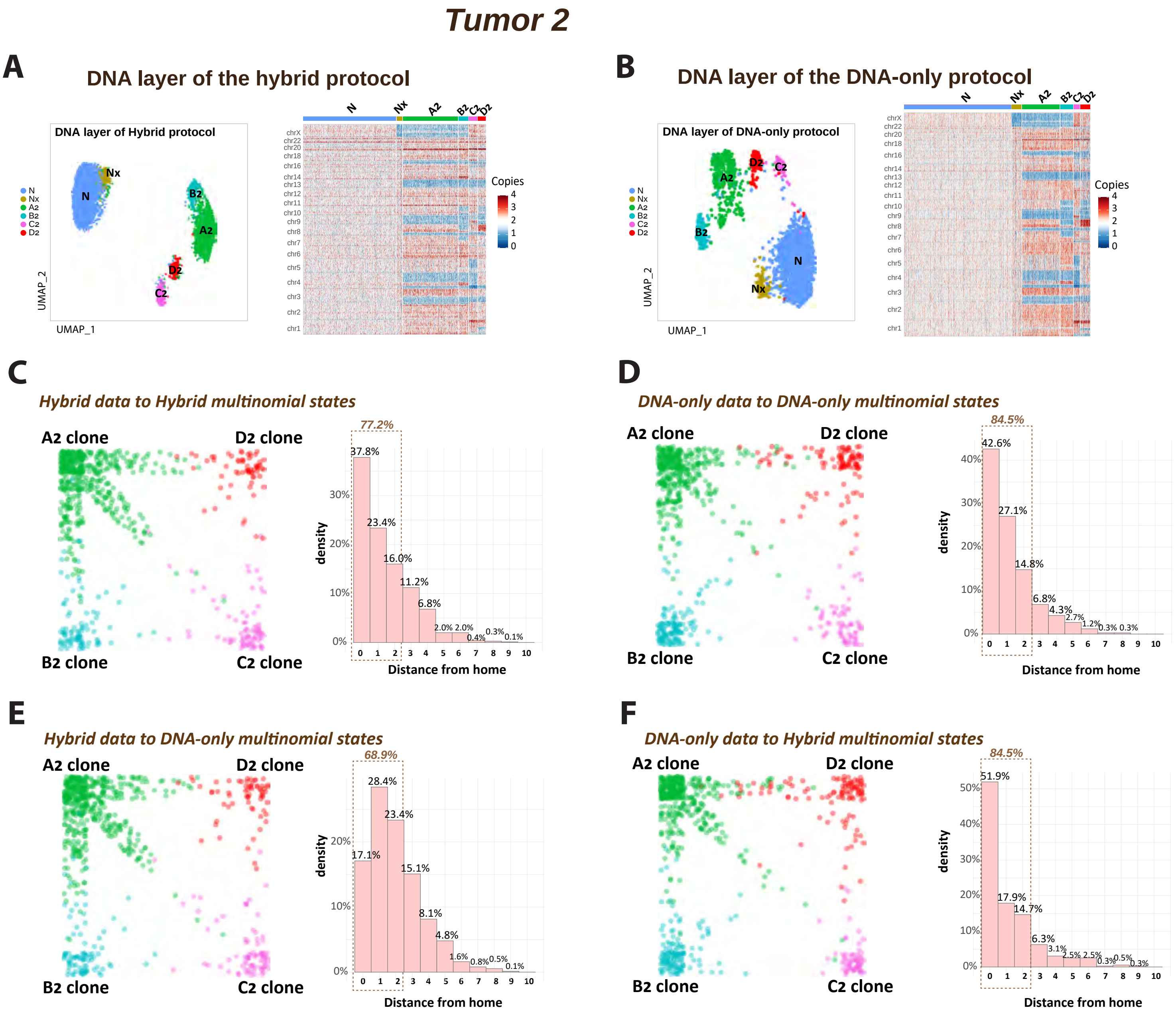


**Figure S8. Multinomial analysis comparing the DNA layer from the hybrid protocol with the DNA-only protocol for similar clustering efficacy**.

**A**: UMAP (left) and copy-number heatmap (right) for genomic copy-number clustering of the DNA layer from the hybrid protocol. **B**: UMAP (left) and copy-number heatmap (right) for genomic copy-number clustering of the DNA layer from the DNA-only protocol. **C**: Multinomial wheel (left) showing each nucleus from the hybrid protocol in its most likely position as determined by the Seurat-cluster centroids from the same hybrid protocol. A corresponding histogram (right) illustrates the distribution of the distance from each nucleus to its corresponding cluster centroid. **D**: Multinomial wheel (left) showing each nucleus from the DNA-only protocol in its most likely position as determined by the Seurat-cluster centroids from the same DNA-only protocol. A corresponding histogram (right) illustrates the distribution of the distance from each nucleus to its corresponding cluster centroid. **E**: Multinomial wheel (left) showing each nucleus from the hybrid protocol in its most likely position as determined by the Seurat-cluster centroids from the DNA-only protocol. A corresponding histogram (right) illustrates the distribution of the distance from each nucleus to its corresponding cluster centroid. **F**: Multinomial wheel (left) showing each nucleus from the DNA-only protocol in its most likely position as determined by the Seurat-cluster centroids from the hybrid protocol. A corresponding histogram (right) illustrates the distribution of the distance from each nucleus to its corresponding cluster centroid.

#### Figure S9

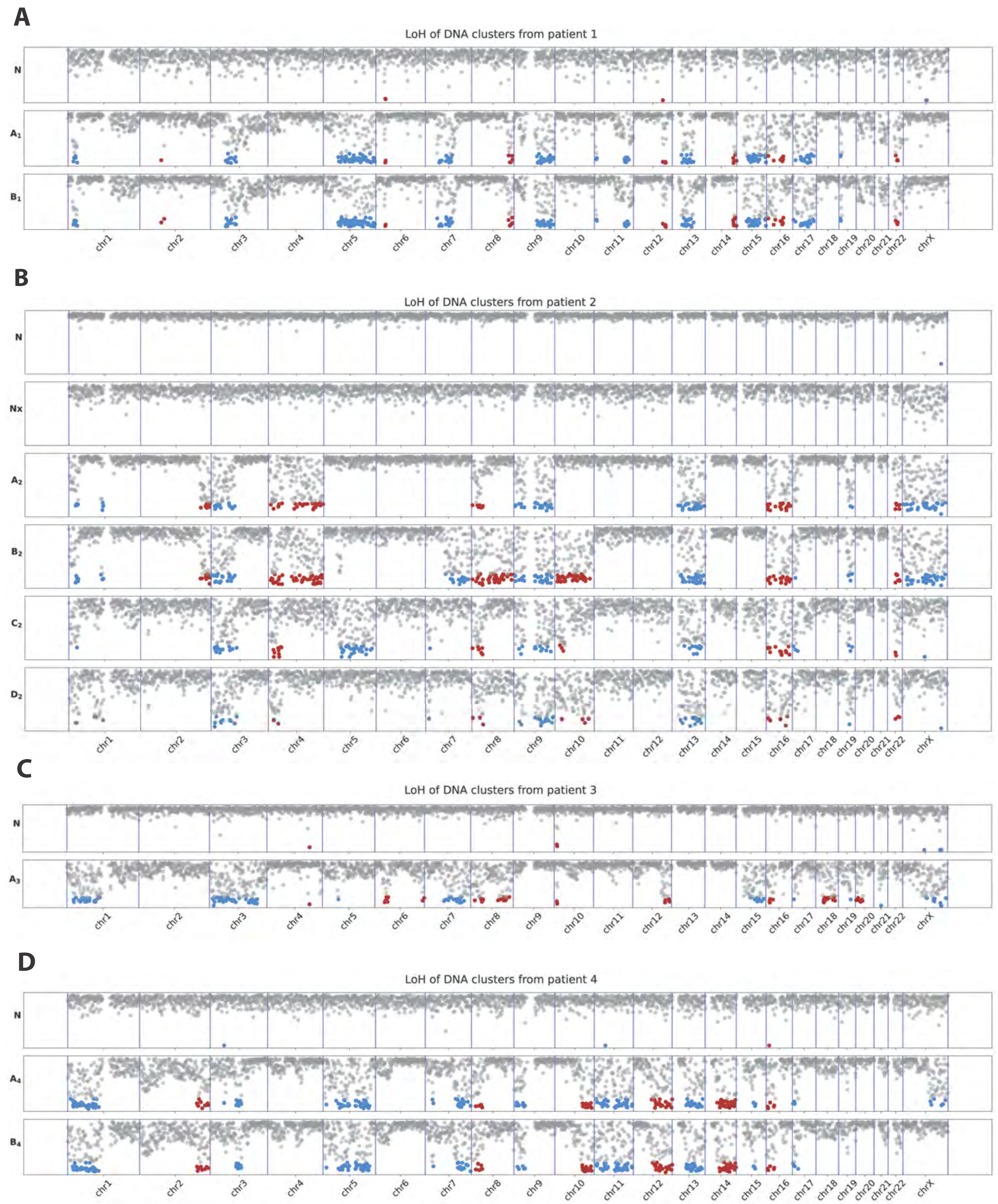
A
LoH of DNA clusters from patient 1
N
A₁
B₁
chr1 chr2 chr3 chr4 chr5 chr6 chr7 chr8 chr9 chr10 chr11 chr12 chr13 chr14 chr15 chr16 chr17 chr18 chr19 chr20 chr21 chr22 chrX
B
LoH of DNA clusters from patient 2
N
Nx
A₂
B₂
C₂
D₂
chr1 chr2 chr3 chr4 chr5 chr6 chr7 chr8 chr9 chr10 chr11 chr12 chr13 chr14 chr15 chr16 chr17 chr18 chr19 chr20 chr21 chr22 chrX
C
LoH of DNA clusters from patient 3
N
A₃
chr1 chr2 chr3 chr4 chr5 chr6 chr7 chr8 chr9 chr10 chr11 chr12 chr13 chr14 chr15 chr16 chr17 chr18 chr19 chr20 chr21 chr22 chrX
D
LoH of DNA clusters from patient 4
N
A₄
B₄
chr1 chr2 chr3 chr4 chr5 chr6 chr7 chr8 chr9 chr10 chr11 chr12 chr13 chr14 chr15 chr16 chr17 chr18 chr19 chr20 chr21 chr22 chrX

E

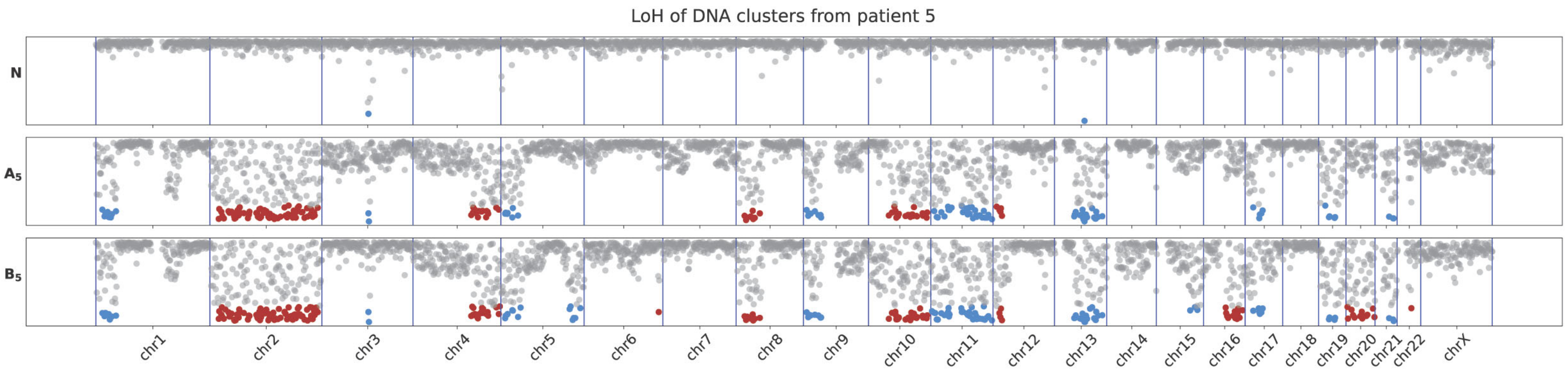


**Figure S9: Allele imbalance analysis in DNA clones from Tumors 1-5**.

Allele imbalance plots for each DNA cluster, as determined from aggregated nuclei from the DNA-only data of tumors 1-5. The plots demonstrate agreement in loss of heterozygosity (LoH) patterns with the copy number variation illustrated in Figures 2, 3 and Figures S4-S7. **A-E** apply to the tumor samples from patients 1-5 respectively. Each panel contains a plot for each DNA cluster, with the genome broken into bins of imputed phasing. For each bin, we plot the minor frequency such that y-axis varies from 0 to 0.5, where a value of 0.5 indicates perfectly balanced alleles and values below 0.05 are highlighted red or blue (colors alternate by chromosome to make chromosome boundaries distinct) to mark regions of LoH.

### Figure S10

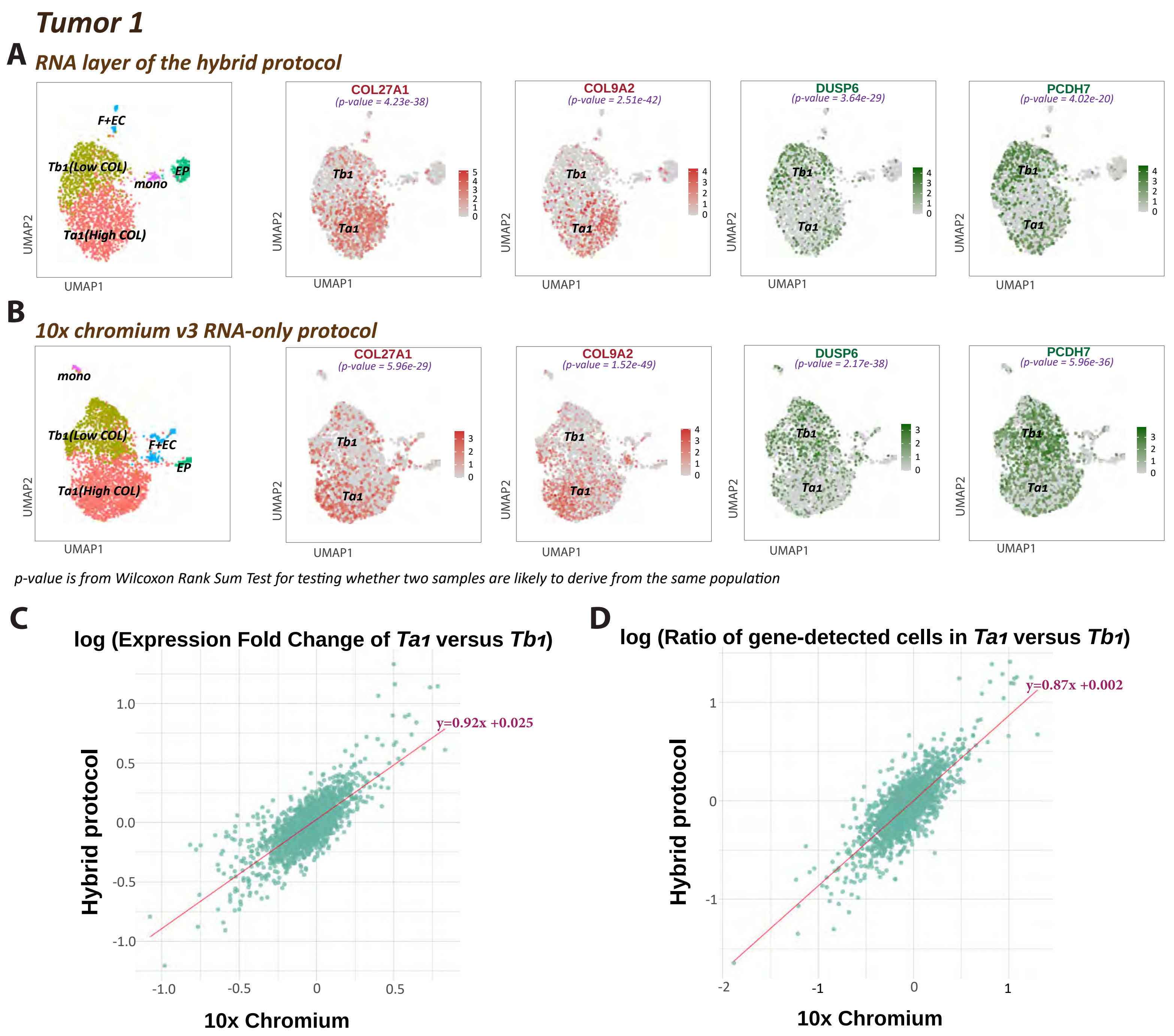


**Figure S10. Comparing RNA clustering between the hybrid protocol and 10x chromium v3 RNA-only protocol.**
**A and B** display UMAP clusterings of nuclei from Tumor 1, identifying two distinct tumor RNA clusters ($Ta_1$ and $Tb_1$) primarily based on collagen expression levels, which were comparable between the hybrid protocol (A) and the 10x Chromium V3 protocol (B). Feature plots for four marker genes highlight similar expression differences between the two tumor RNA clusters in both protocols.
**C** presents a scatter plot illustrating the log fold change of gene expression between clusters $Ta_1$ and $Tb_1$ for all genes detected in at least 10% of nuclei in either cluster, across both protocols. Each point represents a gene. Linear regression analysis shows a slope of 0.92 and a y-intercept of 0.025, indicating a strong correlation (Pearson's r, p-value < 2.2e-16).
**D** shows a scatter plot comparing the log ratio of the number of cells in which a gene is detected in Ta1 versus Tb1 clusters using both protocols. Each point represents a gene. Linear regression reveals a slope of 0.87 and a y-intercept of 0.002, with a strong correlation demonstrated by the correlation test (p-value < 2.2e-16).

#### Figure S11

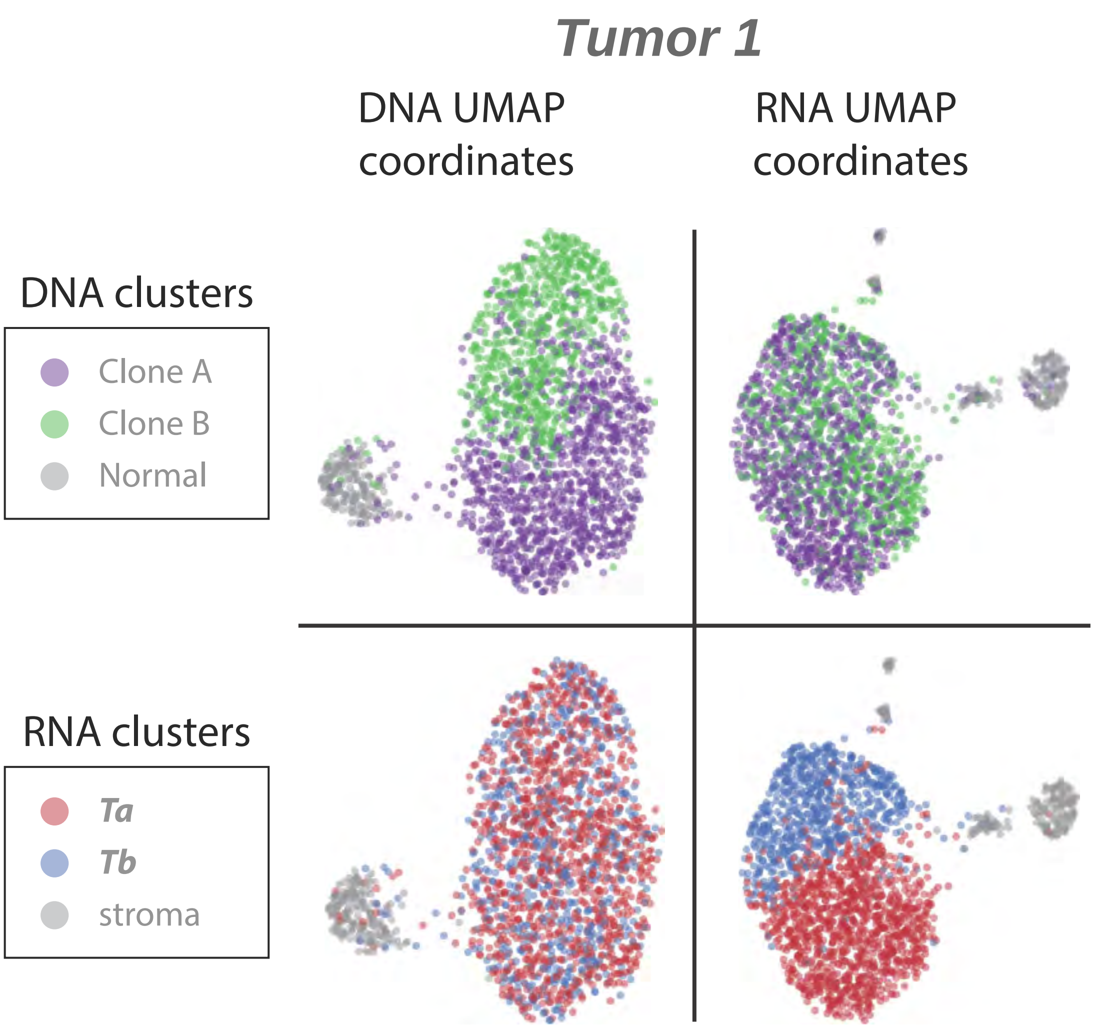


**Figure S11: Projection of RNA/DNA-cluster identities onto DNA/RNA UMAP plots showing independent DNA and RNA clustering for Tumor 1.**

A 2x2 UMAP plot matrix visually represents the clustering of the DNA-layer and RNA-layer, using hybrid data from Tumor 1.

Top Left: UMAP plot of DNA clustering, colored by DNA clonal information.

Bottom Right: UMAP plot of RNA clustering, colored by RNA clustering information.

Bottom Left: RNA clustering identities projected onto DNA UMAP coordinates.

Top Right: DNA clonal identities projected onto RNA UMAP coordinates

### Figure S12

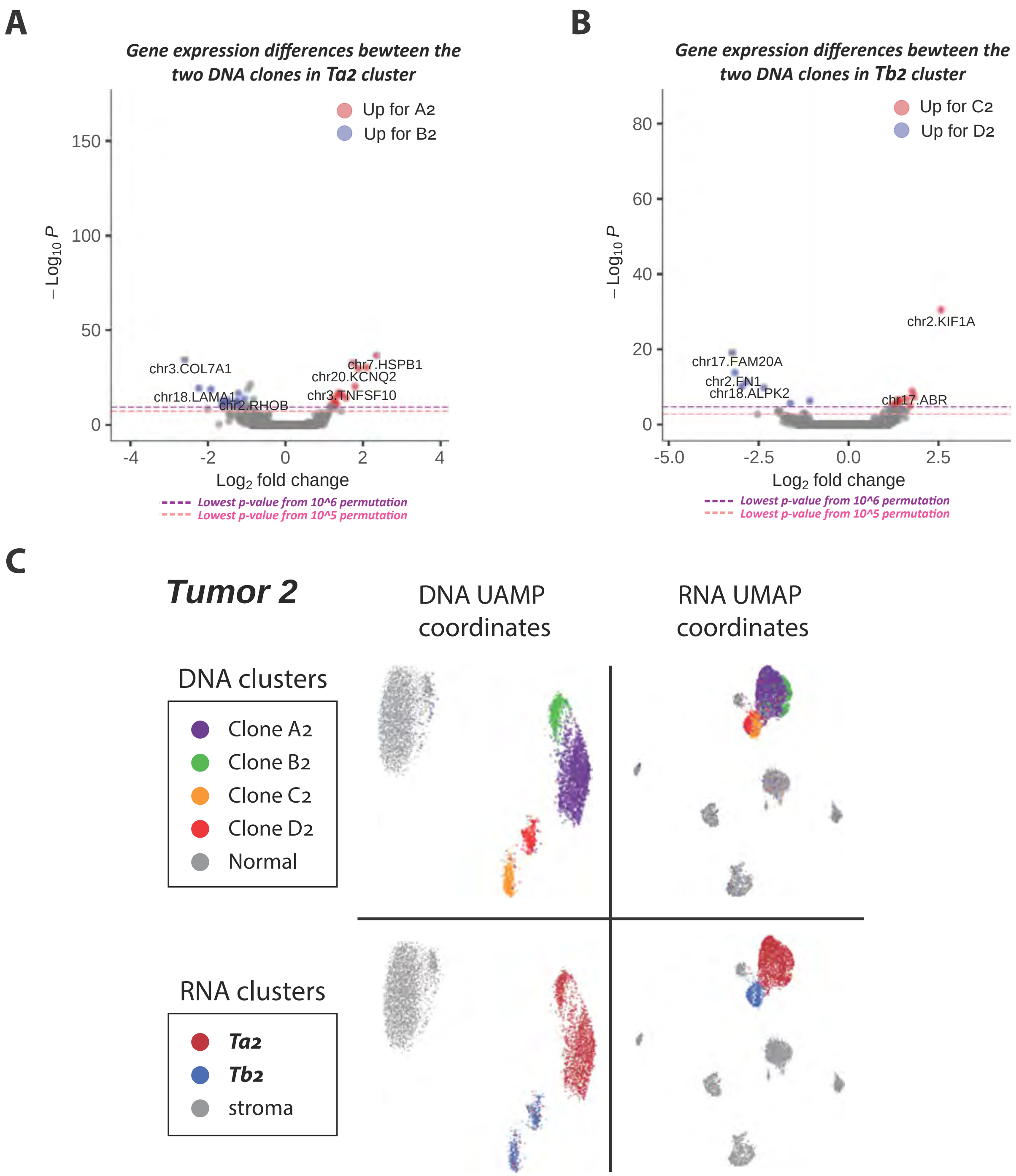


**Figure S12. Splitting RNA clusters by DNA clones of Tumor 2.**
**A:** A volcano plot highlights the genes with statistically significant expression differences between nuclei with $Ta_2$ expression from clone $A_2$ ($Ta_2$-$A_2$) and nuclei with $Ta_2$ expression from clone $B_2$ ($Ta_2$-$B_2$). The log2 fold-change is plotted along the x-axis with the statistical significance of the observed counts on the y-axis.
**B:** A volcano plot highlights the genes with statistically significant expression differences between nuclei with $Tb_2$ expression from clone $C_2$ ($Tb_2$-$C_2$) and nuclei with $Tb_2$ expression from clone $D_2$ ($Tb_2$-$D_2$). The log2 fold-change is plotted along the x-axis with the statistical significance of the observed counts on the y-axis.
**C:** UMAP visualizations of DNA and RNA data from Tumor 2. The top left plot shows DNA clusters projected onto DNA UMAP coordinates, while the top right plot shows DNA clusters projected onto RNA UMAP coordinates. The bottom left plot shows RNA clusters projected onto DNA UMAP coordinates, and the bottom right plot shows RNA clusters projected onto RNA UMAP coordinates. For the DNA layer, the "Normal" population (grey) combines N and Nx clusters. For the RNA layer, the "stroma" population (grey) combines all non-tumor types.

#### Figure S13

##### *Tumor 3*

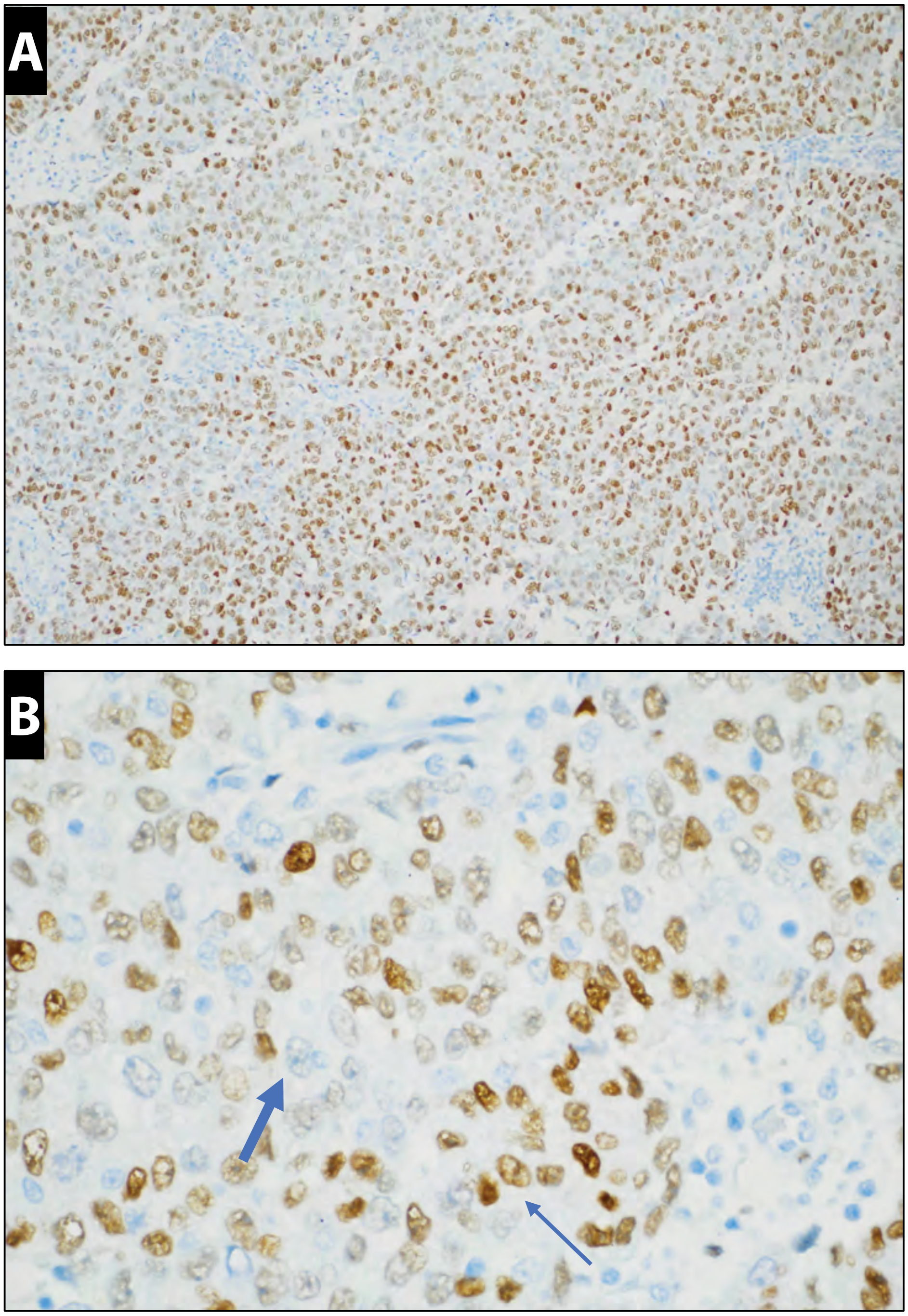


**Figure S13. ER protein expression in an adenocarcinoma tumor (Tumor 3) by IHC.**
The expression data from the tumor tissue of patient 3 showed two distinct expression clusters with significant differences in ER expression.
**A:** ER immunostaining at ×200, exhibits solid architecture with diffuse estrogen receptor (ER) positivity in the neoplastic cells.
**B** at increased magnification of ×400, shows that ER immunoreactivity and high grade cytologic atypia is demonstrated in ER positive (thin arrow) and ER negative (thick arrow) tumor cells. The nuclei of the tumor cells are moderately pleomorphic with irregular contours and some contain prominent nucleoli.

### Figure S14

#### Tumor 4

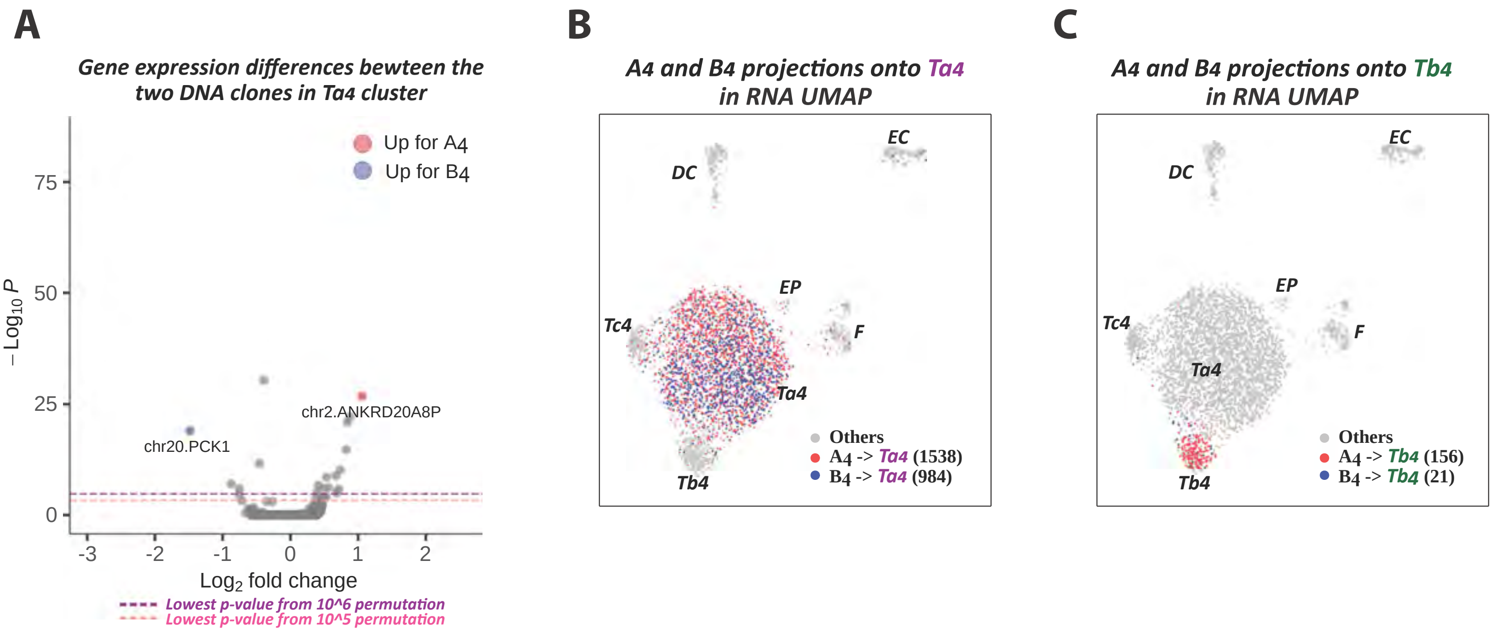


**Figure S14. Differential gene analysis and tumor DNA clone projections onto RNA clusters using Tumor 4.**
**A**: A volcano plot highlights the genes with statistically significant expression differences between nuclei with Ta4 expression from clone A4 (Ta4-A4) and nuclei with Ta4 expression from clone B4 (Ta4-B4). The log2 fold-change is plotted along the x-axis with the statistical significance of the observed counts on the y-axis.
**B**: Projection of DNA clonal identities A4 and B4 onto the RNA UMAP of the Ta4 cluster.
**C**: Projection of DNA clonal identities A4 and B4 onto the RNA UMAP of the Tb4 cluster.

### Figure S15

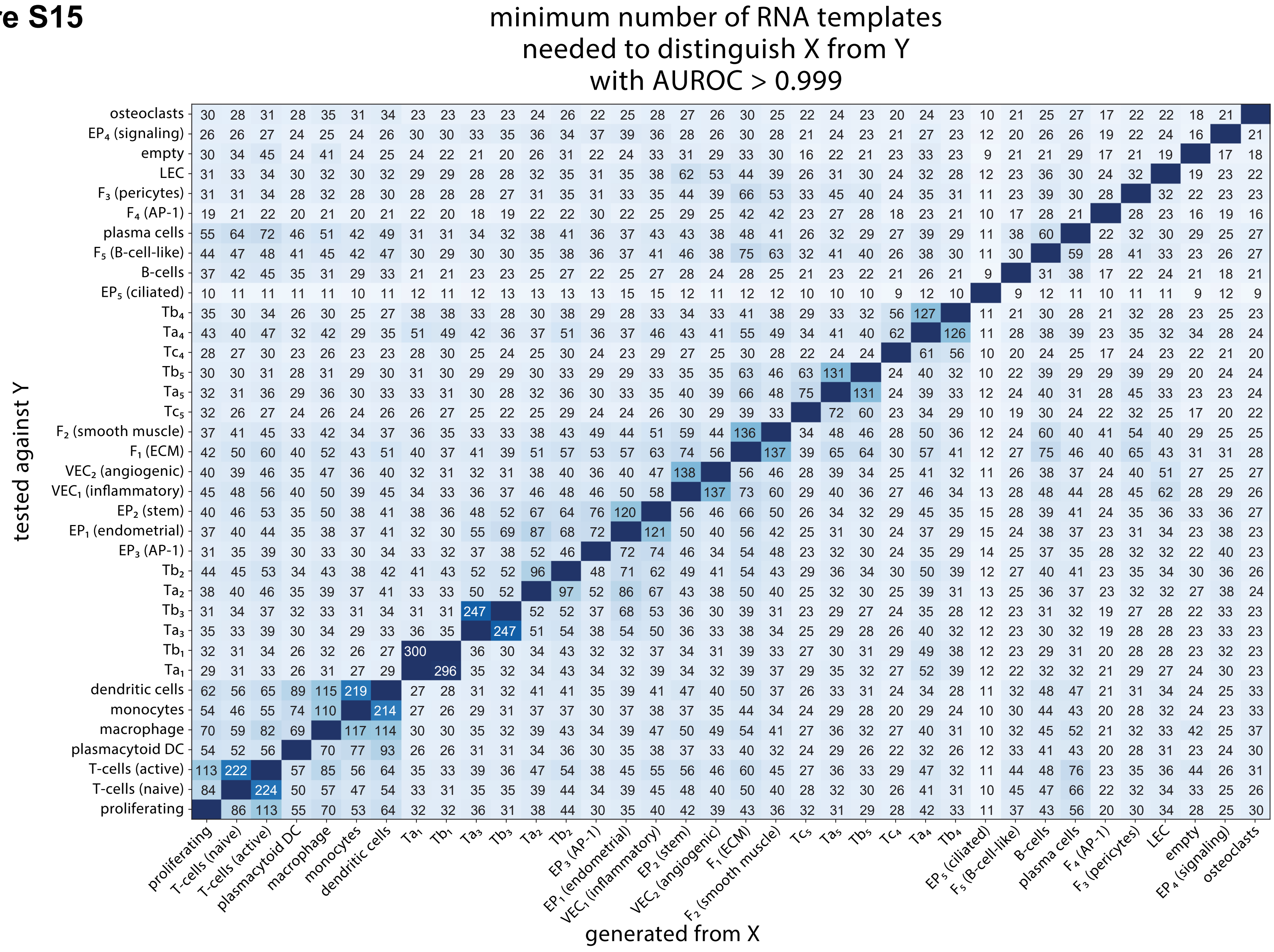


**Figure S15. Quantifying cluster similarity using multinomial distributions.** This heatmap visualized the minimum number of RNA templates, M[i, j], required to confidently differentiate a cell from cluster i on X-axis from belonging to cluster j on the Y-axis, with an AUROC greater than 0.999. Smaller values indicate a high distinctiveness between cluster pairs, allow for differentiation of nuclei with fewer observations. Conversely, larger values suggest greater similarity, requiring more reads to reliably distinguish one from the other. The diagonal, representing comparisons of each cluster to itself, is intentionally blank since a cluster is not differentiable from itself for any number of reads.

### Figure S16

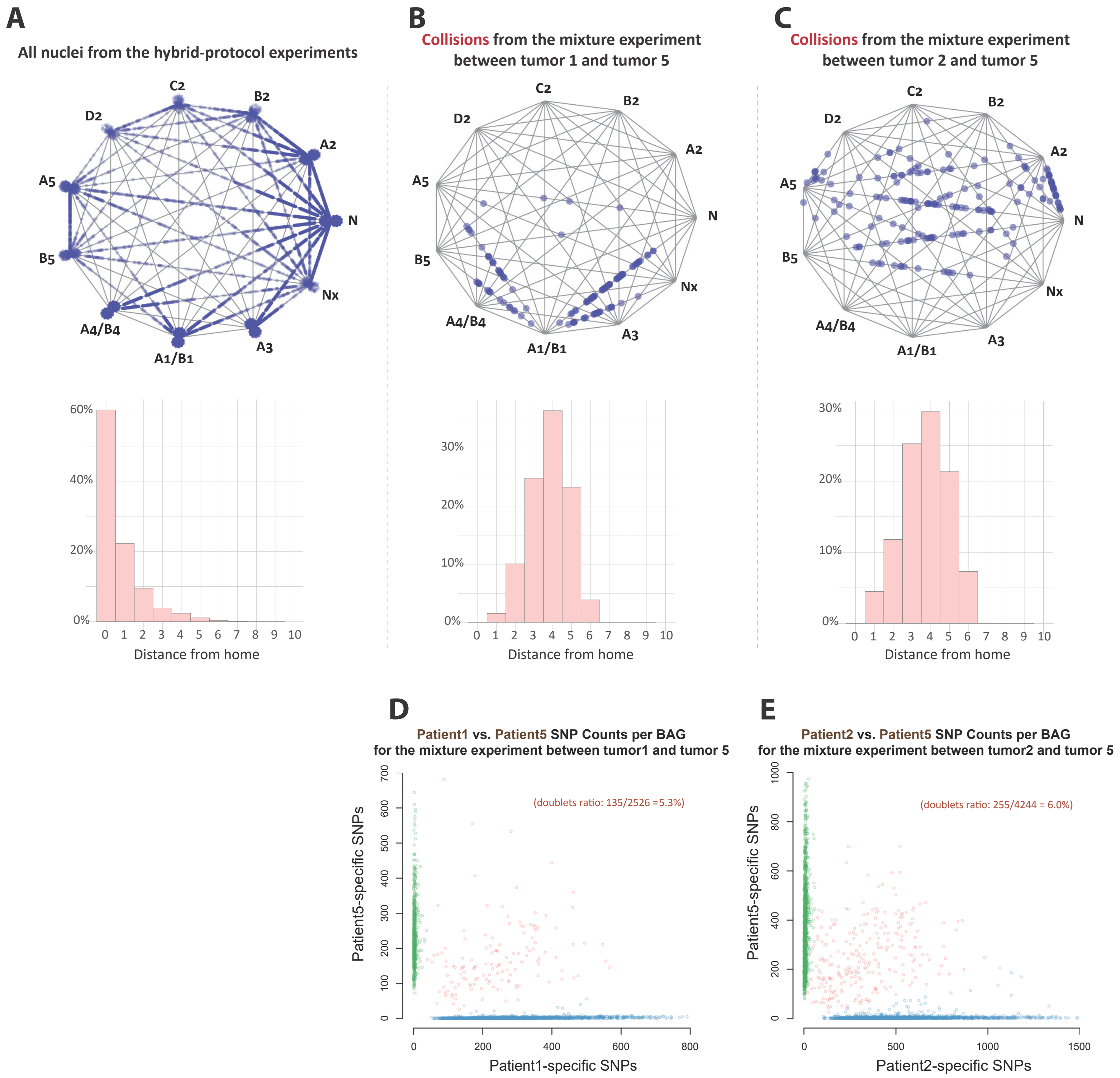
A
All nuclei from the hybrid-protocol experiments
C2
B2
D2
A2
A5
N
B5
Nx
A4/B4
A3
A1/B1
60%
40%
20%
0%
0 1 2 3 4 5 6 7 8 9 10
Distance from home
B
Collisions from the mixture experiment
between tumor 1 and tumor 5
30%
20%
10%
0%
0 1 2 3 4 5 6 7 8 9 10
Distance from home
C
Collisions from the mixture experiment
between tumor 2 and tumor 5
30%
20%
10%
0%
0 1 2 3 4 5 6 7 8 9 10
Distance from home
D
Patient1 vs. Patient5 SNP Counts per BAG
for the mixture experiment between tumor1 and tumor 5
(doublets ratio: 135/2526 =5.3%)
Patient5-specific SNPs
0 100 200 300 400 500 600 700
0 200 400 600 800
Patient1-specific SNPs
E
Patient2 vs. Patient5 SNP Counts per BAG
for the mixture experiment between tumor2 and tumor 5
(doublets ratio: 255/4244 = 6.0%)
Patient5-specific SNPs
0 200 400 600 800 1000
0 500 1000 1500
Patient2-specific SNPs

**Figure S16: Multinomial wheel to identify DNA crossovers.**

**A:** The DNA bin counts of hybrid nuclei are modeled using a multinomial distribution based on eleven distinct copy number variation (CNV) profiles. To identify nuclei that poorly fit their most probably cluster or "home" (**H**), we expand the multinomial space to include pairwise linear combinations of vector **H** with another cluster (**Z**), of the form $\boldsymbol{\alpha}\mathbf{H}$ + (1 - $\boldsymbol{\alpha}$)**Z** where $\boldsymbol{\alpha}$ ranges from 0 to 1, in 0.1 increments. This extension results in 101 possible multinomial distributions per home. For each nucleus (blue dots), we assign the wheel vertex that maximizes the likelihood of its bin counts. Nuclei perfectly matching their home cluster are positioned just outside the wheel perimeter. The histogram below represents the "distance from home" for all nuclei, calculated as 10$\boldsymbol{\alpha}$. The data show that over 90% of nuclei are situated within a distance 2 from their home cluster.

**B and C** echo the methodology of panel A, focusing exclusively on nuclei identified as collisions based on single nucleotide variant (SNV) data from two mixture experiments: one combining tumor tissue materiel from patient 1 with that from patient 5 (**B**); and another combining tumor tissue materiel from patient 2 with that from patient 5 (**C**). Unlike Panel A, these panels visualize only the collision instances as detailed in the subsequent Panels D and E. The histograms illustrate that 10-15% of collisions are within 2 units of home.

**D** presents a per-nucleus scatter plot showing the number of germline SNVs unique to patient 1 (x-axis) or patient 5 (y-axis). Nuclei that are confidently from patient 1 are colored in blue, those that are confidently from patient 5 are colored in green, and suspected collisions are marked in red. In total, we observe 135 collisions from a total of 2526 nuclei for a rate of 5.3%.

**E** presents a per-nucleus scatter plot showing the number of germline SNVs unique to patient 2 (x-axis) or patient 5 (y-axis). Nuclei that are confidently from patient 2 are colored in blue, those that are confidently from patient 5 are colored in green, and suspected collisions are marked in red. In total, we observe 255 collisions from 4244 total nuclei for a rate of 6.0%.
