## Supplementary Protocol for "DNA and RNA from the same single nucleus reveals interactions between genomic and transcriptomic landscapes in human tumor samples"

### Single-nuclei 3ps Hybrid BAG-seq Laboratory Protocol

#### Equipment

| Name | Company | Catalog Number |
|---|---|---|
| Inverted Microscope | Motic | AE31 |
| Legato 100 Syringe Pumps (3) | KD Scientific | 788100 |
| Tube Revolver | Thermo Fisher Scientific | 88881001 |
| Isotemp 60L incubator | Thermo Fisher Scientific | 151030513 |
| 3 mL Syringes | Becton Dickinson | 309657 |
| 10 mL Syringes | Becton Dickinson | 302995 |
| PE-2 Tubing | Scientific Commodities | BB31695-PE/2 |
| 26 Gauge Needles | Becton Dickinson | 305111 |
| 22 Gauge Needles | Becton Dickinson | 305155 |
| 16 Gauge Needles | Becton Dickinson | 305197 |
| PDMS co-flow microfluidic droplet generation device for "Drop-Seq" | Nanoshift, LLC | |
| 150 µm PluriStrainer | PluriSelect | 43-50150-03 |
| 40 µm Cell Strainer | Corning | 431750 |
| Microseal 'B' plate seals | Bio-Rad | MSB1001 |
| SimpliAmp Thermal Cycler | Thermo Fisher Scientific | A24812 |

#### Primers:

| Name | Sequence |
|---|---|
| TG-vt-K | /5ACryd//iSp18/TGTGTTGGGTGTGTTTGG NBBBB KKKKKKKGKKKKKKKKNN |
| TG-vt-polyT | /5ACryd//iSp18/TGTGTTGGGTGTGTTTGG DDDDDNNN T(30) |
| Template-switch-oligo (TSO) | AAGGCTCTTGACATGAAGGATAGrGrGrG |
| Adapter1-BC1-vt1-X (X = 1,2,…,96) | ACGCAGAGTCCGCTC <BAG-barcode1> NNNN CA+T+G (detailed sequences at bottom) |
| Adapter2-BC2-vt2-Adapter1-Y (Y = 1,2,…,96) | GACCGACTCGCATTACCCTAT <BAG-barcode2> NNNN ACGCAGAGTCCGCTC/3InvdT/ (detailed sequences at bottom) |
| Adapter3-BC3-Adapter2-Z (Z = 1,2,…,96) | AAGGCTCTTGACACGAAGGATAG <BAG-barcode3> GACCGACTCGCATTACCCT*A*T (detailed sequences at bottom) |
| P5-Adapter3 | AATGATACGGCGACCACCGAGATCTACAC CGCTGTCC AAGGCTCTTGACACGAAGGAT*A*G |
| TG primer | TGTGTTGGGTGTGTTT*G*G |
| Read1-Adapter3 | CGCTGTCC AAGGCTCTTGACACGAAGGATAG |
| Nextera_N70x (x = 1,2,…,12) | CAAGCAGAAGACGGCATACGAGAT <index_x> GTCTCGTGGGCTCGG (x = 1,2,…,12) (detailed sequences at bottom) |

#### Chemical Reagents:

| Name | Company | Catalog Number |
|---|---|---|
| **Nuclei Preparation** | | |
| 10× PBS | Thermo Fisher Scientific | AM9624 |

| | | |
|---|---|---|
| UltraPure BSA | Thermo Fisher Scientific | AM2618 |
| DAPI | Thermo Fisher Scientific | 62248 |
| RNase Plus RNase Inhibitor | Promega Corporation | N2615 |
| Surfact-Amps NP-40 (10% solution) | Thermo Fisher Scientific | 28324 |
| **Aqueous Phase 2** | | |
| Acrylamide/bis-acrylamide, 40% solution | Sigma-Aldrich | A9926 |
| Acrylamide solution, 40% | Sigma-Aldrich | A4058 |
| 0.5 M EDTA | Thermo Fisher Scientific | AM9260G |
| 20% Sarkosyl | Sigma-Aldrich | L7414 |
| 0.1 M DTT | Thermo Fisher Scientific | 707265ML |
| Ammonium persulfate | Sigma-Aldrich | 09913-100G |
| 10% NP40 | | |
| **Oil Phase** | | |
| HFE-7500 | Oakwood Chemical | 051243 |
| FC-40 | Sigma-Aldrich | F9755 |
| 008-Fluorosurfactant | Ran Technologies | 008-FluoroSurfactant-5G |
| TEMED | Bio-Rad | 1610801 |
| **Post-Droplet Generation** | | |
| Mineral Oil | Sigma-Aldrich | 69794-500ML |
| 20× SSC | Thermo Fisher Scientific | 15557044 |
| Perfluorooctanol (PFO) | Sigma-Aldrich | 370533 |
| 10 mM dNTP | Sigma-Aldrich | 11814362001 |
| DNA Polymerase I, Large (Klenow) Fragment | New England BioLabs | M0210M |
| Recombinant RNase Inhibitor | Takara Bio USA, Inc. | 2313A |
| Exonuclease I | New England BioLabs | M0293 |
| NlaIII | New England BioLabs | R0125L |
| Klenow Fragment (3-5 exo minus) | New England BioLabs | M0212L |
| Quick Ligase | New England BioLabs | M2200 |
| KAPA HiFi HotStart ReadyMix | Roche | KK2602 |
| Agencourt AMPure XP beads | Beckman Coulter | A63880 |
| Nextera XT DNA Library Prep Kit | Illumina | FC-131-1024 |

#### Chemical Solutions to Prepare

NST buffer (Recipe refers to Navin, Nicholas, et al. "Tumour evolution inferred by single-cell sequencing." Nature 472.7341 (2011): 90-94.)

PBS-BSA-DAPI buffer (1× PBS, 0.05% BSA, 0.5 µg/mL DAPI,  0.8 u/µL RNasein Plus RNase Inhibitor)

40% Surfactant in HFE-7500

10% Ammonium persulfate (make fresh solution for every experiment)

6× SSC

TE-SDS buffer (10 mM Tris pH 8.0, 1 mM EDTA, 0.5% SDS)

STOP-25 buffer (10 mM Tris pH 8.0, 25 mM EDTA, 0.1% Tween-20, 0.1 M KCl)

STOP-10 buffer (10 mM Tris pH 8.0, 10 mM EDTA, 0.1% Tween-20, 0.1 M KCl)

STOP-1 buffer (10 mM Tris pH 8.0, 1 mM EDTA, 0.1% Tween-20, 0.1 M KCl)

HBW buffer (10 mM Tris pH 8.0, 1 mM EDTA, 0.1% Tween-20)

### Single-nuclei Hybrid BAG-seq Experiment:

#### Equipment preparation

1. Set a heat block to 50°C
2. Set a heat block to 85°C
3. Set Isotemp 60L incubator to 37°C
4. Prepare Drop-Seq device

#### Nuclei Preparation

1. Prepare an ice bucket. Perform the following nuclei preparation steps on ice:
    a. Take a tube containing frozen pulverized tissue out of -80°C freezer, and keep the tube on ice.
    b. Add 1 mL ice-cold NST buffer into the sample tube. Pipette up and down several times.
    c. Transfer nuclei solution to a 15 mL conical tube prefilled with 5 mL NST buffer on ice. Pipette to mix.
    d. Place the conical tube on ice for 15 minutes with frequent shaking by hand (every 3 minutes).
    e. Centrifuge nuclei at 300 rcf and 4°C for 5 minutes. Discard supernatant.
    f. Resuspend nuclei pellet using 2 mL ice-cold PBS-BSA-DAPI buffer.
    g. Pass nuclei solution through 40 µm cell strainer.
2. Perform single-nucleus sorting based on DAPI-H vs. DAPI-A to remove debris and clumps. Sort nuclei into a 1.5 mL Eppendorf tube prefilled with 200 µL PBS-BSA-DAPI buffer.

#### Oil Phase Preparation

Combine the following; vortex to mix:

300 µL 40% surfactant
9.6 µL TEMED
2.1 mL HFE7500

#### Aqueous Phase 2 Preparation

Combine the following; pipet to mix:

| Aqueous Phase 2 Solution | Volume (in µL) | Final Concentration |
|---|---|---|
| Acrylamide/bis-acrylamide, 40% | 180 | |
| Acrylamide solution, 40% | 129 | |
| $H_2O$ | 131 | |
| 1 M Tris pH 7.5 | 80 | 40 mM |
| 0.5 M EDTA pH 8.0 | 50 | 12.5 mM |
| 500 µM TG-vt-polyT primer | 40 | 10 µM |
| 500 µM TG-vt-K primer | 80 | 20 µM |
| 20 % Sarkosyl | 20 | 0.2% |
| RNasin Plus RNase Inhibitor | 100 | |
| 0.1 M DTT | 30 | 1.5 mM |
| 10% NP-40 | 100 | 0.5% |
| 10% APS (add last) | 60 | 0.3% |
| ***Total*** | ***1 mL*** | |

#### Microfluidics Setup & Oil Droplet Collection

1. Turn on microscope. Place Drop-Seq microfluidic device on the microscope stage.
2. Insert a piece of PE/2 tubing into the outlet and place the other end into a waste container (a 15 mL conical tube).
3. Use the following syringes to collect the three solutions.
    a. Oil Phase: Use a 10 mL syringe with a 16G ½ needle to aspirate oil phase into syringe.
    b. Cell Solution or Aqueous Phase 2: Use a 3 mL syringe and a 22G ½ needle to aspirate the solution into respective syringes.
4. For each syringe, change the needle to a 26G ½ needle and insert it into a piece of PE/2 tubing.
5. Place syringes in syringe pumps and insert tubing into the corresponding inlets of the microfluidics device
    a. From left to right: Oil inlet – Aqueous Phase 2 inlet – Cell Solution inlet – Outlet
6. Once all syringes and tubing are in place, set the following flow rates:
    a. Oil Phase: 3,000 µL/hr
    b. Cell Solution: 750 µL/hr
    c. Aqueous Phase 2: 750 µL/hr
7. Begin by first turning on the cell solution pump.
8. Once cell solution is flowing, turn on the Aqueous Phase 2 pump.
9. Once the interphase between cell solution and Aqueous Phase 2 has stabilized, turn on the oil phase pump and wait for the stable droplets to form.
10. Once the flow has stabilized, place the output tubing on a glass slide under the microscope to assess the size uniformity of oil droplets.
11. Once droplet size is uniform, collect droplets in 1.5 mL Eppendorf tubes prefilled with 300 µL mineral oil.

12. After collection, incubate BAGs at 37°C for 2.5 hours -> 85°C heat block for 5 minutes -> 50°C for 10 minutes -> Room temperature cool-down for 15 minutes.

13. Set Isotemp 60L incubator to 42°C.

#### BAG Collection

1. Remove the top-layer oil using pipette. Remove the bottom-layer oil using syringe and 22G ½ needle.
2. Add 600 µL 6× SSC buffer and 150 µL perfluorooctanol (PFO). Invert the tube several times by hand to separate the oil phase from BAGs.
3. Centrifuge at 400 rcf for 1.5 minutes.
4. Discard the top and bottom layers, only leaving the middle translucent BAG layer in the tube.
5. Add 800 µL 6× SSC buffer and shake the tube multiple times. Centrifuge at 400 rcf for 1.5 minutes. Discard the top layer and any leftover PFO from the bottom layer.

#### Reverse Transcription

1. Follow the following steps to start the reverse transcription:
   a. Transfer ~350 µL BAGs from the BAG layer to a new 1.5 mL Eppendorf tube and wash once using the Pre-wash solution (850 µl 5x RT Buffer, 25 µl 0.1 M DTT, and 25 µl Takara Recombinant RNase Inhibitor).
   b. Pipette up and down, centrifuge at 400 rcf for 1.5 minutes, and discard the supernatant.
   c. Prepare the following solution in 1.5 mL Eppendorf tube on ice

| **BAG Solution** | **Volume (in µL)** |
|---|---|
| 5× RT Buffer | 100 |
| 10 mM dNTP | 100 |
| Takara Recombinant RNase Inhibitor | 25 |
| 0.1 M DTT | 20 |
| $H_2O$ | 570 |
| Maxima H- Reverse Transcriptase | 50 (Add last, on ice) |
| ***Total*** | ***865 µL*** |

   d. Immediately add the above mix into BAGs. Mix by pipetting up and down 10 times.
2. Incubate with rotation:
   a. Incubate at room temperature with rotation for 15 minutes.
   b. Add 25 µL 100 µM Template-switch-oligo (TSO).
   c. Incubate at room temperature with rotation for another 15 minutes.
   d. Incubate at 42°C with rotation for 1 hour.
   e. Add 10 µL DNA Polymerase I, Large (Klenow) Fragment (M0210M).
   f. Incubate at 42°C with rotation for another 30 minutes.

3. Stop the reaction
    a. Prefill a 15 mL conical tube with 5 mL STOP-25 buffer.
    b. Transfer all the BAG solution from the Eppendorf reaction tube to the 15 mL conical tube. Mix by pipetting 10 times.
    c. Incubate at room temperature for 5 minutes.
    d. Spin down at 300 rcf for 5 minutes. Remove supernatant.
    e. Wash BAGs twice using 1 mL STOP-10 buffer.

--------------------Safe Stopping Point: Store BAGs in STOP-10 buffer at 4°C overnight------------------

#### Exonuclease and RNase H Treatment

1. Set Isotemp 60L Incubator to 37°C.
2. If BAGs are stored in STOP-10, discard the supernatant.
3. Wash once with 1 mL STOP-1 buffer.
4. Wash once with 1× Exo I buffer (900 µL $H_2O$ + 100 µL 10× Exo I buffer).
5. Prepare the following exonuclease treatment solution and vortex.

| **Exonuclease Treatment Solution** | **Volume (µL)** |
|---|---|
| $H_2O$ | 835 |
| 10× Exo I buffer | 100 |
| Exo I enzyme | 50 |
| RNase H | 15 |
| ***Total*** | ***1 mL*** |

6. Transfer 1 mL Exonuclease Treatment Solution to the tube of BAGs; pipette to mix.
7. Incubate at 37°C with rotation for 1.5 hours.
8. Centrifuge at 400 rcf for 1.5 minutes. Discard supernatant.
9. Stop the reaction with the following washes:
    a. 1 mL STOP-25 twice
    b. 1 mL STOP-10 once.
10. Continue to NlaIII cutting or resuspend in 1 mL STOP-10 and store at 4°C.

--------------------Safe Stopping Point: Store BAGs in STOP-10 at 4°C overnight--------------------

#### NlaIII cutting

1. If BAGs are stored in STOP-10, discard the supernatant.
2. Wash once with STOP-1.
3. Wash once with 1× CutSmart buffer (900 µL $H_2O$ + 100 µL 10× CutSmart buffer). Remove supernatant.
4. Prepare the following NlaIII solution 1 and vortex.

| **NlaIII Solution 1** | **Volume (µL)** |
|---|---|
| $H_2O$ | 850 |
| 10× CutSmart buffer | 100 |
| NlaIII enzyme | 50 |
| ***Total*** | ***1 mL*** |

5. Transfer NlaIII Solution 1 to the tube of BAGs; pipette to mix.
6. Incubate at 37°C with rotation for 1.5 hours.
7. Centrifuge at 400 rcf for 1.5 minutes. Discard supernatant.
8. Prepare the following NlaIII solution 2 and vortex.

| NlaIII Solution 2 | Volume (µL) |
|---|---|
| $H_2O$ | 880 |
| CutSmart buffer | 100 |
| NlaIII enzyme | 20 |
| ***Total*** | ***1 mL*** |

9. Transfer NlaIII Solution 2 to the tube of BAGs; pipette to mix.
10. Incubate at 37°C with rotation for 1 hour.
11. Centrifuge at 400 rcf for 1.5 minutes. Discard supernatant.
12. Stop the reaction with the following washes:
    a. 1 mL STOP-25 twice
    b. 1 mL STOP-10 once.
13. Continue to First split-pool or resuspend in 1 mL STOP-10 and store at 4°C.

-------------------Safe Stopping Point: Store BAGs in STOP-10 at 4°C overnight-------------------

#### First Split-Pool

1. Preparation:
    a. If the BAGs are stored in STOP-10, discard the supernatant.
    b. Wash twice using 1 mL HBW buffer. Discard the supernatant.
2. Prepare the following Hybridization Master Mix. Pipet to mix.

| Hybridization Master Mix | Volume (µL) |
|---|---|
| BAGs in HBW | 330 |
| 2× Quick Ligase Buffer | 550 |
| dNTP | 110 |
| ***Total*** | ***990 µL*** |

    a. Quickly add 120 µL into each tube of one PCR strip. Use multipipette to transfer 9 µL Hybridization Master Mix to each well of a 96-well PCR plate.
    b. Add 2 µL 10 µM "Adapter1-BC1-vt1-X" primers into each well. Each well should have a unique $1^{st}$-split well-specific primer. Use adhesive plate seals to seal the plate. Briefly spin down, and invert the plate.
    c. Leave the plate inverted at room temperature for 5 minutes.
    d. Incubate at 50°C for 5 minutes in the thermal cycler with heated lid.
    e. Transfer plates directly to a cold PCR plate rack and move into a 4°C fridge.
    f. Keep the plate inverted at 4°C for 10 minutes, then rotate at 4°C for another 10 minutes. Proceed to extension.

3. On ice, prepare the following Extension Master Mix; vortex to mix.

| **Extension Master Mix** | **Volume (µL)** |
|---|---|
| $H_2O$ | 412.5 |
| 2× Quick Ligase Buffer | 550 |
| Klenow 3-5 exo minus | 82.5 |
| Quick Ligase enzyme | 55 |
| ***Total*** | ***1100 µL*** |

   a. Transfer 135 µL Extension Master Mix to each tube of an 8-tube PCR strip. Transfer the plate with BAGs to a cold rack, and use a multipipette to add 10 µL Extension Master Mix to each well of the BAG plate. Pipette up and down 10 times.
   b. Move the plate back into 4°C fridge with rotation for 10 minutes.
   c. Transfer plates to thermocycler at 10°C for 10 minutes.
   d. Incubate at room temperature with rotation for 20 minutes.
   e. Incubate at 37°C in the Isotemp incubator with rotation for 20 minutes.
4. Stop the reaction with the following:
   a. Add 100 µL STOP-25 to each well of the plate. Wait for 5 minutes. Centrifuge the BAG plate at 900 rcf for 2 minutes.
   b. Without disturbing the BAG pellet, remove 85 µL solution from the top of each well.
   a. Prefill a solution basin with 5 mL STOP-25 buffer. Transfer the remaining BAG solutions from each well of the plate to the solution basin (36 µL remaining volume per well).
   b. Transfer BAG solution from the solution basin to a 15 mL conical tube. Centrifuge at 300 rcf for 5 minutes. Discard supernatant.
   c. Add 1 mL STOP-10, pipette up and down, and transfer BAG solution to a 1.5 mL tube. Centrifuge 400 rcf for 1.5 minutes. Discard supernatant. Resuspend in 1 mL STOP-10 buffer.

--------------------Safe Stopping Point: Store BAGs in STOP-10 at 4°C overnight--------------------

#### Second Split-Pool

1. Preparation:
   a. If BAGs are stored in STOP-10 overnight, discard the supernatant.
   b. Wash the BAGs using 1 mL HBW buffer. Centrifuge 400 rcf for 1.5 minutes. Discard supernatant. Repeat for a second HBW wash.
2. Denaturation
   a. Prepare fresh Denaturation Solution using the following recipe:

      9.7 mL $H_2O$ + 150 µL 10 M NaOH + 150 µL 30% (wt/wt) Brij-35

   b. Add 1 mL Denaturation Solution to BAGs. Pipette up and down 5 times. Incubate for 10 minutes at room temperature with rotation. Centrifuge at 400 rcf for 2 minutes at room temperature. Discard supernatant.
   c. Wash the BAGs two more times using the following protocol:
      Add 1 mL Denaturation Solution to BAGs. Pipet up and down 5 times. Incubate for 3 minutes at room temperature with rotation. Centrifuge at 400 rcf for 2 minutes at room temperature. Discard supernatant.
3. Neutralization
   a. Prepare Neutralization Solution using the following recipe:
      7.7 mL $H_2O$ + 1 mL 1 M Tris (pH 8.0) + 1 mL 1 M NaCl + 200 µL 0.5 M EDTA + 100 µL 10% (vol/vol) Tween-20
   b. Wash BAGs twice with Neutralization Buffer using the following protocol:
      Add 1 mL Neutralization Solution to BAGs. Pipette up and down 5 times. Incubate for 3 minutes at room temperature with rotation. Centrifuge at 400 rcf for 2 minutes at room temperature. Discard supernatant.
   c. Wash BAGs once with HBW buffer
      Add 1 mL HBW to BAGs. Pipette up and down 5 times. Centrifuge at 400 rcf for 1.5 minutes. Discard supernatant.
   d. Resuspend BAGs in HBW (400 µL in total)
4. Preset thermal cycler to 75°C and Isotemp incubator to 57°C

5. Prepare the following Isothermal Amplification Buffer (IAB) Mixture in a 2mL tube; vortex to mix:

| **IAB Mixture** | **Volume (µL)** for one 96-well plate | *Volume (µL) per well* |
|---|---|---|
| $H_2O$ | 1300 | |
| 10× IABuffer | 210 | *2.1* |
| 10 mM dNTP | 150 | *1.5* |
| BAGs in HBW | 400 | *4* |
| ***Total:*** | ***2060 µL*** | ***20 µL*** |

   a. Transfer 250 µL IAB Mixture to each tube of an 8-tube PCR strip. Quickly transfer 20 µL to each well of a 96-well plate using a multipipette.
   b. Add 2 µL of the 96 different 10 µM "Adapter2-BC2-vt2-Adapter1-Y" (Y = 1,2,…,96) primers into each well. Each well should have a unique 2$^{nd}$-split primer.
   c. Invert plate for 2 minutes. Then incubate the plate in a thermal cycler at 75°C for 5 minutes.
   d. Incubate plate at 57°C in the incubator with rotation for 20 minutes.

6. Prepare the following BST Master Mix on ice; pipette to mix:

| **BST Master Mix** | **Volume (µL)** for one 96-well plate | *Volume (µL) per well* |
|---|---|---|
| $H_2O$ | 372 | 3.1 |
| 10× IABuffer | 48 | 0.4 |
| Bst2.0 | 60 | 0.5 |
| ***Total:*** | ***480 µL*** | ***4 µL*** |

   a. Transfer 60 µL BST Master Mix to each tube of an 8-tube PCR strip. Transfer 4 µL to each well of a 96-well plate using a multipipette. Pipette to mix.
   b. Incubate at 57°C with rotation for 40 minutes.
7. Stop the reaction with the following washes:
   a. Add 100 µL STOP-25 to each well of the plate. Wait for at least 5 minutes. Then centrifuge plates at 900 rcf for 2 minutes at room temperature.
   b. Without disturbing the BAG pellet, discard 85 µL solution from the top of each tube.
   c. Transfer the remaining BAG solutions from the plate to a solution basin prefilled with 5 mL STOP-25 (41 µL remaining volume per well).
   d. Transfer BAG solution from the solution basin to a 15 mL conical. Centrifuge at 300 rcf for 5 minutes. Discard supernatant.
   e. Add 1 mL STOP-10, pipette up and down, and transfer BAG solution to a 1.5 mL Eppendorf tube. Centrifuge at 400 rcf for 1.5 minutes. Discard supernatant.
8. Continue to the third split-pool or resuspend BAGs in 1 mL STOP-10 and store at 4°C.

--------------------Safe Stopping Point: Store BAGs in STOP-10 at 4°C overnight-------------------

#### Third Split-Pool

1. Filter BAGs through a 150 µm pluriStrainer cell strainer to remove fused BAGs.
2. Preparation:
   Thaw 100 µM "TG primer" and 10 µM "Adapter3-BC3-Adapter2-Z" (Z = 1,2,…,96) barcode primers. Take out BAGs from 4°C, discard supernatant, and wash the BAGs twice using 1 mL HBW buffer. Discard supernatant.
3. Use microscope to count the concentration of BAGs.
   a. Suspend 1 µL from BAG layer in 4 µL $H_2O$.
   b. Spread BAG suspension in 1 µL droplets on a clean glass microscope slide.
   c. Count the number of BAGs in each droplet. Multiply by the total volume of BAG layer for approximate BAG count.
4. Create the following TG primer/BAG mix:

| **TG primer/BAG mix** | **Volume (µL)** |
|---|---|
| 100 µM "TG primer" | 10 |
| BAGs in HBW buffer | $x$ |
| $H_2O$ | $890 - x$ |
| ***Total*** | ***900 µL*** |

   a. Quickly transfer 110 µL TG primer/BAG mix to each tube of an 8-tube PCR strip. Use a multipipette to transfer 9 µL TG primer/BAG mix to each well of a 96-well plate.
   b. Add 1 µL 10 µM "Adapter3-BC3-Adapter2-Z" (Z = 1,2,…,96) primers into each well. Each well should have a unique 3$^{rd}$-split primer. Briefly spin.
   c. On ice, add 10 µL Kapa HiFi Master Mix to each well of the 96-well plate. Pipette to mix.
5. Proceed to PCR:

| |
|---|
| **Kapa Gel PCR program**<br>95°C 1 min<br>**10 cycles of:**<br>98°C 30s<br>60°C 1 min<br>72°C 3 min<br>**Then:**<br>72°C 5 min<br>4°C infinite |

6. Purify PCR product twice with AMPure XP beads using the following protocol (allow beads to warm to room temperature for 30 minutes):
   a. First Pool: Pool 10 µL from each of the eight tubes in each column into a new PCR tube. Reseal the plate and store at -20°C for future usage.

(ex. 1A-1H) into a new PCR tube, as shown.

| 1A | 2 | 3 | 4 | 5 | 6 | 7 | 8 | 9 | 10 | 11 | 12 |
|---|---|---|---|---|---|---|---|---|---|---|---|
| B | | | | | | | | | | | |
| C | | | | | | | | | | | |
| D | | | | | | | | | | | |
| E | | | | | | | | | | | |
| F | | | | | | | | | | | |
| G | | | | | | | | | | | |
| H | | | | | | | | | | | |

| | | | | | | | | | | | |
|---|---|---|---|---|---|---|---|---|---|---|---|
| 1 | 2 | 3 | 4 | 5 | 6 | 7 | 8 | 9 | 10 | 11 | 12 |

(After pool: 80 µL per tube)

b. First Purification: Purify these 12 samples with 0.8× ratio of AMPure XP beads (64 uL), and elute with 16 µL $H_2O$ using AMPure XP protocol.

c. Second Pool:

Pool 14 µl from six tubes into one tube.

| | | | | | | | | | | | |
|---|---|---|---|---|---|---|---|---|---|---|---|
| 1 | 2 | 3 | 4 | 5 | 6 | 7 | 8 | 9 | 10 | 11 | 12 |

| | |
|---|---|
| 1 | 2 |

(After pool: 84 µL each)

d. Second Purification: Purify these two samples with another 0.8× ratio (67 µL) of AMPure XP beads, and elute with 16 µL $H_2O$.

7. Combine the product from both tubes, and run a High Sensitivity DNA Chip on the Agilent Bioanalyzer.

#### Tagmentation to Make the Final Sequencing Library

1. Based on the Bioanalyzer measurement, combine 1.5 ng of 3$^{rd}$-split PCR product with $H_2O$ to a total volume of 5 µL. Add 10 µL Nextera TD buffer.
   a. On ice, add 5 µL Amplicon Tagment enzyme. Pipette to mix.
   b. Incubate sample at 55°C for 4 minutes in the thermal cycler.
   c. Quickly add 5 µL Neutralization Buffer. Pipette to mix.
   d. Incubate at room temperature for 5 minutes
2. On ice, add the following reagents in order; pipette to mix:

| | |
|---|---|
| $H_2O$ | 8 µL |
| 10 µM Nextera_N70x oligo | 1 µL |
| 10 µM “P5-Adapter3” primer | 1 µL |
| Nextera PCR mix | 15 µL |

3. Place on thermal cycler using the following program:

| |
|---|
| **Tagmentation PCR** |
| 95°C 30s |
| **10 cycles of:** |
| 95°C 10 s |
| 55°C 30 s |
| 72°C 30 s |
| **Then:** |
| 72°C 5 min |
| 4°C infinite |

4. Purify tagmentation PCR product using 0.8× ratio (40 µL) of Ampure XP Beads, and elute using 30 µL $H_2O$
5. Run Bioanalyzer to check the final concentration.
6. Sequence sample using Illumina sequencer with custom Read1 sequencing primer "Read1-Adapter3".

### 1st-split barcodes

| name | sequence | name | sequence |
|---|---|---|---|
| Adapter1-BC1-vt1-1 | ACGCAGAGTCCGCTC CTAAG NNNN CA+T+G | Adapter1-BC1-vt1-49 | ACGCAGAGTCCGCTC ACTAG NNNN CA+T+G |
| Adapter1-BC1-vt1-2 | ACGCAGAGTCCGCTC ACAAT NNNN CA+T+G | Adapter1-BC1-vt1-50 | ACGCAGAGTCCGCTC GCACG NNNN CA+T+G |
| Adapter1-BC1-vt1-3 | ACGCAGAGTCCGCTC TAATG NNNN CA+T+G | Adapter1-BC1-vt1-51 | ACGCAGAGTCCGCTC AGAAG NNNN CA+T+G |
| Adapter1-BC1-vt1-4 | ACGCAGAGTCCGCTC CTTCG NNNN CA+T+G | Adapter1-BC1-vt1-52 | ACGCAGAGTCCGCTC ACTTC NNNN CA+T+G |
| Adapter1-BC1-vt1-5 | ACGCAGAGTCCGCTC TATAG NNNN CA+T+G | Adapter1-BC1-vt1-53 | ACGCAGAGTCCGCTC GATTA NNNN CA+T+G |
| Adapter1-BC1-vt1-6 | ACGCAGAGTCCGCTC GGTCA NNNN CA+T+G | Adapter1-BC1-vt1-54 | ACGCAGAGTCCGCTC CGTCT NNNN CA+T+G |
| Adapter1-BC1-vt1-7 | ACGCAGAGTCCGCTC TGATA NNNN CA+T+G | Adapter1-BC1-vt1-55 | ACGCAGAGTCCGCTC TATTC NNNN CA+T+G |
| Adapter1-BC1-vt1-8 | ACGCAGAGTCCGCTC CGACA NNNN CA+T+G | Adapter1-BC1-vt1-56 | ACGCAGAGTCCGCTC GTAAT NNNN CA+T+G |
| Adapter1-BC1-vt1-9 | ACGCAGAGTCCGCTC TGAGT NNNN CA+T+G | Adapter1-BC1-vt1-57 | ACGCAGAGTCCGCTC ATATA NNNN CA+T+G |
| Adapter1-BC1-vt1-10 | ACGCAGAGTCCGCTC ATTCC NNNN CA+T+G | Adapter1-BC1-vt1-58 | ACGCAGAGTCCGCTC AGACT NNNN CA+T+G |
| Adapter1-BC1-vt1-11 | ACGCAGAGTCCGCTC GATCG NNNN CA+T+G | Adapter1-BC1-vt1-59 | ACGCAGAGTCCGCTC ATTAA NNNN CA+T+G |
| Adapter1-BC1-vt1-12 | ACGCAGAGTCCGCTC TCAGA NNNN CA+T+G | Adapter1-BC1-vt1-60 | ACGCAGAGTCCGCTC TCTCG NNNN CA+T+G |
| Adapter1-BC1-vt1-13 | ACGCAGAGTCCGCTC CATGG NNNN CA+T+G | Adapter1-BC1-vt1-61 | ACGCAGAGTCCGCTC CGATG NNNN CA+T+G |
| Adapter1-BC1-vt1-14 | ACGCAGAGTCCGCTC TCTGT NNNN CA+T+G | Adapter1-BC1-vt1-62 | ACGCAGAGTCCGCTC GCTAA NNNN CA+T+G |
| Adapter1-BC1-vt1-15 | ACGCAGAGTCCGCTC ACTGA NNNN CA+T+G | Adapter1-BC1-vt1-63 | ACGCAGAGTCCGCTC TGTGG NNNN CA+T+G |
| Adapter1-BC1-vt1-16 | ACGCAGAGTCCGCTC ACACA NNNN CA+T+G | Adapter1-BC1-vt1-64 | ACGCAGAGTCCGCTC GTATG NNNN CA+T+G |
| Adapter1-BC1-vt1-17 | ACGCAGAGTCCGCTC AATAT NNNN CA+T+G | Adapter1-BC1-vt1-65 | ACGCAGAGTCCGCTC ACATG NNNN CA+T+G |
| Adapter1-BC1-vt1-18 | ACGCAGAGTCCGCTC TCAAG NNNN CA+T+G | Adapter1-BC1-vt1-66 | ACGCAGAGTCCGCTC GGATT NNNN CA+T+G |
| Adapter1-BC1-vt1-19 | ACGCAGAGTCCGCTC TTAGG NNNN CA+T+G | Adapter1-BC1-vt1-67 | ACGCAGAGTCCGCTC CAATA NNNN CA+T+G |
| Adapter1-BC1-vt1-20 | ACGCAGAGTCCGCTC AATGC NNNN CA+T+G | Adapter1-BC1-vt1-68 | ACGCAGAGTCCGCTC GCAGT NNNN CA+T+G |
| Adapter1-BC1-vt1-21 | ACGCAGAGTCCGCTC AGTAC NNNN CA+T+G | Adapter1-BC1-vt1-69 | ACGCAGAGTCCGCTC TTACC NNNN CA+T+G |
| Adapter1-BC1-vt1-22 | ACGCAGAGTCCGCTC CGAAT NNNN CA+T+G | Adapter1-BC1-vt1-70 | ACGCAGAGTCCGCTC TCATC NNNN CA+T+G |
| Adapter1-BC1-vt1-23 | ACGCAGAGTCCGCTC CAACG NNNN CA+T+G | Adapter1-BC1-vt1-71 | ACGCAGAGTCCGCTC AGATC NNNN CA+T+G |
| Adapter1-BC1-vt1-24 | ACGCAGAGTCCGCTC TAACA NNNN CA+T+G | Adapter1-BC1-vt1-72 | ACGCAGAGTCCGCTC GTTAG NNNN CA+T+G |
| Adapter1-BC1-vt1-25 | ACGCAGAGTCCGCTC GAACT NNNN CA+T+G | Adapter1-BC1-vt1-73 | ACGCAGAGTCCGCTC GTTGA NNNN CA+T+G |
| Adapter1-BC1-vt1-26 | ACGCAGAGTCCGCTC ATTGG NNNN CA+T+G | Adapter1-BC1-vt1-74 | ACGCAGAGTCCGCTC CAAGT NNNN CA+T+G |
| Adapter1-BC1-vt1-27 | ACGCAGAGTCCGCTC CATCC NNNN CA+T+G | Adapter1-BC1-vt1-75 | ACGCAGAGTCCGCTC CTTGT NNNN CA+T+G |
| Adapter1-BC1-vt1-28 | ACGCAGAGTCCGCTC ATAGT NNNN CA+T+G | Adapter1-BC1-vt1-76 | ACGCAGAGTCCGCTC GAAGA NNNN CA+T+G |
| Adapter1-BC1-vt1-29 | ACGCAGAGTCCGCTC TATCT NNNN CA+T+G | Adapter1-BC1-vt1-77 | ACGCAGAGTCCGCTC TGTAA NNNN CA+T+G |
| Adapter1-BC1-vt1-30 | ACGCAGAGTCCGCTC TCTTA NNNN CA+T+G | Adapter1-BC1-vt1-78 | ACGCAGAGTCCGCTC TCACT NNNN CA+T+G |
| Adapter1-BC1-vt1-31 | ACGCAGAGTCCGCTC GTACA NNNN CA+T+G | Adapter1-BC1-vt1-79 | ACGCAGAGTCCGCTC GCAAC NNNN CA+T+G |
| Adapter1-BC1-vt1-32 | ACGCAGAGTCCGCTC GTAGC NNNN CA+T+G | Adapter1-BC1-vt1-80 | ACGCAGAGTCCGCTC ACAGC NNNN CA+T+G |
| Adapter1-BC1-vt1-33 | ACGCAGAGTCCGCTC TAAGC NNNN CA+T+G | Adapter1-BC1-vt1-81 | ACGCAGAGTCCGCTC CTACT NNNN CA+T+G |
| Adapter1-BC1-vt1-34 | ACGCAGAGTCCGCTC ATAAC NNNN CA+T+G | Adapter1-BC1-vt1-82 | ACGCAGAGTCCGCTC GAATC NNNN CA+T+G |
| Adapter1-BC1-vt1-35 | ACGCAGAGTCCGCTC CGTAG NNNN CA+T+G | Adapter1-BC1-vt1-83 | ACGCAGAGTCCGCTC AGAGA NNNN CA+T+G |
| Adapter1-BC1-vt1-36 | ACGCAGAGTCCGCTC GATAC NNNN CA+T+G | Adapter1-BC1-vt1-84 | ACGCAGAGTCCGCTC AGTTA NNNN CA+T+G |
| Adapter1-BC1-vt1-37 | ACGCAGAGTCCGCTC TGACG NNNN CA+T+G | Adapter1-BC1-vt1-85 | ACGCAGAGTCCGCTC CATAA NNNN CA+T+G |
| Adapter1-BC1-vt1-38 | ACGCAGAGTCCGCTC GATGT NNNN CA+T+G | Adapter1-BC1-vt1-86 | ACGCAGAGTCCGCTC GGTTG NNNN CA+T+G |
| Adapter1-BC1-vt1-39 | ACGCAGAGTCCGCTC AGTGT NNNN CA+T+G | Adapter1-BC1-vt1-87 | ACGCAGAGTCCGCTC GGTAT NNNN CA+T+G |
| Adapter1-BC1-vt1-40 | ACGCAGAGTCCGCTC AGTCG NNNN CA+T+G | Adapter1-BC1-vt1-88 | ACGCAGAGTCCGCTC CTAGA NNNN CA+T+G |
| Adapter1-BC1-vt1-41 | ACGCAGAGTCCGCTC AATCA NNNN CA+T+G | Adapter1-BC1-vt1-89 | ACGCAGAGTCCGCTC ATACG NNNN CA+T+G |
| Adapter1-BC1-vt1-42 | ACGCAGAGTCCGCTC ACTCT NNNN CA+T+G | Adapter1-BC1-vt1-90 | ACGCAGAGTCCGCTC CGTTC NNNN CA+T+G |
| Adapter1-BC1-vt1-43 | ACGCAGAGTCCGCTC AATTG NNNN CA+T+G | Adapter1-BC1-vt1-91 | ACGCAGAGTCCGCTC TGAAC NNNN CA+T+G |
| Adapter1-BC1-vt1-44 | ACGCAGAGTCCGCTC CTATC NNNN CA+T+G | Adapter1-BC1-vt1-92 | ACGCAGAGTCCGCTC GTTCT NNNN CA+T+G |
| Adapter1-BC1-vt1-45 | ACGCAGAGTCCGCTC CGTGA NNNN CA+T+G | Adapter1-BC1-vt1-93 | ACGCAGAGTCCGCTC CTTAC NNNN CA+T+G |
| Adapter1-BC1-vt1-46 | ACGCAGAGTCCGCTC TCTAC NNNN CA+T+G | Adapter1-BC1-vt1-94 | ACGCAGAGTCCGCTC TTATT NNNN CA+T+G |
| Adapter1-BC1-vt1-47 | ACGCAGAGTCCGCTC TGTCC NNNN CA+T+G | Adapter1-BC1-vt1-95 | ACGCAGAGTCCGCTC TATGA NNNN CA+T+G |
| Adapter1-BC1-vt1-48 | ACGCAGAGTCCGCTC GCATA NNNN CA+T+G | Adapter1-BC1-vt1-96 | ACGCAGAGTCCGCTC CTAAG NNNN CA+T+G |

#### 2nd-split barcodes

| name | sequence |
|---|---|
| Adapter2-BC2-vt2-Adapter1-1 | GACCGACTCGCATTACCCTAT AATTTC NNNN ACGCAGAGTCCGCTC/3InvdT/ |
| Adapter2-BC2-vt2-Adapter1-2 | GACCGACTCGCATTACCCTAT ATTGGT NNNN ACGCAGAGTCCGCTC/3InvdT/ |
| Adapter2-BC2-vt2-Adapter1-3 | GACCGACTCGCATTACCCTAT ACCTGC NNNN ACGCAGAGTCCGCTC/3InvdT/ |
| Adapter2-BC2-vt2-Adapter1-4 | GACCGACTCGCATTACCCTAT TGAACA NNNN ACGCAGAGTCCGCTC/3InvdT/ |
| Adapter2-BC2-vt2-Adapter1-5 | GACCGACTCGCATTACCCTAT TTCCCA NNNN ACGCAGAGTCCGCTC/3InvdT/ |
| Adapter2-BC2-vt2-Adapter1-6 | GACCGACTCGCATTACCCTAT TTTAGT NNNN ACGCAGAGTCCGCTC/3InvdT/ |
| Adapter2-BC2-vt2-Adapter1-7 | GACCGACTCGCATTACCCTAT AACTTT NNNN ACGCAGAGTCCGCTC/3InvdT/ |
| Adapter2-BC2-vt2-Adapter1-8 | GACCGACTCGCATTACCCTAT GAACAA NNNN ACGCAGAGTCCGCTC/3InvdT/ |
| Adapter2-BC2-vt2-Adapter1-9 | GACCGACTCGCATTACCCTAT TATGGC NNNN ACGCAGAGTCCGCTC/3InvdT/ |
| Adapter2-BC2-vt2-Adapter1-10 | GACCGACTCGCATTACCCTAT AACCTG NNNN ACGCAGAGTCCGCTC/3InvdT/ |
| Adapter2-BC2-vt2-Adapter1-11 | GACCGACTCGCATTACCCTAT ACTGGC NNNN ACGCAGAGTCCGCTC/3InvdT/ |
| Adapter2-BC2-vt2-Adapter1-12 | GACCGACTCGCATTACCCTAT GTTTTA NNNN ACGCAGAGTCCGCTC/3InvdT/ |
| Adapter2-BC2-vt2-Adapter1-13 | GACCGACTCGCATTACCCTAT GAACCC NNNN ACGCAGAGTCCGCTC/3InvdT/ |
| Adapter2-BC2-vt2-Adapter1-14 | GACCGACTCGCATTACCCTAT AAAGAT NNNN ACGCAGAGTCCGCTC/3InvdT/ |
| Adapter2-BC2-vt2-Adapter1-15 | GACCGACTCGCATTACCCTAT TCTTCA NNNN ACGCAGAGTCCGCTC/3InvdT/ |
| Adapter2-BC2-vt2-Adapter1-16 | GACCGACTCGCATTACCCTAT GCCTCT NNNN ACGCAGAGTCCGCTC/3InvdT/ |
| Adapter2-BC2-vt2-Adapter1-17 | GACCGACTCGCATTACCCTAT AGCGTT NNNN ACGCAGAGTCCGCTC/3InvdT/ |
| Adapter2-BC2-vt2-Adapter1-18 | GACCGACTCGCATTACCCTAT TACTCC NNNN ACGCAGAGTCCGCTC/3InvdT/ |
| Adapter2-BC2-vt2-Adapter1-19 | GACCGACTCGCATTACCCTAT AAGCGT NNNN ACGCAGAGTCCGCTC/3InvdT/ |
| Adapter2-BC2-vt2-Adapter1-20 | GACCGACTCGCATTACCCTAT AGAACC NNNN ACGCAGAGTCCGCTC/3InvdT/ |
| Adapter2-BC2-vt2-Adapter1-21 | GACCGACTCGCATTACCCTAT TATTCT NNNN ACGCAGAGTCCGCTC/3InvdT/ |
| Adapter2-BC2-vt2-Adapter1-22 | GACCGACTCGCATTACCCTAT ATGGAT NNNN ACGCAGAGTCCGCTC/3InvdT/ |
| Adapter2-BC2-vt2-Adapter1-23 | GACCGACTCGCATTACCCTAT GACCAT NNNN ACGCAGAGTCCGCTC/3InvdT/ |
| Adapter2-BC2-vt2-Adapter1-24 | GACCGACTCGCATTACCCTAT AACAAG NNNN ACGCAGAGTCCGCTC/3InvdT/ |
| Adapter2-BC2-vt2-Adapter1-25 | GACCGACTCGCATTACCCTAT TAGCCT NNNN ACGCAGAGTCCGCTC/3InvdT/ |
| Adapter2-BC2-vt2-Adapter1-26 | GACCGACTCGCATTACCCTAT TTGCTG NNNN ACGCAGAGTCCGCTC/3InvdT/ |
| Adapter2-BC2-vt2-Adapter1-27 | GACCGACTCGCATTACCCTAT AGAGCG NNNN ACGCAGAGTCCGCTC/3InvdT/ |
| Adapter2-BC2-vt2-Adapter1-28 | GACCGACTCGCATTACCCTAT CTTCCA NNNN ACGCAGAGTCCGCTC/3InvdT/ |
| Adapter2-BC2-vt2-Adapter1-29 | GACCGACTCGCATTACCCTAT AAGATT NNNN ACGCAGAGTCCGCTC/3InvdT/ |
| Adapter2-BC2-vt2-Adapter1-30 | GACCGACTCGCATTACCCTAT GGGTTT NNNN ACGCAGAGTCCGCTC/3InvdT/ |
| Adapter2-BC2-vt2-Adapter1-31 | GACCGACTCGCATTACCCTAT CTTGGC NNNN ACGCAGAGTCCGCTC/3InvdT/ |
| Adapter2-BC2-vt2-Adapter1-32 | GACCGACTCGCATTACCCTAT GACTGG NNNN ACGCAGAGTCCGCTC/3InvdT/ |
| Adapter2-BC2-vt2-Adapter1-33 | GACCGACTCGCATTACCCTAT GTTACT NNNN ACGCAGAGTCCGCTC/3InvdT/ |
| Adapter2-BC2-vt2-Adapter1-34 | GACCGACTCGCATTACCCTAT GTGCGT NNNN ACGCAGAGTCCGCTC/3InvdT/ |
| Adapter2-BC2-vt2-Adapter1-35 | GACCGACTCGCATTACCCTAT GATAGA NNNN ACGCAGAGTCCGCTC/3InvdT/ |
| Adapter2-BC2-vt2-Adapter1-36 | GACCGACTCGCATTACCCTAT GTTAGC NNNN ACGCAGAGTCCGCTC/3InvdT/ |
| Adapter2-BC2-vt2-Adapter1-37 | GACCGACTCGCATTACCCTAT GATGCA NNNN ACGCAGAGTCCGCTC/3InvdT/ |
| Adapter2-BC2-vt2-Adapter1-38 | GACCGACTCGCATTACCCTAT AGTGAG NNNN ACGCAGAGTCCGCTC/3InvdT/ |
| Adapter2-BC2-vt2-Adapter1-39 | GACCGACTCGCATTACCCTAT TAATTT NNNN ACGCAGAGTCCGCTC/3InvdT/ |
| Adapter2-BC2-vt2-Adapter1-40 | GACCGACTCGCATTACCCTAT AAATCC NNNN ACGCAGAGTCCGCTC/3InvdT/ |
| Adapter2-BC2-vt2-Adapter1-41 | GACCGACTCGCATTACCCTAT ATACCG NNNN ACGCAGAGTCCGCTC/3InvdT/ |
| Adapter2-BC2-vt2-Adapter1-42 | GACCGACTCGCATTACCCTAT GTTCCC NNNN ACGCAGAGTCCGCTC/3InvdT/ |
| Adapter2-BC2-vt2-Adapter1-43 | GACCGACTCGCATTACCCTAT ATGAGG NNNN ACGCAGAGTCCGCTC/3InvdT/ |
| Adapter2-BC2-vt2-Adapter1-44 | GACCGACTCGCATTACCCTAT AACCGA NNNN ACGCAGAGTCCGCTC/3InvdT/ |
| Adapter2-BC2-vt2-Adapter1-45 | GACCGACTCGCATTACCCTAT TTCGGC NNNN ACGCAGAGTCCGCTC/3InvdT/ |
| Adapter2-BC2-vt2-Adapter1-46 | GACCGACTCGCATTACCCTAT AGTCAC NNNN ACGCAGAGTCCGCTC/3InvdT/ |
| Adapter2-BC2-vt2-Adapter1-47 | GACCGACTCGCATTACCCTAT TACCGT NNNN ACGCAGAGTCCGCTC/3InvdT/ |
| Adapter2-BC2-vt2-Adapter1-48 | GACCGACTCGCATTACCCTAT ATTTAC NNNN ACGCAGAGTCCGCTC/3InvdT/ |
| Adapter2-BC2-vt2-Adapter1-49 | GACCGACTCGCATTACCCTAT TAAGCT NNNN ACGCAGAGTCCGCTC/3InvdT/ |
| Adapter2-BC2-vt2-Adapter1-50 | GACCGACTCGCATTACCCTAT GAAATC NNNN ACGCAGAGTCCGCTC/3InvdT/ |
| Adapter2-BC2-vt2-Adapter1-51 | GACCGACTCGCATTACCCTAT AGTTTG NNNN ACGCAGAGTCCGCTC/3InvdT/ |
| Adapter2-BC2-vt2-Adapter1-52 | GACCGACTCGCATTACCCTAT AGAATT NNNN ACGCAGAGTCCGCTC/3InvdT/ |
| Adapter2-BC2-vt2-Adapter1-53 | GACCGACTCGCATTACCCTAT TTAAGG NNNN ACGCAGAGTCCGCTC/3InvdT/ |
| Adapter2-BC2-vt2-Adapter1-54 | GACCGACTCGCATTACCCTAT TTCCAG NNNN ACGCAGAGTCCGCTC/3InvdT/ |
| Adapter2-BC2-vt2-Adapter1-55 | GACCGACTCGCATTACCCTAT CCTGCA NNNN ACGCAGAGTCCGCTC/3InvdT/ |
| Adapter2-BC2-vt2-Adapter1-56 | GACCGACTCGCATTACCCTAT AGCCTC NNNN ACGCAGAGTCCGCTC/3InvdT/ |
| Adapter2-BC2-vt2-Adapter1-57 | GACCGACTCGCATTACCCTAT AAAATG NNNN ACGCAGAGTCCGCTC/3InvdT/ |
| Adapter2-BC2-vt2-Adapter1-58 | GACCGACTCGCATTACCCTAT TGTTAA NNNN ACGCAGAGTCCGCTC/3InvdT/ |
| Adapter2-BC2-vt2-Adapter1-59 | GACCGACTCGCATTACCCTAT GACGAA NNNN ACGCAGAGTCCGCTC/3InvdT/ |
| Adapter2-BC2-vt2-Adapter1-60 | GACCGACTCGCATTACCCTAT TTTGGA NNNN ACGCAGAGTCCGCTC/3InvdT/ |
| Adapter2-BC2-vt2-Adapter1-61 | GACCGACTCGCATTACCCTAT ATTCCT NNNN ACGCAGAGTCCGCTC/3InvdT/ |
| Adapter2-BC2-vt2-Adapter1-62 | GACCGACTCGCATTACCCTAT TAAACC NNNN ACGCAGAGTCCGCTC/3InvdT/ |
| Adapter2-BC2-vt2-Adapter1-63 | GACCGACTCGCATTACCCTAT ATGCAC NNNN ACGCAGAGTCCGCTC/3InvdT/ |
| Adapter2-BC2-vt2-Adapter1-64 | GACCGACTCGCATTACCCTAT GACTAC NNNN ACGCAGAGTCCGCTC/3InvdT/ |
| Adapter2-BC2-vt2-Adapter1-65 | GACCGACTCGCATTACCCTAT TGGCGA NNNN ACGCAGAGTCCGCTC/3InvdT/ |
| Adapter2-BC2-vt2-Adapter1-66 | GACCGACTCGCATTACCCTAT AAAGTA NNNN ACGCAGAGTCCGCTC/3InvdT/ |
| Adapter2-BC2-vt2-Adapter1-67 | GACCGACTCGCATTACCCTAT TGATGA NNNN ACGCAGAGTCCGCTC/3InvdT/ |
| Adapter2-BC2-vt2-Adapter1-68 | GACCGACTCGCATTACCCTAT TTGACT NNNN ACGCAGAGTCCGCTC/3InvdT/ |
| Adapter2-BC2-vt2-Adapter1-69 | GACCGACTCGCATTACCCTAT TTTACG NNNN ACGCAGAGTCCGCTC/3InvdT/ |
| Adapter2-BC2-vt2-Adapter1-70 | GACCGACTCGCATTACCCTAT GGTGAC NNNN ACGCAGAGTCCGCTC/3InvdT/ |
| Adapter2-BC2-vt2-Adapter1-71 | GACCGACTCGCATTACCCTAT GAAGCG NNNN ACGCAGAGTCCGCTC/3InvdT/ |
| Adapter2-BC2-vt2-Adapter1-72 | GACCGACTCGCATTACCCTAT TAACCA NNNN ACGCAGAGTCCGCTC/3InvdT/ |
| Adapter2-BC2-vt2-Adapter1-73 | GACCGACTCGCATTACCCTAT TTGGCC NNNN ACGCAGAGTCCGCTC/3InvdT/ |
| Adapter2-BC2-vt2-Adapter1-74 | GACCGACTCGCATTACCCTAT ATCGAG NNNN ACGCAGAGTCCGCTC/3InvdT/ |
| Adapter2-BC2-vt2-Adapter1-75 | GACCGACTCGCATTACCCTAT GTCTTT NNNN ACGCAGAGTCCGCTC/3InvdT/ |
| Adapter2-BC2-vt2-Adapter1-76 | GACCGACTCGCATTACCCTAT TTGAGC NNNN ACGCAGAGTCCGCTC/3InvdT/ |
| Adapter2-BC2-vt2-Adapter1-77 | GACCGACTCGCATTACCCTAT AAGGCC NNNN ACGCAGAGTCCGCTC/3InvdT/ |
| Adapter2-BC2-vt2-Adapter1-78 | GACCGACTCGCATTACCCTAT AAAGGC NNNN ACGCAGAGTCCGCTC/3InvdT/ |
| Adapter2-BC2-vt2-Adapter1-79 | GACCGACTCGCATTACCCTAT AGCTTA NNNN ACGCAGAGTCCGCTC/3InvdT/ |
| Adapter2-BC2-vt2-Adapter1-80 | GACCGACTCGCATTACCCTAT TAGGAA NNNN ACGCAGAGTCCGCTC/3InvdT/ |
| Adapter2-BC2-vt2-Adapter1-81 | GACCGACTCGCATTACCCTAT GGACGT NNNN ACGCAGAGTCCGCTC/3InvdT/ |
| Adapter2-BC2-vt2-Adapter1-82 | GACCGACTCGCATTACCCTAT TGGTGC NNNN ACGCAGAGTCCGCTC/3InvdT/ |
| Adapter2-BC2-vt2-Adapter1-83 | GACCGACTCGCATTACCCTAT ATCCGG NNNN ACGCAGAGTCCGCTC/3InvdT/ |
| Adapter2-BC2-vt2-Adapter1-84 | GACCGACTCGCATTACCCTAT GTAAAC NNNN ACGCAGAGTCCGCTC/3InvdT/ |
| Adapter2-BC2-vt2-Adapter1-85 | GACCGACTCGCATTACCCTAT TGAGGT NNNN ACGCAGAGTCCGCTC/3InvdT/ |
| Adapter2-BC2-vt2-Adapter1-86 | GACCGACTCGCATTACCCTAT ACATGA NNNN ACGCAGAGTCCGCTC/3InvdT/ |
| Adapter2-BC2-vt2-Adapter1-87 | GACCGACTCGCATTACCCTAT ATAGCA NNNN ACGCAGAGTCCGCTC/3InvdT/ |
| Adapter2-BC2-vt2-Adapter1-88 | GACCGACTCGCATTACCCTAT ATTCAG NNNN ACGCAGAGTCCGCTC/3InvdT/ |
| Adapter2-BC2-vt2-Adapter1-89 | GACCGACTCGCATTACCCTAT GAGGTA NNNN ACGCAGAGTCCGCTC/3InvdT/ |
| Adapter2-BC2-vt2-Adapter1-90 | GACCGACTCGCATTACCCTAT GTGCAA NNNN ACGCAGAGTCCGCTC/3InvdT/ |
| Adapter2-BC2-vt2-Adapter1-91 | GACCGACTCGCATTACCCTAT AATACT NNNN ACGCAGAGTCCGCTC/3InvdT/ |
| Adapter2-BC2-vt2-Adapter1-92 | GACCGACTCGCATTACCCTAT TGCAGG NNNN ACGCAGAGTCCGCTC/3InvdT/ |
| Adapter2-BC2-vt2-Adapter1-93 | GACCGACTCGCATTACCCTAT CGACTT NNNN ACGCAGAGTCCGCTC/3InvdT/ |
| Adapter2-BC2-vt2-Adapter1-94 | GACCGACTCGCATTACCCTAT GGCTAT NNNN ACGCAGAGTCCGCTC/3InvdT/ |
| Adapter2-BC2-vt2-Adapter1-95 | GACCGACTCGCATTACCCTAT AAACTT NNNN ACGCAGAGTCCGCTC/3InvdT/ |
| Adapter2-BC2-vt2-Adapter1-96 | GACCGACTCGCATTACCCTAT CTGTTT NNNN ACGCAGAGTCCGCTC/3InvdT/ |

### 3rd-split barcodes

| name | sequence |
|---|---|
| Adapter3-BC3-Adapter2-1 | AAGGCTCTTGACACGAAGGATAG TACCGTNN GACCGACTCGCATTACCCT*A*T |
| Adapter3-BC3-Adapter2-2 | AAGGCTCTTGACACGAAGGATAG CCATTTNN GACCGACTCGCATTACCCT*A*T |
| Adapter3-BC3-Adapter2-3 | AAGGCTCTTGACACGAAGGATAG TCTTTTNN GACCGACTCGCATTACCCT*A*T |
| Adapter3-BC3-Adapter2-4 | AAGGCTCTTGACACGAAGGATAG AAACATNN GACCGACTCGCATTACCCT*A*T |
| Adapter3-BC3-Adapter2-5 | AAGGCTCTTGACACGAAGGATAG GTTTGGNN GACCGACTCGCATTACCCT*A*T |
| Adapter3-BC3-Adapter2-6 | AAGGCTCTTGACACGAAGGATAG CAAACTNN GACCGACTCGCATTACCCT*A*T |
| Adapter3-BC3-Adapter2-7 | AAGGCTCTTGACACGAAGGATAG ACCTATNN GACCGACTCGCATTACCCT*A*T |
| Adapter3-BC3-Adapter2-8 | AAGGCTCTTGACACGAAGGATAG TCACTTNN GACCGACTCGCATTACCCT*A*T |
| Adapter3-BC3-Adapter2-9 | AAGGCTCTTGACACGAAGGATAG AGTTGTNN GACCGACTCGCATTACCCT*A*T |
| Adapter3-BC3-Adapter2-10 | AAGGCTCTTGACACGAAGGATAG CTTGGANN GACCGACTCGCATTACCCT*A*T |
| Adapter3-BC3-Adapter2-11 | AAGGCTCTTGACACGAAGGATAG CTGGATNN GACCGACTCGCATTACCCT*A*T |
| Adapter3-BC3-Adapter2-12 | AAGGCTCTTGACACGAAGGATAG TTCACGNN GACCGACTCGCATTACCCT*A*T |
| Adapter3-BC3-Adapter2-13 | AAGGCTCTTGACACGAAGGATAG CTCTATNN GACCGACTCGCATTACCCT*A*T |
| Adapter3-BC3-Adapter2-14 | AAGGCTCTTGACACGAAGGATAG AGAATGNN GACCGACTCGCATTACCCT*A*T |
| Adapter3-BC3-Adapter2-15 | AAGGCTCTTGACACGAAGGATAG CATCCANN GACCGACTCGCATTACCCT*A*T |
| Adapter3-BC3-Adapter2-16 | AAGGCTCTTGACACGAAGGATAG GTTTCTNN GACCGACTCGCATTACCCT*A*T |
| Adapter3-BC3-Adapter2-17 | AAGGCTCTTGACACGAAGGATAG TCAGATNN GACCGACTCGCATTACCCT*A*T |
| Adapter3-BC3-Adapter2-18 | AAGGCTCTTGACACGAAGGATAG CAATGGNN GACCGACTCGCATTACCCT*A*T |
| Adapter3-BC3-Adapter2-19 | AAGGCTCTTGACACGAAGGATAG TGCTCANN GACCGACTCGCATTACCCT*A*T |
| Adapter3-BC3-Adapter2-20 | AAGGCTCTTGACACGAAGGATAG TCTGGANN GACCGACTCGCATTACCCT*A*T |
| Adapter3-BC3-Adapter2-21 | AAGGCTCTTGACACGAAGGATAG CTTTTGNN GACCGACTCGCATTACCCT*A*T |
| Adapter3-BC3-Adapter2-22 | AAGGCTCTTGACACGAAGGATAG ACTCAGNN GACCGACTCGCATTACCCT*A*T |
| Adapter3-BC3-Adapter2-23 | AAGGCTCTTGACACGAAGGATAG AACTAGNN GACCGACTCGCATTACCCT*A*T |
| Adapter3-BC3-Adapter2-24 | AAGGCTCTTGACACGAAGGATAG CCCTTANN GACCGACTCGCATTACCCT*A*T |
| Adapter3-BC3-Adapter2-25 | AAGGCTCTTGACACGAAGGATAG CTGATGNN GACCGACTCGCATTACCCT*A*T |
| Adapter3-BC3-Adapter2-26 | AAGGCTCTTGACACGAAGGATAG CATCAGNN GACCGACTCGCATTACCCT*A*T |
| Adapter3-BC3-Adapter2-27 | AAGGCTCTTGACACGAAGGATAG AACCTGNN GACCGACTCGCATTACCCT*A*T |
| Adapter3-BC3-Adapter2-28 | AAGGCTCTTGACACGAAGGATAG CAAATGNN GACCGACTCGCATTACCCT*A*T |
| Adapter3-BC3-Adapter2-29 | AAGGCTCTTGACACGAAGGATAG TCCCTANN GACCGACTCGCATTACCCT*A*T |
| Adapter3-BC3-Adapter2-30 | AAGGCTCTTGACACGAAGGATAG TTTATGNN GACCGACTCGCATTACCCT*A*T |
| Adapter3-BC3-Adapter2-31 | AAGGCTCTTGACACGAAGGATAG CTAAGANN GACCGACTCGCATTACCCT*A*T |
| Adapter3-BC3-Adapter2-32 | AAGGCTCTTGACACGAAGGATAG AATACANN GACCGACTCGCATTACCCT*A*T |
| Adapter3-BC3-Adapter2-33 | AAGGCTCTTGACACGAAGGATAG CTAACGNN GACCGACTCGCATTACCCT*A*T |
| Adapter3-BC3-Adapter2-34 | AAGGCTCTTGACACGAAGGATAG TTCTTANN GACCGACTCGCATTACCCT*A*T |
| Adapter3-BC3-Adapter2-35 | AAGGCTCTTGACACGAAGGATAG GATGGANN GACCGACTCGCATTACCCT*A*T |
| Adapter3-BC3-Adapter2-36 | AAGGCTCTTGACACGAAGGATAG GGGATTNN GACCGACTCGCATTACCCT*A*T |
| Adapter3-BC3-Adapter2-37 | AAGGCTCTTGACACGAAGGATAG GATATTNN GACCGACTCGCATTACCCT*A*T |
| Adapter3-BC3-Adapter2-38 | AAGGCTCTTGACACGAAGGATAG AACTTTNN GACCGACTCGCATTACCCT*A*T |
| Adapter3-BC3-Adapter2-39 | AAGGCTCTTGACACGAAGGATAG GCAACANN GACCGACTCGCATTACCCT*A*T |
| Adapter3-BC3-Adapter2-40 | AAGGCTCTTGACACGAAGGATAG AGAGAGNN GACCGACTCGCATTACCCT*A*T |
| Adapter3-BC3-Adapter2-41 | AAGGCTCTTGACACGAAGGATAG TTTGGGNN GACCGACTCGCATTACCCT*A*T |
| Adapter3-BC3-Adapter2-42 | AAGGCTCTTGACACGAAGGATAG GAGAGANN GACCGACTCGCATTACCCT*A*T |
| Adapter3-BC3-Adapter2-43 | AAGGCTCTTGACACGAAGGATAG TCCTCTNN GACCGACTCGCATTACCCT*A*T |
| Adapter3-BC3-Adapter2-44 | AAGGCTCTTGACACGAAGGATAG CCTCTANN GACCGACTCGCATTACCCT*A*T |
| Adapter3-BC3-Adapter2-45 | AAGGCTCTTGACACGAAGGATAG CTTAGGNN GACCGACTCGCATTACCCT*A*T |
| Adapter3-BC3-Adapter2-46 | AAGGCTCTTGACACGAAGGATAG TTTTATNN GACCGACTCGCATTACCCT*A*T |
| Adapter3-BC3-Adapter2-47 | AAGGCTCTTGACACGAAGGATAG TAGCTTNN GACCGACTCGCATTACCCT*A*T |
| Adapter3-BC3-Adapter2-48 | AAGGCTCTTGACACGAAGGATAG TCAACTNN GACCGACTCGCATTACCCT*A*T |

| name | sequence |
|---|---|
| Adapter3-BC3-Adapter2-49 | AAGGCTCTTGACACGAAGGATAG AGTATTNN GACCGACTCGCATTACCCT*A*T |
| Adapter3-BC3-Adapter2-50 | AAGGCTCTTGACACGAAGGATAG TGCTATNN GACCGACTCGCATTACCCT*A*T |
| Adapter3-BC3-Adapter2-51 | AAGGCTCTTGACACGAAGGATAG GTTCCANN GACCGACTCGCATTACCCT*A*T |
| Adapter3-BC3-Adapter2-52 | AAGGCTCTTGACACGAAGGATAG TGAACGNN GACCGACTCGCATTACCCT*A*T |
| Adapter3-BC3-Adapter2-53 | AAGGCTCTTGACACGAAGGATAG TGATGGNN GACCGACTCGCATTACCCT*A*T |
| Adapter3-BC3-Adapter2-54 | AAGGCTCTTGACACGAAGGATAG GTATTTNN GACCGACTCGCATTACCCT*A*T |
| Adapter3-BC3-Adapter2-55 | AAGGCTCTTGACACGAAGGATAG GTACAGNN GACCGACTCGCATTACCCT*A*T |
| Adapter3-BC3-Adapter2-56 | AAGGCTCTTGACACGAAGGATAG AAACCGNN GACCGACTCGCATTACCCT*A*T |
| Adapter3-BC3-Adapter2-57 | AAGGCTCTTGACACGAAGGATAG CATTTANN GACCGACTCGCATTACCCT*A*T |
| Adapter3-BC3-Adapter2-58 | AAGGCTCTTGACACGAAGGATAG AGGGATNN GACCGACTCGCATTACCCT*A*T |
| Adapter3-BC3-Adapter2-59 | AAGGCTCTTGACACGAAGGATAG AAATCANN GACCGACTCGCATTACCCT*A*T |
| Adapter3-BC3-Adapter2-60 | AAGGCTCTTGACACGAAGGATAG GGAACTNN GACCGACTCGCATTACCCT*A*T |
| Adapter3-BC3-Adapter2-61 | AAGGCTCTTGACACGAAGGATAG CTCATANN GACCGACTCGCATTACCCT*A*T |
| Adapter3-BC3-Adapter2-62 | AAGGCTCTTGACACGAAGGATAG TAATCGNN GACCGACTCGCATTACCCT*A*T |
| Adapter3-BC3-Adapter2-63 | AAGGCTCTTGACACGAAGGATAG TAGTTANN GACCGACTCGCATTACCCT*A*T |
| Adapter3-BC3-Adapter2-64 | AAGGCTCTTGACACGAAGGATAG AATATGNN GACCGACTCGCATTACCCT*A*T |
| Adapter3-BC3-Adapter2-65 | AAGGCTCTTGACACGAAGGATAG TAGTCTNN GACCGACTCGCATTACCCT*A*T |
| Adapter3-BC3-Adapter2-66 | AAGGCTCTTGACACGAAGGATAG CATTCTNN GACCGACTCGCATTACCCT*A*T |
| Adapter3-BC3-Adapter2-67 | AAGGCTCTTGACACGAAGGATAG TTATGANN GACCGACTCGCATTACCCT*A*T |
| Adapter3-BC3-Adapter2-68 | AAGGCTCTTGACACGAAGGATAG CCTCATNN GACCGACTCGCATTACCCT*A*T |
| Adapter3-BC3-Adapter2-69 | AAGGCTCTTGACACGAAGGATAG GACTTANN GACCGACTCGCATTACCCT*A*T |
| Adapter3-BC3-Adapter2-70 | AAGGCTCTTGACACGAAGGATAG ACAACGNN GACCGACTCGCATTACCCT*A*T |
| Adapter3-BC3-Adapter2-71 | AAGGCTCTTGACACGAAGGATAG GATTTGNN GACCGACTCGCATTACCCT*A*T |
| Adapter3-BC3-Adapter2-72 | AAGGCTCTTGACACGAAGGATAG GACCTTNN GACCGACTCGCATTACCCT*A*T |
| Adapter3-BC3-Adapter2-73 | AAGGCTCTTGACACGAAGGATAG CGAATTNN GACCGACTCGCATTACCCT*A*T |
| Adapter3-BC3-Adapter2-74 | AAGGCTCTTGACACGAAGGATAG TTATTGNN GACCGACTCGCATTACCCT*A*T |
| Adapter3-BC3-Adapter2-75 | AAGGCTCTTGACACGAAGGATAG ACTTTGNN GACCGACTCGCATTACCCT*A*T |
| Adapter3-BC3-Adapter2-76 | AAGGCTCTTGACACGAAGGATAG AACTGANN GACCGACTCGCATTACCCT*A*T |
| Adapter3-BC3-Adapter2-77 | AAGGCTCTTGACACGAAGGATAG GCTACTNN GACCGACTCGCATTACCCT*A*T |
| Adapter3-BC3-Adapter2-78 | AAGGCTCTTGACACGAAGGATAG AAATTGNN GACCGACTCGCATTACCCT*A*T |
| Adapter3-BC3-Adapter2-79 | AAGGCTCTTGACACGAAGGATAG ACCTCANN GACCGACTCGCATTACCCT*A*T |
| Adapter3-BC3-Adapter2-80 | AAGGCTCTTGACACGAAGGATAG GGATGANN GACCGACTCGCATTACCCT*A*T |
| Adapter3-BC3-Adapter2-81 | AAGGCTCTTGACACGAAGGATAG GACATGNN GACCGACTCGCATTACCCT*A*T |
| Adapter3-BC3-Adapter2-82 | AAGGCTCTTGACACGAAGGATAG AATCTTNN GACCGACTCGCATTACCCT*A*T |
| Adapter3-BC3-Adapter2-83 | AAGGCTCTTGACACGAAGGATAG TAATTTNN GACCGACTCGCATTACCCT*A*T |
| Adapter3-BC3-Adapter2-84 | AAGGCTCTTGACACGAAGGATAG TCCTGANN GACCGACTCGCATTACCCT*A*T |
| Adapter3-BC3-Adapter2-85 | AAGGCTCTTGACACGAAGGATAG GTTCTTNN GACCGACTCGCATTACCCT*A*T |
| Adapter3-BC3-Adapter2-86 | AAGGCTCTTGACACGAAGGATAG GCTTCANN GACCGACTCGCATTACCCT*A*T |
| Adapter3-BC3-Adapter2-87 | AAGGCTCTTGACACGAAGGATAG TTAGAGNN GACCGACTCGCATTACCCT*A*T |
| Adapter3-BC3-Adapter2-88 | AAGGCTCTTGACACGAAGGATAG TAACTGNN GACCGACTCGCATTACCCT*A*T |
| Adapter3-BC3-Adapter2-89 | AAGGCTCTTGACACGAAGGATAG CTGTTANN GACCGACTCGCATTACCCT*A*T |
| Adapter3-BC3-Adapter2-90 | AAGGCTCTTGACACGAAGGATAG CACCATNN GACCGACTCGCATTACCCT*A*T |
| Adapter3-BC3-Adapter2-91 | AAGGCTCTTGACACGAAGGATAG AAATGTNN GACCGACTCGCATTACCCT*A*T |
| Adapter3-BC3-Adapter2-92 | AAGGCTCTTGACACGAAGGATAG CTCTCANN GACCGACTCGCATTACCCT*A*T |
| Adapter3-BC3-Adapter2-93 | AAGGCTCTTGACACGAAGGATAG TTCCCTNN GACCGACTCGCATTACCCT*A*T |
| Adapter3-BC3-Adapter2-94 | AAGGCTCTTGACACGAAGGATAG TTCGATNN GACCGACTCGCATTACCCT*A*T |
| Adapter3-BC3-Adapter2-95 | AAGGCTCTTGACACGAAGGATAG CTATGTNN GACCGACTCGCATTACCCT*A*T |
| Adapter3-BC3-Adapter2-96 | AAGGCTCTTGACACGAAGGATAG TCATCANN GACCGACTCGCATTACCCT*A*T |

### Nextera N70x sequences

| name | sequence |
|---|---|
| Nextera_N701 | CAAGCAGAAGACGGCATACGAGAT TCGCCTTA GTCTCGTGGGCTCGG |
| Nextera_N702 | CAAGCAGAAGACGGCATACGAGAT CTAGTACG GTCTCGTGGGCTCGG |
| Nextera_N703 | CAAGCAGAAGACGGCATACGAGAT TTCTGCCT GTCTCGTGGGCTCGG |
| Nextera_N704 | CAAGCAGAAGACGGCATACGAGAT GCTCAGGA GTCTCGTGGGCTCGG |
| Nextera_N705 | CAAGCAGAAGACGGCATACGAGAT AGGAGTCC GTCTCGTGGGCTCGG |
| Nextera_N706 | CAAGCAGAAGACGGCATACGAGAT CATGCCTA GTCTCGTGGGCTCGG |
| Nextera_N707 | CAAGCAGAAGACGGCATACGAGAT GTAGAGAG GTCTCGTGGGCTCGG |
| Nextera_N708 | CAAGCAGAAGACGGCATACGAGAT CCTCTCTG GTCTCGTGGGCTCGG |
| Nextera_N709 | CAAGCAGAAGACGGCATACGAGAT AGCGTAGC GTCTCGTGGGCTCGG |
| Nextera_N710 | CAAGCAGAAGACGGCATACGAGAT CAGCCTCG GTCTCGTGGGCTCGG |
| Nextera_N711 | CAAGCAGAAGACGGCATACGAGAT TGCCTCTT GTCTCGTGGGCTCGG |
| Nextera_N712 | CAAGCAGAAGACGGCATACGAGAT TCCTCTAC GTCTCGTGGGCTCGG |
